# Single-Cell Study Designs Are Systematically Underpowered for Small-Effect Genes

**DOI:** 10.64898/2026.09.17.752063

**Authors:** Chris Crampton, Salman Fawad, Toby Clark, Hiru Dash, Bence Kövér, Garry Cotton, Donghoon Lee, Eugene Duff, Leonardo Bottolo, Paul Matthews, Nathan Skene

## Abstract

Single cell differential expression analysis enables biologists to make statistical conclusions about which genes are up or downregulated in a particular cell type, between two conditions, such as those with or without a disease. However, due to biological and technical noise, these differences are hard to detect reliably. Determining if an experiment has sufficient power is often derived from simulations or small pilot samples, if done at all. Here, we use sex-biased differential expression on 1,494 donors in three brain cell types to derive empirically grounded power estimates for a variety of experimental setups.

Our work reveals a substantial lack of power for reliably detecting the small effects in the range that many studies report, even in experiments containing 600 donors. When reducing the astrocyte data to the poor sequencing characteristics of microglia, over half the differentially expressed genes (DEGs) detected in the full set were lost, highlighting the uncertainty introduced by insufficient sequencing depth and cell counts. Despite standard thresholds of adjusted p-value with multiple testing correction, only the top quartile of significant genes were reproducible. Furthermore, from fitting a predictive model to the empirical outcomes, we find cell count and expression levels of genes to be strong determinants of power. As such, we advocate for future studies to employ cell type enrichment and deeper sequencing, especially for rarer populations like microglia, emphasise the importance of powering experiments of this type when seeking robust findings, and suggest stricter significance thresholds for future discoveries.

## Introduction

Single-cell and single-nucleus RNA-seq (sc/snRNA-seq) have become standard tools for profiling gene expression in individual cells. Simply put, gene expression is the process of transcribing DNA into an RNA transcript. RNA-seq methods break open (lyse) the cells, and use the number of transcripts present as a proxy measure for gene expression. This enables comparisons between individual cell populations within or across samples, for example between cell types or disease states. Differential expression analysis provides the statistical framework for these comparisons by identifying genes with statistically significant differences in expression between populations, commonly described by their log_2_ fold-change (LFC). The challenge comes from both real differences between cells and the many opportunities for information to be lost or distorted as a sample is processed. Gene expression detected is influenced by biological and technical factors, so even similar cells can express genes at very different levels. On top of this, some transcripts are degraded when cells are lysed, some are never captured or copied, and the sequencing process may read some transcripts many times while missing others completely. Single-cell RNA sequencing therefore provides only a partial snapshot of each cell, and the considerable variation between cells makes it difficult to draw simple conclusions about the population as a whole. Pseudobulk aggregation, where counts are summed across all cells of a given type within each replicate, has emerged as the recommended approach for reproducibility. This is due to controlling pseudoreplication, which is the inflation of sample size between highly correlated datapoints. This dramatically reduces false positives, and appropriately handles inflated zero-counts (dropout)^1–6^.

Single-cell studies typically rely on modest cohort sizes of 10-100 donors (Supp. Table 1), with variable sequencing depths. The statistical power of these studies to detect differential expression is often difficult to assess, and the experimental requirements for reliable detection remain poorly defined. Concerns about statistical power are not unique to single-cell transcriptomics. Across many areas of biology, underpowered studies have been shown to inflate effect-size estimates, increase false-positive rates and reduce reproducibility^7,8^. In neuroimaging, a 2013 meta-analysis of 730 studies estimated median statistical power at just 21%^8^, while subsequent work showed that standard cluster-based fMRI inference can produce false-positive rates of up to 70% in standard analysis settings^9^. More recently, investigation into the replicability of research in preclinical cancer biology found that the median effect size in the authors’ independent replications was 85% smaller than in the original experiments^10^. These findings underscore the need for empirical power benchmarks in contexts with high dimensional data and small cohorts, as seen in current single-cell transcriptomic experiments.

Several methods have been developed to estimate statistical power for single-cell differential expression studies^11–14^. These methods employ a variety of approaches, but all rely on assumptions of the underlying data, imputing effects on a variety of simulated experiments to determine expected detection rate. While these approaches provide a practical framework for study planning, their predictions are fundamentally limited by the uncertainty of parameter estimates obtained from their typically small pilot datasets and have undergone little formal validation against large empirical datasets. Single-cell case-control studies of brain disease typically target modest expression differences: the foundational Alzheimer’s snRNA-seq study by Mathys^15^ calls DEGs at |LFC| > 0.25, and the largest single-cell schizophrenia meta-analysis to date uses |LFC| > 0.1 as its significance threshold^16^, ∼19% and ∼7% changes respectively. Strong stimulation contrasts however, routinely produce shifts in the |LFC| > 2 regime: cytokine stimulation contrasts in single-cell data drive canonical interferon-stimulated genes to |LFC| of 4–8^17^, a 16- and 256-fold increase respectively. Larger effect sizes are easier to declare confidently with smaller sample sizes, highlighting the need for power analyses to be tailored to their experiments, with particular importance for studies seeking subtle expression changes.

Our approach splits a filtered 1,494 donor dataset into two halves, each of which are an order of magnitude greater than most other datasets of this kind. We treat the DE results of one half as gold-standard ground truth, accepting false positives as a consequence of keeping all true positives, and downsample the other into a variety of pseudo-experiments whose findings are compared with the independent half’s ground truth. Figure 1 shows a summary of the analytical framework. This process is repeated across three glial cell types, due to their relevance to neurological conditions, and three differential expression packages to mitigate method bias. We use sex-biased expression as the case-control contrast because cohorts split near 50:50, sex-biased DEGs are well-documented and reproducible across cohorts, and small-effect sex-biased DEGs are predominantly autosomal, supporting extrapolation to other comparisons. We fit machine learning models to the empirical results to show how the experimental setup and target gene characteristics, such as mean expression and dispersion, affects the probability of detection. We perform an orthogonal experiment into the effects of count-depth and cell numbers per donor by downsampling the more prevalent astrocyte population to match the characteristics of the rare microglial population, where we see over half of the DEGs previously found, to be lost. The results reveal that smaller-effect genes (|LFC| < 2) remain systematically underdetected across the observable design space, providing a case study quantifying the reproducibility crisis facing biology. To improve reproducibility in future practice, we make three recommendations: 1) consideration of the characteristics of genes of interest, 2) performing cell type enrichment for rare populations like microglia, due to the significant impact more cells per donor has on power, and 3) using more stringent thresholds for significance.

**Figure 1:**
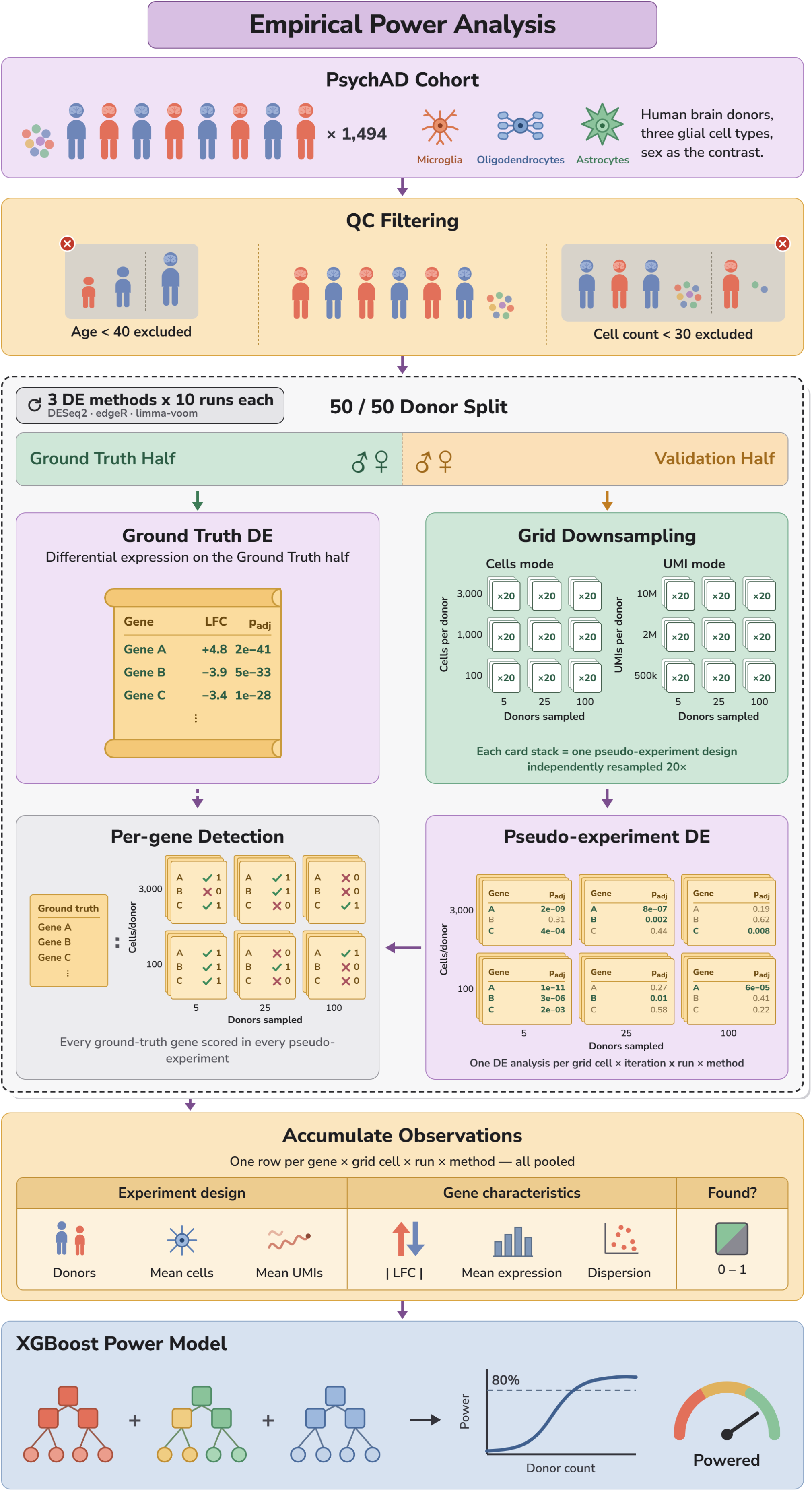
Overview of the empirical power analysis framework: the filtered PsychAD cohort is split ten times into 50/50 ground truth and validation sets, with the validation set systematically downsampled across grids in both cells-based and UMI-based modes. This is repeated for three differential expression packages. Per-gene detection outcomes between each pseudo-experiment and the corresponding ground truth are accumulated and fit to machine learning models.

**Figure 2:**
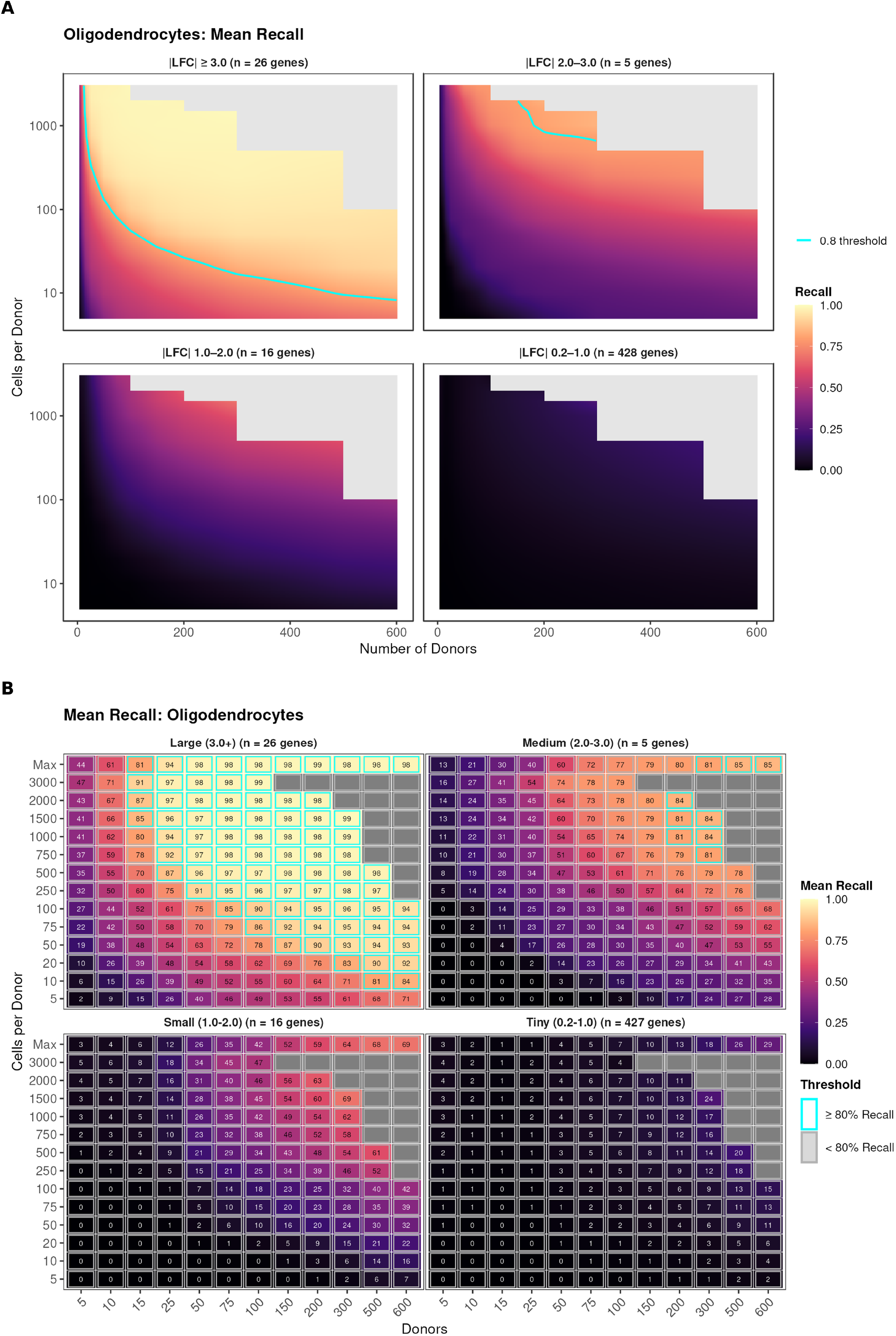
Small and tiny effect genes are systematically undetected across all pseudo-experimental setups tested. Mean recall is the mean proportion of genes that the ground truth declared were differentially expressed, that were recovered by each pseudo-experiment within a grid point, with a value of 1 indicating all genes were recovered. (A) Continuous interpolated recall surface (y-axis log-transformed) across donor count and cells per donor, faceted by |LFC| bin: tiny (0.2–1.0), small (1.0–2.0), medium (2.0–3.0) and large (>3.0). Surfaces are bilinear interpolations of the discrete outcome grid on fixed axes to allow more reliable interpretation of trends. (B) Discrete recall heatmap for the same design space; cyan borders mark configurations achieving ≥80% recall. Large-effect genes reach 80% recall at 15 donors when ≥1,500 cells per donor are available, whereas small- and tiny-effect genes remain below 80% across the entire explored space. Regions marked in grey indicate an infeasible design space due to insufficient donors with the minimum cell count required. Astrocyte and microglial recall surfaces are in Supplementary Figure 24.

## Results

### Dataset Characteristics and Filtering Rationale

To find the most significant factors affecting donor to donor variance, we applied principal component analysis on the unfiltered PsychAD dataset. Principal component 1 (PC1) showed that the largest driver was cell count across all three cell types, with donors below 30 cells showing progressively larger deviations from the mean (Supplementary Figure 1). PC2 showed a trend with donor age and clear clustering in the case of microglia for donors younger than 40. These observations motivated filtering criteria to improve stability: a minimum of 30 cells per donor and age ≥ 40 years to stabilise the age covariate with minimal loss of donors. The filtered analysis cohort comprised 1,293 astrocyte donors (mean 523 cells), 1,214 microglia donors (mean 209 cells), and 1,303 oligodendrocyte donors (mean 1,603 cells). Per-cell library sizes varied substantially: astrocytes exhibited the most UMIs (mean 9,456 UMIs per cell), followed by oligodendrocytes (5,742 UMIs) and microglia (4,426 UMIs). The difference in mean number of UMIs per donor between microglia and oligodendrocytes is ten-fold (∼0.9 million UMIs per donor for microglia versus 9.2 million for oligodendrocytes). This difference affects power for ground truth estimation, which we discuss in more detail throughout.

To characterise sex-biased differentially expressed genes, we created a set from the whole PsychAD cohort (*p*_*adj*_ < 0.05, |LFC| > 0.2), using DESeq2^18^. It comprised 408 genes in astrocytes, 236 in microglia and 425 in oligodendrocytes. The genes are predominantly autosomal (82%, 77% and 77% respectively). Allosomal genes (those on the sex chromosomes) consistently have the largest effect sizes (|LFC|). Comparing those from both groups within the 0.2 - 1 |LFC| range shows autosomal genes have median dispersion roughly three to eight fold higher (Astro 0.46 vs 0.10; Micro 0.59 vs 0.07; Oligo 0.51 vs 0.14) and median expression five to six fold lower (Astro 9.7 vs 62.4; Micro 5.1 vs 27.9; Oligo 8.1 vs 44.7). Sex-biased genes therefore offer an opportunity to examine both cases discussed in the Introduction, in parallel: subtle regulatory effects modeled by autosomal differentially expressed genes, and large, high expression effects modeled by allosomal genes.

### Empirical Power Analysis Across Experimental Designs

To empirically quantify detection power, we developed a grid downsampling framework. The filtered PsychAD cohort was split 10 times into sex-stratified ground truth (GT) and validation sets of equal size, per cell type. The validation half was systematically downsampled across a grid spanning 12 different numbers of donors (5–600) and 14 different numbers of cells per donor (5–3,000), with 20 iterations per grid point. In practice, the full grid search space was not tested owing to insufficient donors with enough cells each. This was repeated with varying UMI-counts per donor. We characterise four classes of effect size throughout the analysis: tiny (|LFC| 0.2–1), small (|LFC| 1-2), medium (|LFC| 2-3), and large (|LFC| > 3). We performed this experiment using the same samples in each case for three differential expression packages: DESeq2, edgeR, and limma-voom^19,20^. The reported results are aggregated across the methods, but method-specific results can be found in Supplementary Figures 2–5, and full per-method (DESeq2, edgeR, limma-voom) downsampling surfaces for every cell type in Supplementary Figures 6–23.

Power varied substantially with target effect size. Large-effect genes achieved 80% recall with as few as 15 donors when at least 1,500 cells each were available. At 50 donors with 500 cells, oligodendrocyte large-|LFC| recall was 87%, and even 10 cells per donor yielded 81% recall with 500 donors. Genes with moderate effect sizes only consistently crossed the recall threshold for experiments with over 750 cells per donor, and 200-300 donors. This should be interpreted with caution however as there was only an average of five genes in this set, albeit repeated 200 times per grid cell each. On average, genes with |LFC| < 2 did not exceed 80% recall across the entire experimental space, even at the maximum configuration of 600 donors with all available cells, except for with limma-voom, which showed a stricter truth set than the other two methods consistently, biasing towards higher recall. In the tiny effect bin, 75 donors were required to reach even 10% recall, regardless of cell count. This only increased to 21% with 200 donors, and never exceeded 42% across the whole space. This pattern was consistent across all three cell types tested (Supplementary Figure 24).

We find an extreme difference in recall when partitioned by adjusted p-value, despite all genes passing standard thresholds, namely Benjamini-Hochberg multiple testing correction (Figure 3, panel A). Mean expression is also shown to be a significant determinant in recall (Figure 3, panels B,C), where there is clear separation between genes with mean expression over 200 counts in the tiny-effect bin, and those below. Mean expression is calculated as the raw mean pseudobulk count for each gene across all donors in the filtered dataset. We see a similar clear separation by mean expression in the small-effect genes, however, there were 0 high expression genes in the small bin.

**Figure 3:**
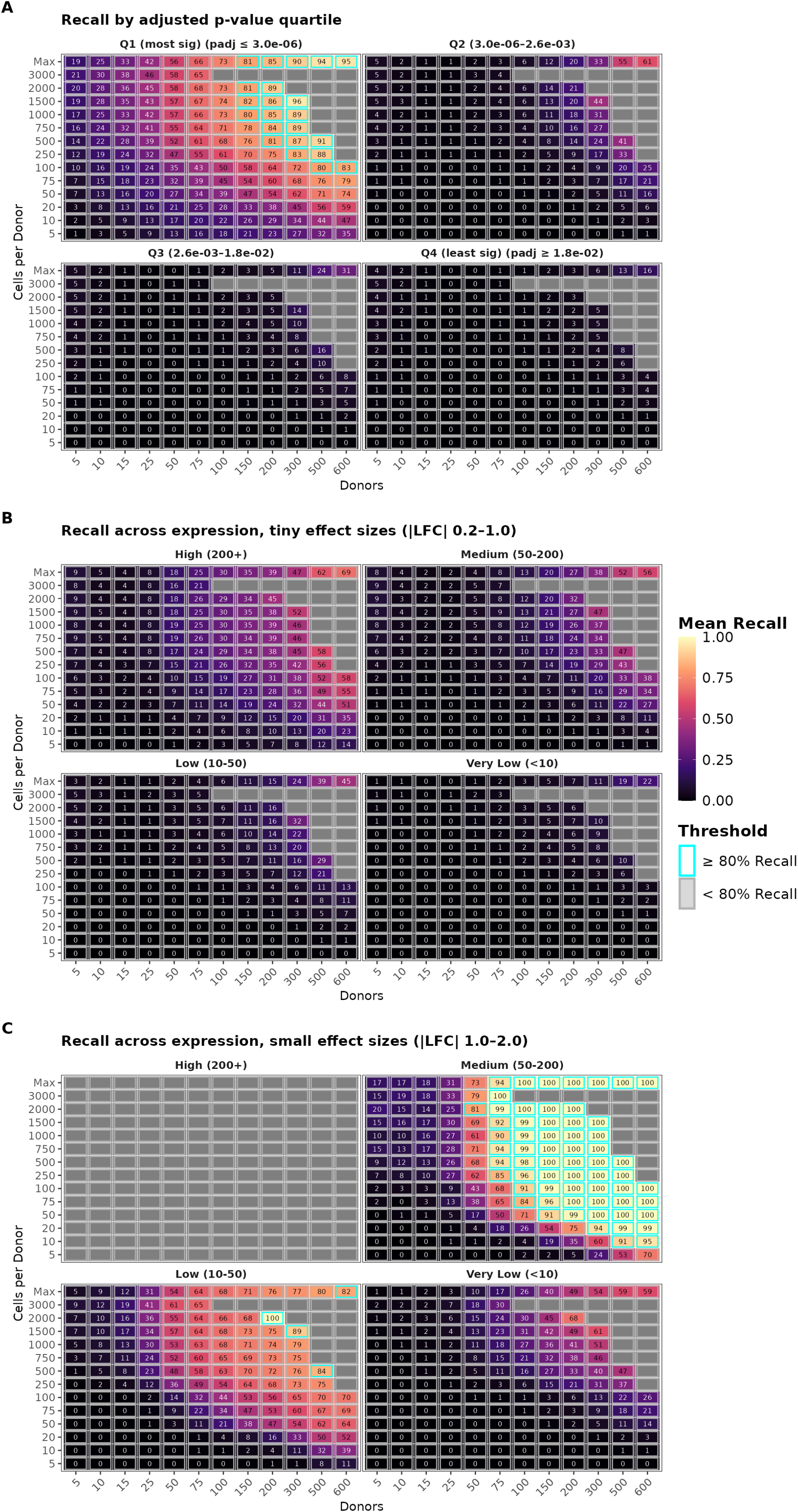
Only the most significant differentially expressed genes in the ground truth were robustly detected, despite all passing the standard threshold (*p*_*adj*_ < 0.05), and gene-expression is a strong determinant of detectability. Mean recall is the mean proportion of genes that the ground truth declared were differentially expressed, that were recovered by each pseudo-experiment within a grid point, with a value of 1 indicating all genes were recovered. Each grid shows mean recall across donor count (x-axis) and cells per donor (y-axis, log-scaled; ‘Max’ denotes all available cells), with cyan borders marking configurations reaching ≥80% recall; panels share a common recall colour scale. (A) Recall stratified into quartiles of ground-truth adjusted p-value, from the most significant genes (Q1) to the least significant (Q4). Recall degrades sharply from Q1 to Q4, showing that detectability tracks ground-truth significance. (B) Recall for tiny-effect genes (|LFC| 0.2–1.0) split into four mean-expression categories, from high (200+) to very low (<10); power collapses as expression falls even within a single effect-size bin. Mean expression is calculated as the raw mean pseudobulk count for each gene across all donors in the filtered dataset. (C) The same expression decomposition for small-effect genes (|LFC| 1.0–2.0).

Precision patterns were qualitatively consistent across cell types but depended sharply on effect-size (Figure 4, panels A,B; Supplementary Figure 25). For large-effect genes, precision exceeded 90% across nearly the entire grid in all three cell types, with breakdown confined to the 5 donor column. Medium-effect genes showed an intermediate pattern, with precision climbing above 80% only beyond 25 donors, though estimates were noisy owing to smaller truth sets (n = 5 genes/run). Precision was poor in the small and tiny effect bins, only reaching the 80% threshold at 300 donors or above, or only once in the tiny bin at 100 donors. There was poor consistency in the tiny bin in particular, indicating it to be the noisiest and likely most prone to false positives. Across the board, precision declined as cells per donor increased, the opposite of the recall trend. We propose two primary causes for this: higher cell counts bring low-expression, unstable genes into the testable universe; and the ground truth missing true DEGs which the validation half correctly found, but were therefore deemed false positives. A lack of power could be present in the ground truth sets due to the inclusion of low cell-count donors diluting the signal or increasing dispersion through dropout. This is supported by recall often being poorer with no minimum cell-count enforcement than with the 1500-3000 cell experiments.

**Figure 4:**
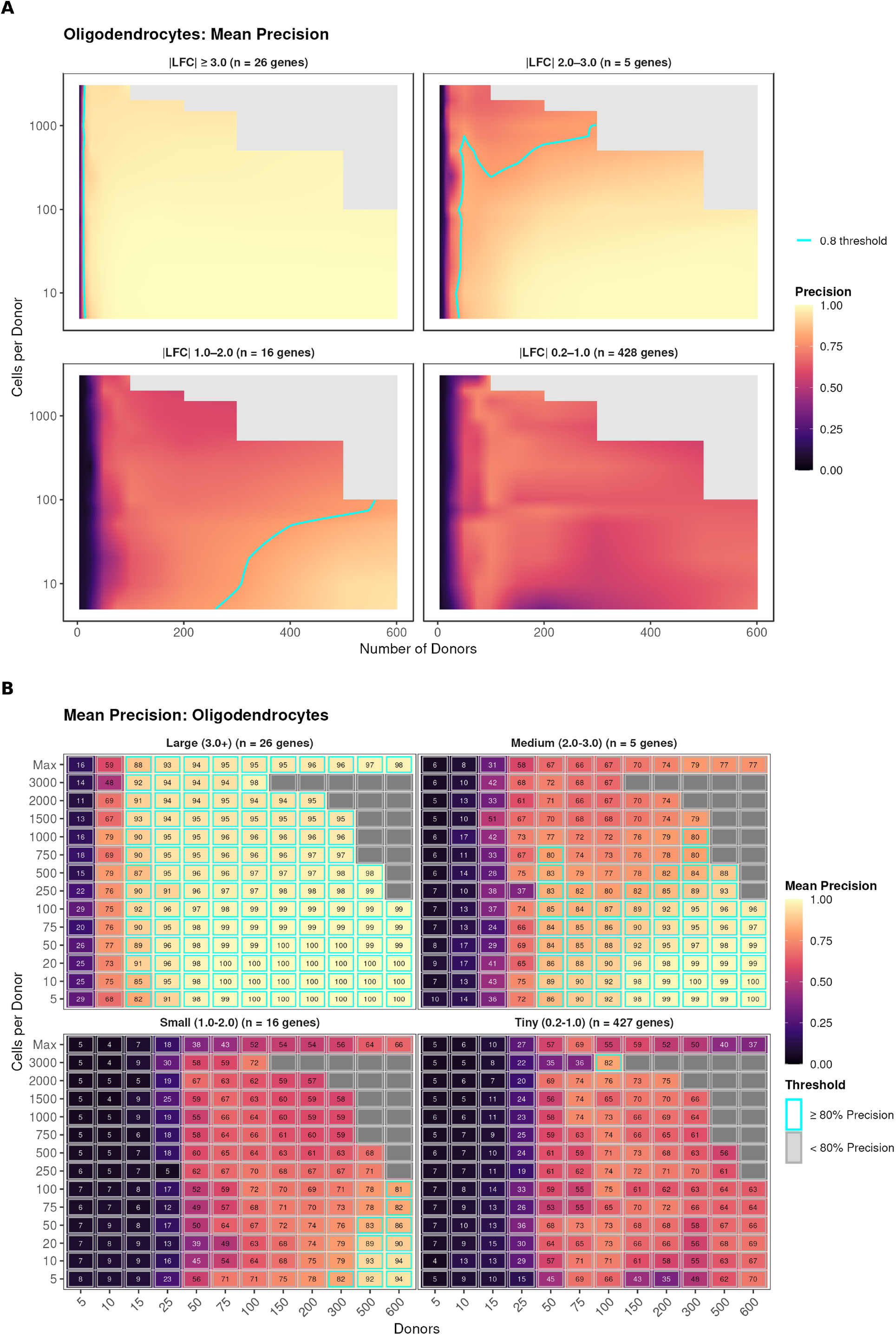
The smaller the effect size, the more likely it is to be a false positive. Mean precision is the mean proportion of genes that the pseudo-experiment declared were differentially expressed, that were agreed with by the ground truth, with a value of 1 indicating no false positives. (A) Continuous interpolated precision surface (y-axis log-transformed) across donor count and cells per donor, faceted by |LFC| bin: tiny (0.2–1.0), small (1.0–2.0), medium (2.0–3.0) and large (>3.0). Surfaces are bilinear interpolations of the discrete outcome grid on fixed axes to allow more reliable interpretation of trends. (B) Discrete precision heatmap for the same design space; cyan borders mark configurations achieving ≥80% precision. Large-effect genes reach 80% precision at 5 donors when any number of cells per donor are available, whereas small- and tiny-effect genes exhibit poor precision. Regions marked in grey indicate an infeasible design space due to insufficient donors with the minimum cell count required.

UMI-budget recall patterns were qualitatively identical to cells-based results. We performed a parallel analysis using UMI-based downsampling, where grid axes were donor count and total UMI budget per donor rather than cells per donor (Supplementary Figure 24). This provided an alternative sampling approach that would mitigate the bias in cells-based downsampling towards cell types whose cells contained more UMIs natively. In practice this showed a similar hierarchy as with cells-based downsampling, but with oligodendrocytes and astrocytes outperforming each other interchangeably. The fixed UMI budget naturally disadvantages a cell type like astrocytes, as each cell on average has double the UMIs of a microglia, making the same budget yield half the cells.

### Rare Cell Types Give more Reproducible Results, at First Glance

We observed recall to be anticorrelated with the amount of data for each cell type: microglia > astrocytes > oligodendrocytes. For example, in the tiny-effect bin for 300 donors and 100 cells each: microglia had 21% recall, astrocytes 12%, and oligodendrocytes 12% (Figure 2, panel B); Supplementary Figures 26). To probe this phenomenon, we generated three depth-equalised astrocyte sets to match the cells per donor and the UMI per cell distributions of microglia. There were four configurations: unchanged, cells downsampled, UMIs downsampled, and both downsampled. We performed DE analysis using the three packages on every ground truth set to see how the set size changed. We find that in the fully matched examples, the count of DEGs decreases by roughly 55% for both DESeq2 and edgeR compared with the full set of genes found with |LFC| ≥ 0.2 (Supplementary Table 3), with limma-voom proving numerically unstable and instead inflating the number of DEGs to over five times that of the original set (Supplementary Table 3). This suggests that the under-representation of microglia in RNA-seq datasets means many DEGs are being missed purely due to the smaller effective sample size arising from natural cell type abundance. Decreasing the number of cells had a larger impact on the number of DEGs recovered than matching the UMI distribution, indicating that prioritising maximising cell count for, a given type per donor, is preferable to increasing the number of reads. For rare cell types like microglia this would require cell sorting, as they are only a tiny fraction of the population within a tissue sample, so processing a larger portion of the tissue to increase microglial cell yield, would be disproportionately costly.

### Different Genes, Different Power Requirements

For a clearer picture of the relationship between gene characteristics, experimental setup and power, we fit the outcomes of the pseudo-experiments to an XGBoost machine learning model^21^, restricting training to only the realistic regime of experiments (cells per donor ≥ 100). XGBoost is a model that makes predictions by combining many simple decision-making steps, each new one focusing on correcting the mistakes of the previous ones. For example, the first tree might declare all genes with |LFC| > 2 as 80% detectable regardless of experimental setup, but the second tree can say if you have less than 50 donors then it drops down to 30%, and so on, each time tackling a smaller and smaller case until there are no worthwhile separations left to make. We tested 36 different hyperparameter combinations of the model to both test sensitivity and improve fit. We measure the success of a model by bin-balanced mean absolute error (MAE), quantifying the average distance between the model’s prediction and the observed power, adjusted to account for most data living at the extremes. Models partition cleanly across almost all hyperparameter configurations, preferring: no *α* weighting, larger maximum depth, smaller learning rates, and larger minimum child weight, in that order. The winning model achieves MAE = 0.165 with *α* = 0, depth 6, learning rate 0.03, mcw 10; the worst model reaches 0.197. Reweighted models push predictions toward the tails and under-predict the mid-band by up to 26 percentage points (Figure 5, panel A,B; full 36-configuration ranking in Supplementary Data 4).

**Figure 5:**
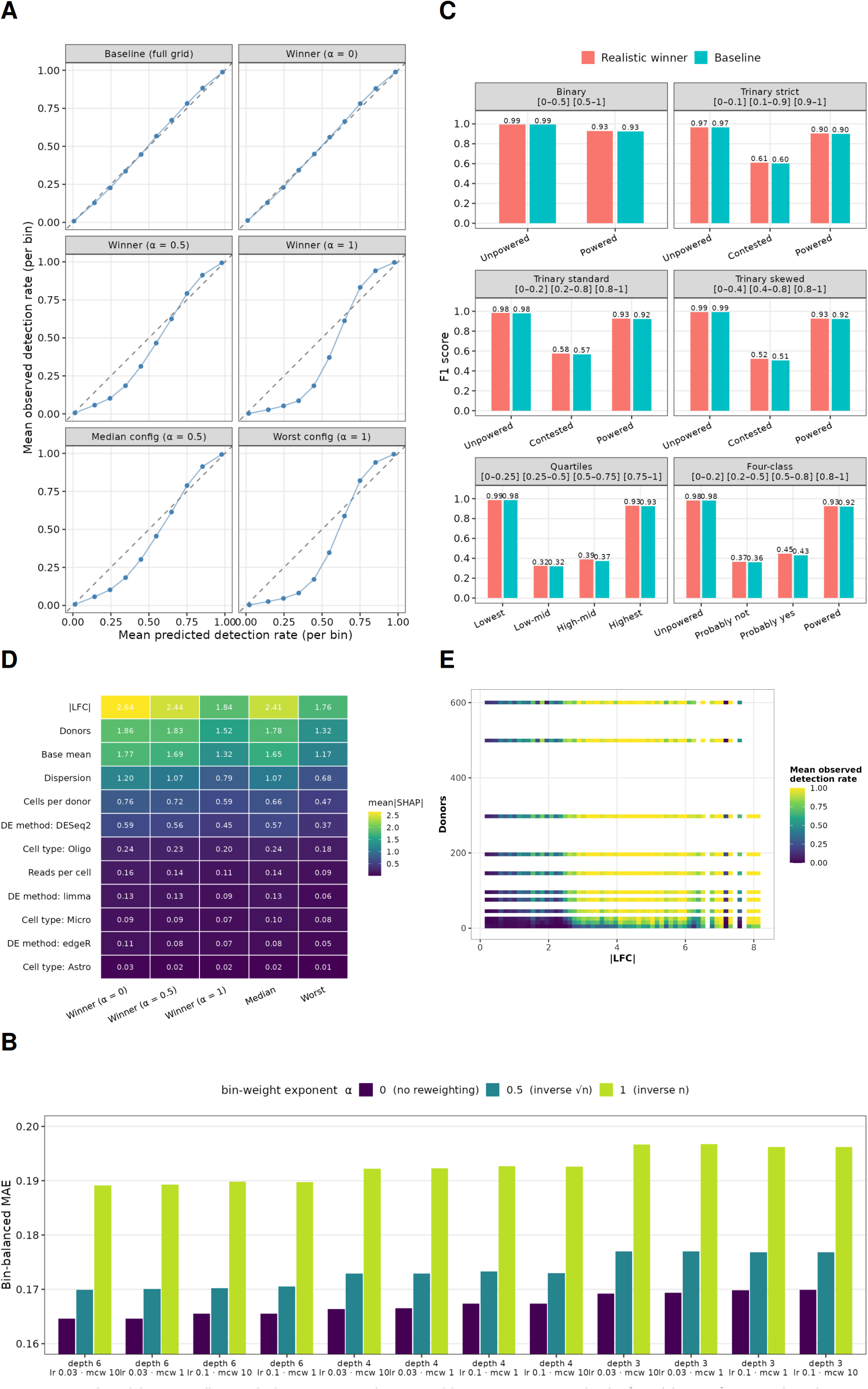
Trained models are excellent at declaring a gene to be impossible or near-guaranteed to be found, but perform poorly in the contested region, and gene-level characteristics are important and consistent predictors. Calibration is defined by how close the model’s power predictions were to the observed value. (A) Out-of-fold calibration curves for the baseline (full-grid) model and five representative realistic models (only trained from pseudo-experiments with 100 or greater cells per donor). The winning model (*α* = 0, depth 6, learning rate 0.03) tracks the diagonal closely at both extremes; loss reweighting (*α* > 0) degrades mid-band calibration progressively. (B) Bin-balanced mean absolute error for models trained on each of the 36 hyperparameter configurations. There is a clear trend of model performance improving with increased tree depth, and minimum child weight, but decreased learning rate and loss weighting. (C) F1 score of six classification methods using the output of the best performing model, against a model trained using the full dataset. Predictions and targets are converted to their respective classes, from which precision and recall metrics are calculated. (D) Mean absolute SHAP importance across five selected configurations; |LFC| dominates across all models, donor count and mean expression rank second and third, and cell type, DE method, and per-cell depth contribute marginally and stably. (E) Mean observed detection rate over the |LFC| × donor-count grid for the best performing model: the design space is bimodal, with the transition band (0.1-0.9) occupying a narrow slab.

The predictions were almost perfect at the extremes: for the winning model, 97.6% for predictions in the least detectable range (0-0.1), and 98.0% in the most detectable range (0.9-1) land within 0.10 of the observed detection rate. Mid-band bins (0.2-0.8) are substantially worse: the proportion within 0.10 collapses to 23-27%, MAE plateaus at 0.22-0.24. Only 7.8% of grid rows fall in the (0.1, 0.9) detection-rate band, 4.3% in (0.2, 0.8), and 2.6% in (0.3, 0.7); the vast majority of the design space is either safely powered or completely unpowered. The poor calibration in the middle is a symptom of measuring the regions most uncertain (highest entropy), partly because they carry the highest Bernoulli variance: flipping a fair coin 20 times will have a much larger range of feasible possibilities than a heavily weighted one. The “contested” region in the middle will also be most severely affected by a small change in each of the predictors, and thus measurement noise for each of the variables affects the probability of detection more than at the extremities. Given the unreliability of raw values, we grouped each prediction into bins and quantified each bin’s F1 score, a metric reconciling recall and precision into one value (Figure 5, panel C). These all rely on the original predictions, and are not retrained models. We compare the best performing model with its baseline counterpart, which includes the datapoints with cells per donor < 100 for each classification, finding little to no improvement (Figure 5, panel C). Trinary skewed classification was chosen as the most useful due to high performing regions showing an experiment to be confidently underpowered or powered, with a smaller contested section between.

Feature importance rankings are stable across the selected configurations (Figure 5, panel D): |LFC| dominates, followed by donor count and mean expression, with cells-per-donor, UMIs per cell, cell type, and DE method contributing marginally. This ordering should not be over-interpreted, as feature importance in tree ensembles also reflects feature range. |LFC|, while a powerful indicator of power, is not sufficient alone, as some high |LFC| genes had consistently poor power across experiments (Figure 5, panel E). Cells per donor, and UMIs per donor are correlated with mean expression. This means that the signal for increased sampling in a per-donor profile is spread across several variables, which should be considered when comparing the contributions of each feature to predicted power. Similarly, counts per cell is correlated with cell type as the mean expression is noticeably different between them. These relationships would normally be accounted for through PCA, or similar, to reduce the feature space, however, it was preferable to keep variables directly related to a single observable for simpler interpretation.

### Different Groups of Genes, Different Power Requirements

Expanding the idea of per-gene power prediction we draw on three external gene annotations: the curated catalogue of human transcription factors of Lambert et al.^22^; the set of loss-of-function intolerant (dosage-sensitive) genes defined by the pLI metric introduced in the ExAC study of Lek et al.^23^ (pLI ≥ 0.9)^24^; and a random set of PANTHER groups (Figure 6). We chose an arbitrary experimental condition, with three |LFC| thresholds to probe the model. Consider an experiment with 50 donors, 1000 oligodendrocyte cells per donor, five and a half thousand UMIs per oligodendrocyte, which aims to detect all DEGs with |LFC| > 2. If an experiment aims to interrogate chromatin regulators, ubiquitin ligases, or any of the loss of function-intolerant genes, then 89-96% of them are confidently detectable (with power > 80%) and the experiment is sufficient. However, if the true targets are intermediate filaments, forkhead transcription factors or cytokines, then only 30-38% are confidently detectable. This drops down marginally to 77-85% and 15-30% respectively for genes of |LFC| = 1. For |LFC| = 0.2, every single gene in the entire count matrix is undetectable for this experimental setup. Other gene groups such as pathways could benefit from similar consideration: if many constituents of a pathway are expressed at low levels, then detecting up or down regulation of the pathway as a whole will be more challenging and require more data.

**Figure 6:**
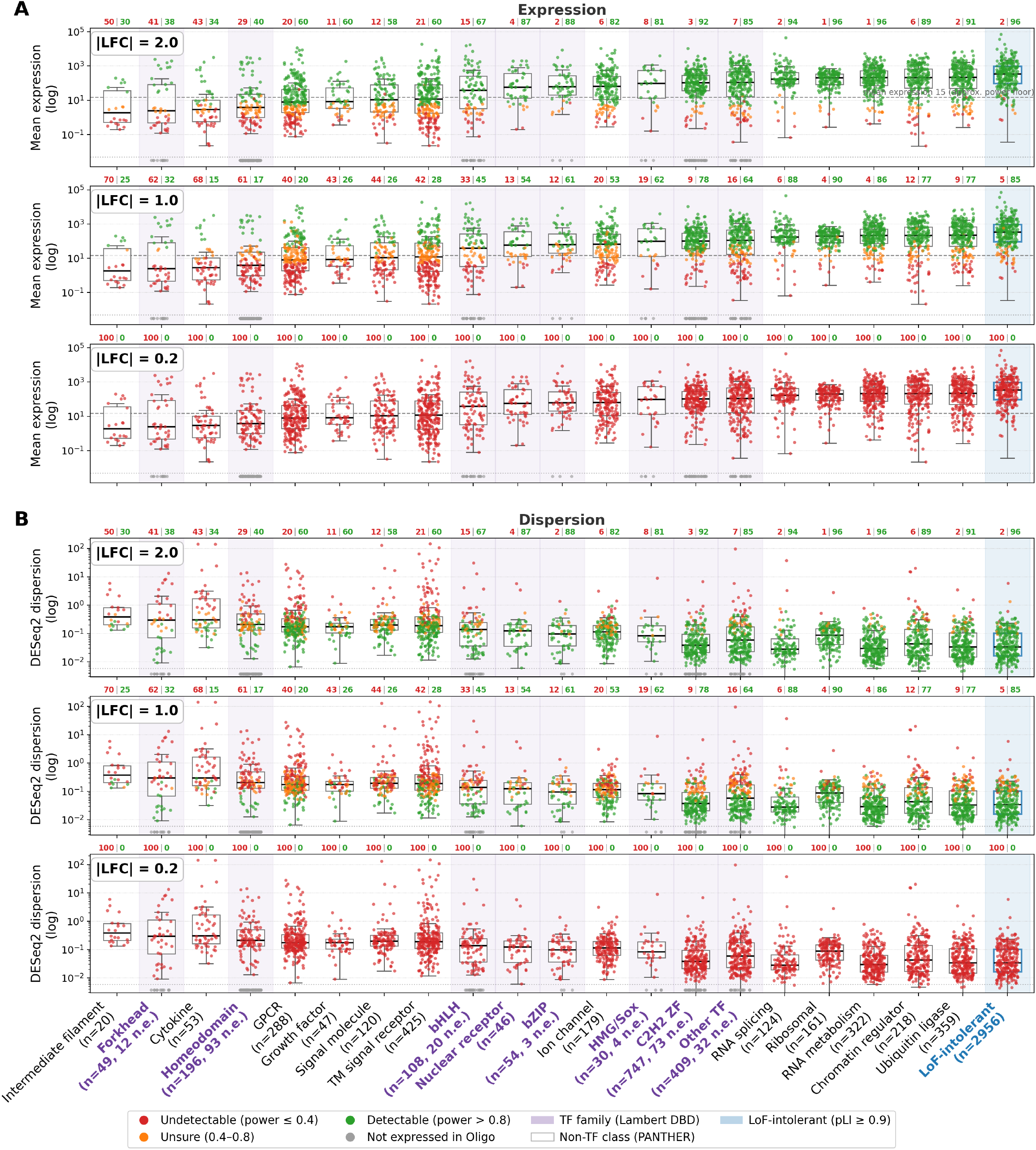
Some types of genes are inherently harder to detect than others, even when the effect size is the same. Predicted power for every oligodendrocyte gene at a fixed design (50 donors, 1,000 cells per donor, 5,742 UMIs per cell; DESeq2), classified as Undetectable (power ≤ 0.4), Unsure (0.4–0.8) or Detectable (> 0.8). Genes are grouped into the seven largest Lambert transcription-factor DNA-binding-domain families, an ‘Other TF’ catch-all (so that all 1,639 catalogued transcription factors are represented), non-transcription-factor PANTHER protein classes, and a cross-cutting loss-of-function-intolerant set (pLI ≥ 0.9). Columns are ordered by median expression (low to high) and are identical in every panel. The top three panels show raw pseudobulk mean expression (log scale); the bottom three show DESeq2 dispersion (log scale); each triplet is evaluated at |LFC| = 2.0, 1.0 and 0.2. Because expression and dispersion are effect-size-independent, each gene occupies an identical position in all panels and only its colour, giving the predicted-power class, changes with effect size. Grey points are genes not expressed in oligodendrocytes. The dashed line marks mean expression 15, the approximate power floor at |LFC| = 1.0.

## Discussion

Single-cell/nucleus RNA-seq lacks empirically validated power analyses, free from the distributional assumptions and pilot-derived parameter estimates that limit existing simulation-based power calculators. This limits the characterisation of detection power present in single-cell datasets, permitting numerous underpowered studies, and yielding a significant replication crisis. From empirically quantifying the power and reproducibility within a filtered 1,494 donor dataset, we show systematic underdetection of genes with |LFC| < 2, irreproducible experiments following as a natural consequence of this. We demonstrate individual characteristics of genes to be an important factor in detectability, and from this show that experiments which may be powered for some groups of genes, are not for others. The more cells sequenced per donor for a particular cell type, the more DEGs become detectable. We show that the sparse characteristics of microglia limits the possible discoveries, relative to other cell types. This could be mitigated by cell sorting to enrich these populations.

Standard p-value thresholds are found to yield a majority of irreproducible findings despite multiple testing correction. Standardised metadata columns, covariate modeling, and uniform software packages, all across a single cohort, give a homogeneous method surpassing the consistency expected from independent cross-lab experiments. This frames our concerning results as still an upper bound on expected reproducibility on most study to study comparisons. The picture painted by our findings should elicit methodological changes in future experiments of kind, and field-wide attention to how we should declare genes to be differentially expressed, both retrospectively and in the future.

The notion of a ground truth for empirical analysis is rarely well defined. One approach is to only declare commonly detected DEGs, but this selects only for the most obvious and easily detectable DEGs. This gives inflated power estimates and masks the full set of DEGs. Ground truth genes alternatively can be defined by the results of the largest single dataset available to researchers at the time, with downsampled pseudo-experiments both derived and validated by it^25^. This yields circular results: a ground truth of 40 donors will conclude 40 donors is sufficient to find every single DEG there is. We find when using distinct halves of over 600 donors that full agreement is still never reached, and that false positives are still prevalent at cohort sizes larger than those used in many real experiments. This motivated our choice to treat 600 donor halves as a golden ground truth. We make no assertions that any individual genes are true or false positives, but posit the scale of data used and number of repetitions should be sufficient for mitigating the impact of false positives in distorting results. Most importantly, this ensures we miss fewer true but hard to detect DEGs which help calibrate more realistic power predictions. Extrapolating this philosophy, we expect many true DEGs to be undetectable in our ground truth sets, and accordingly make no assertions of completeness. As such, these results are still optimistic compared to what it would take to identify the true set.

There are several practical applications of our work. The discretised grids of mean power for each effect size interval are useful generalisations to calibrate experimental expectations. When particular pathways or genes are of interest, we show the importance of considering gene-level characteristics. Many labs will not have access to the cohort sizes indicated in this work. For studies targeting large-effect genes, increases in experimental scale are not as necessary, but when reporting genes of small effect significant reforms are required. Where donor numbers are limited, experimenters should compensate by maximising cellular yield per donor, particularly for the cell type of interest. We recommend cell sorting tissue samples rather than a catch-all dissociation, so as to improve reproducibility: fewer, but more confident, results per study. Furthermore, due to the irreproducibility found within our “statistically significant” DEGs, we encourage researchers to employ stricter thresholds for significance, and be more sceptical of previous significant genes near adjusted p-value cutoffs.

## Methods

### Datasets and Cohort Assembly

The analysis used single-nucleus RNA-seq data of human brain tissue from the PsychAD cohort (dorsolateral prefrontal cortex; 1,494 donors after filtering)^26^. Raw data were converted to standardised donor-level shards. Each shard consisted of a single <monospace>SingleCellExperiment </monospace>object per donor, partitioned by cell type, and serialised using the _qs_ binary format. Gene identifiers were harmonised to Ensembl IDs using a shared mapping table derived from the source annotations. Three glial cell types were analysed: astrocytes, microglia, and oligodendrocytes. These were selected for their lower subtype heterogeneity than neurons and their relevance to neurodegenerative conditions.

Sex label errors have been reported in large single-cell atlases^27^, so we verified concordance for all PsychAD donors by fitting a logistic regression on pseudobulk CPM-normalised expression of XIST and Y-chromosome markers (DDX3Y, KDM5D). Four donors (0.3% of the cohort) were flagged as discordant (Supplementary Table 4): three showed simultaneous X- and Y-marker expression, a profile indicative of both sexes and consistent with Klinefelter’s syndrome or mosaicism, and one labelled-male donor lacked detectable Y-marker expression. As there was no evidence to suggest systematic mislabelling, all flagged donors were retained to better represent genetic diversity in populations. The results matched those in the original paper^26^, which showed clean labels with the exception of patients with known sex chromosome aneuploidies.

### Donor and cell filtering

Donors with age below 40, or with fewer than 30 cells in a given cell type, were excluded from that cell type’s analysis. Filtering was applied per donor, per cell type. Filtering thresholds were derived empirically from principal component analysis of the unfiltered PsychAD cohort (Supplementary Figure 1). The first principal component was dominated by per-donor cell count, with donors below approximately 30 cells forming an elongated tail consistent with gene dropout. This threshold is conservative relative to the point at which the tail becomes visible (around 20 cells), leaving a margin for cell type-specific dropout variation. The second principal component exhibited age-dependent structure, with distinct clustering in microglia for younger donors, consistent with published literature on age-dependent microglial state transitions. The 40-year threshold was chosen to stabilise the age covariate with minimal donor loss in the PsychAD cohort, while ensuring that downstream models were not dominated by developmental or early-adult biology.

Sex label accuracy was verified for all PsychAD donors by comparing recorded sex metadata against chromosomal marker expression. For each donor and cell type, pseudobulk expression was computed by summing raw UMI counts for each marker gene across all cells and dividing by the total library (sum of all UMI counts across all genes and cells), yielding a counts-per-million (CPM) value normalised for both sequencing depth and donor cell yield. Two classes of marker were used: *XIST* (ENSG00000229807), an X-inactivation transcript expected to be highly expressed in females and absent in males, and a composite Y-chromosome signal derived from *DDX3Y* (ENSG00000067048) and *KDM5D* (ENSG00000012817). For each cell type, binary logistic regression models were fitted on log_2_ (CPM + 1): one predicting *P* (female ∣ XIST CPM) and one predicting *P* (male ∣ DDX3Y + KDM5D CPM). Donors were flagged as discordant where the model-predicted probability exceeded 0.80 in the direction opposite to their recorded label. This pseudobulk CPM approach produces approximately four orders of magnitude of separation between male and female donors, making the logistic boundary unambiguous and avoiding the sensitivity of per-cell percentage-based thresholds to ambient RNA contamination.

## Differential Expression Analysis

Pseudobulk aggregation summed raw UMI counts across all cells from each donor within the target cell type, producing one count vector per donor per cell type. Pseudobulk aggregation was selected in preference to cell-level differential expression methods (such as MAST, the Wilcoxon test applied to cells, and mixed-effects models on single cells) for three reasons. First, pseudobulk treats the biological replicate (the donor) as the statistical unit, thereby modelling the hierarchical structure of the data, in which cells are not independent observations. Cell-level methods, by contrast, systematically inflate the effective sample size by treating each cell as independent, producing anti-conservative *p*-values and inflated type I error. Second, pseudobulk counts preserve count-based statistics (negative binomial distribution and dispersion estimation), allowing the use of mature count-based differential expression frameworks that have been validated on bulk RNA-seq. Third, pseudobulk aggregation is computationally tractable across the many differential expression analyses performed over the grid, whereas cell-level methods with random-effects structures would have been prohibitive at this scale. The cost is reduced sensitivity to within-donor heterogeneity, which is acceptable given that the biological question concerns population-level sex differences.

### Differential expression methods

Three pseudobulk differential expression methods were run across the full grid: DESeq2^18^, edgeR^19^, and limma-voom^20^. These three were used because they are the most pervasive pseudobulk differential expression methods in the field. Each was run with the design formula _∼ source + ethnicity + disease + age_scaled + sex_, and genes were declared significant at Benjamini and Hochberg adjusted *p*_*adj*_ < 0.05. LFC estimates from all three methods were shrunk with a single common estimator, adaptive shrinkage (ashr)^28^, so that effect sizes are directly comparable across pipelines and any apparent differences between methods reflect the underlying tests rather than differing shrinkage conventions. ashr takes each gene’s estimate and standard error and returns its posterior mean under an adaptive unimodal prior. For DESeq2 this was applied through _lfcShrink(type = “ashr”)_ ; for limma-voom the standard error was taken directly from the moderated fit (the unscaled standard deviation of the coefficient times the square root of the posterior residual variance); and for edgeR, which does not report a coefficient standard error, it was recovered from the single-degree-of-freedom quasi-likelihood F-statistic as 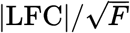on the log_2_ scale. Genes for which a valid positive standard error could not be formed retained their unshrunk estimate. Running all three methods across the grid allows the differential expression method itself to be carried as a covariate of the power model, rather than fixing a single tool.

A threshold of |LFC| > 0.2 was retained for the expanded truth set used in grid downsampling results reporting and the power curve fitting, in order to ensure that the models could be calibrated across a wide range and attempt to model realistic effect sizes typically reported in differential analysis studies.

## Grid Downsampling Framework

### Ground truth and validation split

The filtered PsychAD cohort was randomly partitioned 50/50 into ground truth and validation sets using sex-stratified sampling, which preserves the global sex ratio in each half. This partition was repeated for ten independent robustness runs with different random seeds. Ground truth differential expression was performed on all donors in each ground truth half in order to establish the reference set of differentially expressed genes. Validating power against a held-out truth set guards against the circularity of measuring recall on the same donors used to define truth, also known as data leakage. Fully downsampling the entire dataset would always converge at 100% recall and precision, regardless of dataset size, yielding unreliable power estimates.

The 50:50 ground truth split (half of the donors for establishing truth, half for downsampling) was chosen to maximise the spread of experimental setups that could be tested, but has not been subjected to sensitivity analysis; alternative splits (e.g., 70:30 or 80:20) might yield more robust power estimates in exchange for fewer experimental conditions tested. These modelling choices do not affect the qualitative conclusions: the power ordering, depth censorship effects, and donor-versus-cell trade-offs emerge directly from the empirical grid. Researchers planning experiments without access to 600 donors may wish to shift the split ratio in favour of the ground truth, aiming for more robust predictions in their feasible experimental space, but should be cautioned that precision estimates appeared to deteriorate as donor count increased for the tiny-effect size bin, perhaps indicating that a ground truth with more donors but no deeper sequencing, may strengthen noisy false positives, instead of producing more robust findings.

Sex-stratified sampling was adopted because, without stratification, random splits in small-denominator cell types would occasionally produce sex-imbalanced halves that distort the truth set. Stratifying by the target case-control variable (here, sex) is standard practice in case-control power analyses and eliminates this source of iteration noise.

### Grid design

The validation set was downsampled across a grid of 12 donor counts by 14 cell counts per donor, with 20 iterations per grid point. The donor counts were 5, 10, 15, 25, 50, 75, 100, 150, 200, 300, 500, and 600. The cell counts per donor were 5, 10, 20, 50, 75, 100, 250, 500, 750, 1,000, 1,500, 2,000, 3,000, and a “Max” level representing the all-available-cells configuration. Iteration seeds were derived deterministically from the robustness run, number of donors, number of cells (or read budget), and iteration index. This enables precise debugging and guarantees reproducibility.

Both axes are spaced approximately logarithmically in order to concentrate resolution where detection probability is changing most rapidly (at low donor and cell counts) and to cover multiple orders of magnitude efficiently. The upper cell bound of 3,000 exceeds the per-donor cell count available for most donors, so the “Max” point captures the all-available-cells configuration. Cells were sampled without replacement, and donors were excluded if they had insufficient cells to match the grid point’s count requirements. Grid points that had insufficient donors with enough cells were not performed and are represented in grey in results to indicate an untested region of the search space. Each subsample required a full pseudobulk differential expression analysis under each of the three methods, yielding a substantial compute cost.

### UMI-based mode

A parallel UMI-based downsampling mode replaced the cells-per-donor axis with a total-UMI-per-donor budget axis, spanning 25,000 to 15,000,000 UMIs across 13 levels including a “Max” level, yielding 156 grid points (12 donor counts by 13 read counts). For each donor at each read budget, cells were drawn sequentially and accumulated until the running UMI total exceeded the target; the final “overflow” cell was subjected to multinomial downsampling so as to bring the total exactly to the target, while stochastically preserving representative gene-level count proportions.

The cells-based mode answers the question “for a given number of cells per donor, what power is achieved?”, while the UMI-based mode answers the complementary and arguably more common planning question “for a given sequencing budget per donor, what power is achieved?”. These are not equivalent questions, because cells vary widely in UMI yield (astrocyte cells carry approximately twice the UMIs of microglia cells), and a fixed cells-per-donor number therefore implies different total read budgets across cell types. For experimental planning at a fixed monetary or flow-cell budget, the UMI-based parameterisation is the more appropriate decision axis. The overflow multinomial downsampling is necessary because UMI budgets never land exactly on a cell boundary; stochastic multinomial downsampling of the overflow cell preserves expected gene-level proportions while delivering the exact target budget.

### Recall and precision metrics

Detection power was summarised by two complementary metrics, computed for every grid point (a combination of donor count, cells-per-donor or UMI budget, cell type and differential expression method) and then, unless stated otherwise, mean-pooled across the three differential expression methods. For a given grid point, recall is the mean proportion of the ground-truth differentially expressed genes that were recovered (declared significant at *p*_*adj*_ < 0.05) in the corresponding downsampled pseudo-experiment, averaged over the twenty iterations and ten robustness runs; a recall of 1 indicates that every ground-truth gene was recovered. Precision is the mean proportion of the genes declared significant in the pseudo-experiment that were also present in the ground-truth set, computed over the same iterations and runs; a precision of 1 indicates no false positives relative to the ground truth. Formally, for a grid point with ground-truth gene set *G* (restricted to the relevant bin) and letting *D*_*r*,*i*_ denote the set of genes declared significant in robustness run *r* and iteration *i*,

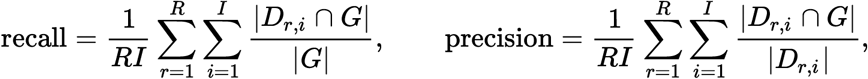

where *R* = 10 robustness runs and *I* = 20 iterations per grid point. Both metrics were computed within each effect-size bin (tiny |LFC| 0.2-1.0, small 1.0-2.0, medium 2.0-3.0, large >3.0) and, where indicated, stratified by chromosome group (autosomal versus sex-chromosome).

To probe what drives recall beyond effect size, the ground-truth genes were further stratified in two ways at a fixed design. First, genes were partitioned into quartiles of their ground-truth Benjamini-Hochberg *p*_*adj*_-value, from the most significant (Q1) to the least significant (Q4), and recall was recomputed within each quartile. Second, within a single effect-size bin, genes were grouped into four mean-expression categories by DESeq2 base mean (high, 200+; medium, 200-50; low, 10-50; and very low, <10), and recall was recomputed within each category. Both stratifications reuse the same per-gene detection outcomes as the pooled metric; only the grouping of the ground-truth genes changes.

## Power Curve Fitting

### Detection-rate response

For each grid subsample and each gene in the ground truth gene set, a binary detection outcome was recorded: 1 if the gene reached *p*_*adj*_ < 0.05 in that subsample’s differential expression analysis, and 0 otherwise. These outcomes were aggregated to a per-grid-point detection rate for each gene, defined as the fraction of iterations in which the gene was detected at a given combination of design parameters, cell type and differential expression method. Each detection rate carried its number of underlying trials, which was used as an observation weight during fitting so that grid points supported by more iterations exerted proportionally more influence.

### Gradient-boosted tree model

The power model is a gradient-boosted decision tree ensemble (XGBoost)^21^ trained with a binary-logistic objective on the weighted per-grid-point detection rates. There is no direct prediction equation; instead power is obtained as an iterative decision tree.

The model uses twelve features. Six are continuous and enter on their raw, untransformed scale: the number of donors, the number of cells per donor, UMIs per cell, and the gene-level characteristics |LFC|, base mean, and dispersion. The remaining features are one-hot encodings of the cell type (astrocyte, microglia, oligodendrocyte) and of the differential expression method (DESeq2, edgeR, limma-voom).

Including the method as an explicit feature lets the model capture systematic differences in detection behaviour between the three pipelines rather than averaging over them. Monotonic constraints were imposed on the continuous features according to their known direction of effect on detection, so that predicted power varies monotonically with each design parameter and gene characteristic.

### Training and model selection

Hyperparameters were tuned by grouped cross-validation, with grouping applied so that all observations from the same grid point were held out together, preventing leakage between training and validation folds. The hyperparameters optimised were: alpha, a parameter for reducing the emphasis on extremes, where most datapoints lie; depth, the maximum depth of each tree; learning rate, determining the amount of correction the new tree is allowed to make; and minimum child weight, determining if a new tree makes enough difference to include. A class-imbalance reweighting sweep was run alongside the search to test whether up-weighting rarer detection outcomes improved calibration in the underpowered regime. The production model was trained on the realistic sub-grid, restricted to grid points with at least 100 cells per donor, reflecting the range of designs that real experiments occupy. Model quality was assessed by cross-validated log loss and by a bin-balanced mean absolute error that averages error evenly across effect-size and expression strata, so that the abundant easy-to-detect genes do not dominate the metric. Writing _S_ for the set of strata and *B*_*s*_ for the out-of-fold observations in stratum *s*, with observed detection rate *y*_*j*_ and predicted power 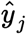,

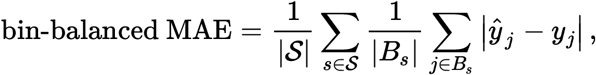

so each stratum contributes equally regardless of how many observations it contains. Feature contributions were characterised using both gain and SHAP values, the latter attributing each prediction to its features and confirming that design parameters and gene characteristics both move predictions materially.

### Classifier evaluation by power class

Because the continuous power predictions are least reliable in the intermediate (contested) region, the model was additionally evaluated as a discrete classifier. The observed and predicted detection rates were each discretised into ordered power classes under several partitioning schemes (a binary 0.5 cut, three trinary schemes with different low- and high-class boundaries, quartiles, and a four-class scheme), and a per-class F1 score, the harmonic mean of precision and recall, was computed by comparing the predicted class against the observed class within each scheme. These F1 scores reuse the fitted model’s out-of-fold predictions and involve no retraining. The production (realistic, *n*_cells_ ≥ 100) model was compared against the baseline full-grid model under every scheme (Figure 5, panel C).

### SHAP Analysis

To interpret the fitted power model, we quantified the contribution of each predictor using SHAP (SHapley Additive exPlanations) values^29^. The TreeSHAP algorithm^30^ was applied to the XGBoost models to decompose every per-observation prediction into additive, feature-level contributions on the log-odds scale, and global feature importance was summarised as the mean absolute SHAP value across a uniform random subsample of the training grid (200,000 observations)^30^. We report SHAP-based importance in preference to the model’s internal split-gain because gain attributes training-loss reduction and is sensitive to feature redundancy, whereas mean absolute SHAP measures each feature’s realised effect on the model’s predictions. This is therefore a more faithful ranking of predictive influence. This analysis spanned all twelve model features (absolute log-fold-change, dispersion, base mean, donor count, cells per donor, reads per cell, and the one-hot-encoded DE method and cell type).

### Gene group detectability by functional class

To relate predicted detectability to gene function, every oligodendrocyte gene was passed through the fitted power model at a single fixed experimental design (50 donors, 1,000 cells per donor, 5,742 UMIs per cell, the oligodendrocyte mean) while holding the effect size constant across all genes, and evaluating three effect sizes in turn (|LFC| = 0.2, 1.0 and 2.0). Each gene supplied its own two intrinsic characteristics, its raw pseudobulk mean expression and its DESeq2 dispersion from the full-cohort sex-design fit, exactly the quantities the model was trained on; the design parameters and effect size were fixed, so only expression and dispersion varied between genes. Predicted power was converted to a skewed trinary call (Undetectable, ≤ 0.4; Unsure, 0.4 to 0.8; Detectable, > 0.8), the thresholds being deliberately asymmetric to reserve the “Detectable” label for genes with high predicted recall. Genes were then grouped by function using two external annotations. Transcription factors were partitioned by DNA-binding-domain family from the Lambert catalogue^22^: the seven largest families were retained as named groups and all remaining transcription factors were pooled into a single “Other” group, so that every catalogued transcription factor is represented, including those absent from the oligodendrocyte pseudobulk (shown as not expressed rather than assigned a power). Non-transcription-factor genes were grouped by PANTHER protein class, selected to span the full expression range from broadly expressed housekeeping machinery (for example ribosomal and RNA-processing proteins) to low-abundance secreted and cell-surface signalling classes (for example cytokines and G-protein coupled receptors). Only groups with at least eight genes were kept. A cross-cutting set of loss-of-function-intolerant (dosage-sensitive) genes, defined by pLI ≥ 0.9 from the gnomAD v2.1.1 constraint release^24^, was added as an additional group. Groups were ordered by median expression to make the joint dependence of detectability on expression and dispersion visible.

## Depth Equalisation

In order to test whether apparent cell type differences in power were driven by per-cell type differences in ground truth sequencing characteristics, astrocyte cells were multinomially downsampled to match microglial sequencing characteristics. Three astrocyte truth set variants were generated. The “Native Astro” variant used the original astrocyte cells, with no downsampling. The “Astro (Depth-Equalised)” variant applied multinomial downsampling to each astrocyte cell’s UMI vector so as to match the microglia mean of approximately 4,426 UMIs per cell, using a downsampling factor of 0.4679 (the ratio of microglia mean UMIs to astrocyte mean UMIs). The “Astro (Depth+Cell-Equalised)” variant was depth-equalised as above, and additionally had per-donor cell counts randomly removed from the ground truth sets so as to match the microglia per-donor cell count distribution in their ground truths. Removing cells only from the ground truth set allowed for copying the exact downsampling iteration jobs as the full astrocyte set for a fair comparison, but still testing if ground truth depth is the problem.

Multinomial downsampling of UMIs was adopted in preference to random per-cell dropout because it preserves the expected gene-level count proportions within each cell while reducing the total count, producing cells that are statistically equivalent to shallowly sequenced versions of the originals. Alternative approaches such as proportionally reducing all UMI counts per gene or whole-cell removal would introduce biases that do not correspond to the technical phenomenon being simulated (namely, differential sequencing depth per cell).

The full grid downsampling and power curve fitting pipelines were rerun on these two new variants, using the same ground truth and validation partitions as the primary analysis on astrocytes. Using identical donor assignments between the native and equalised analyses ensures that differences in recall reflect the depth manipulation rather than sampling variation in the partition itself. This constitutes a tighter comparison than independent repartitioning would allow.

## Literature Survey

To provide a rough calibration of donor counts and LFC thresholds used in current brain sc/snRNA-seq differential-expression practice, we assembled a non-systematic survey of 30 primary human-tissue studies (2022–2026) across neurodegenerative and neuropsychiatric conditions. Candidate studies were identified through three complementary streams: an AI-assisted literature search using Edison (an LLM-based research tool)^31^, a parallel search and excerpt extraction performed with Anthropic Claude^32^, and a manual search of PubMed, bioRxiv, and citation trails from datasets used in this paper. The pooled set of candidates was then collated and parsed with Claude into a single structured table capturing donor counts, and LFC threshold where applicable. We emphasise that this exercise was intended solely as an approximate characterisation of the design space the field currently operates in, not a systematic review, a meta-analysis, or a critique of any individual study’s methodology or analytical choices; given the substantial heterogeneity in tissue, disease, cohort structure, and statistical framework across these studies, like-for-like methodological comparison would not be appropriate, and no such comparison is attempted here.

## Reproducibility and Computational Environment

All analyses are deterministic given a fixed random seed at each level of the hierarchy (partition, iteration, and subsample). Analysis pipelines are organised by functional step (_sharding/_, _full_ground_truth/_, _Cartesian_downsampling/_, _power_curve_fitting/_, _de_diagnostics/_, _depth_equalization/_, _eda_roussos/_, _figure_data_generation/_) and are publicly available (see the Code and Data Availability section), enabling researchers to reproduce the fitted power model and estimate detection probability for arbitrary combinations of design parameters and gene characteristics.

Splitting the code by functional step, rather than by dataset or cell type, enables partial reruns when a single step is updated, independent testing of each stage on toy data, and reuse of individual pipelines (in particular grid downsampling and power curve fitting) for case-control variables other than sex, or alternative statistical methodologies.

## Code and Data Availability

This study reuses previously published data; no new primary data were generated. Accession identifier, licence, and primary citation is summarised in Supplementary Data 1 and recapitulated here.

Processed snRNA-seq count matrices are available from CELLxGENE Discover for PsychAD (collection 84ce6837-548d-4a1f-919f-0bc0d9a3952f);^26^.

Source data for the main and supplementary figures are provided in the Supplementary Data files.

## Acknowledgements

We thank the donors and their families whose tissue donations made the underlying cohorts possible. We acknowledge the data-generating consortia as follows. The PsychAD Consortium data were generated with support from the contributing brain banks and associated NIMH, NIA, and NINDS awards as detailed in Fullard et al.^26^. This work is supported by the UK Dementia Research Institute award number UK DRI-5008 through UK DRI Ltd, principally funded by the UK Medical Research Council. This work was supported by UK Research and Innovation (UKRI) through Future Leaders Fellowships [grant numbers MR/T04327X/1 and UKRI2755]. P.M.M. acknowledges generous personal support from the Edmond J Safra Foundation and Lily Safra and an NIHR Senior Investigator Award. E.P.D. is supported jointly by funding to P.M.M. from the UK Dementia Research Institute and from the NIHR Imperial Biomedical Research Centre Multiple Long-Term Conditions theme.

B.K. was funded by the Wellcome Trust as part of the Advanced Therapies for Regenerative Medicine Wellcome Trust PhD Program (218461/Z/19/Z). L.B. received support from the UK Research and Innovation (UKRI) Medical Research Council (MRC; MR/W029790/1) and the Marmaduke Sheild Fund. This research was supported by the NIHR Cambridge Biomedical Research Centre (NIHR203312). The views expressed are those of the authors and not necessarily those of the NIHR or the Department of Health and Social Care.

