## Supplementary Data 2 for "Single-Cell Study Designs Are Systematically Underpowered for Small-Effect Genes": README.txt

Supplementary Data 2 - Cartesian Downsampling Results (aggregated)
==================================================================

Empirical recall/precision from split-half Cartesian downsampling across the
donor x cells|reads design grid, three glial cell types (Astro, Micro, Oligo),
three DE methods (deseq2, edger, limma) plus a combined all_methods pooling.
Aggregated over robustness runs; per-run raw rows are NOT included in this
supplementary release (aggregated summaries only).

Layout
------
<cell_type>/<mode>/<method>/Aggregated_Summary_Data.qs

  cell_type : Astro | Micro | Oligo
  mode      : cells  (downsample cells per donor, reads held) 
              reads  (downsample reads per cell, cells held)
  method    : deseq2 | edger | limma        (both modes)
              all_methods                    (cells mode only - pooled)

21 files total: 3 cell types x [ cells x {deseq2,edger,limma,all_methods}
                                 + reads x {deseq2,edger,limma} ].

File format
-----------
Each .qs is an R named list (read with qs::qread). It is NOT a flat table.
Aggregated components present (per individual-method file):

  df_summary          per grid-point recall/precision (main table; ~1812 rows x 16)
  df_cart_summary     Cartesian-grid recall/precision summary
  df_bm_summary       base-mean stratified summary
  df_exprfilt_summary expression-filter stratified summary
  df_finecross_summary fine LFC x expression cross-tabulation
  df_pval_q_summary   adjusted-p quartile summary
  df_xist_summary     XIST-positive-control summary
  df_reads_summary    reads-per-cell summary

The all_methods (pooled) file contains the subset that is defined across
methods: df_summary, df_cart_summary(*), df_xist_summary, df_reads_summary,
df_finecross_summary, plus pooled_methods / pooling metadata.
(*) components not applicable to the pooled object are stored as NULL.

Reading example (R)
-------------------
  library(qs)
  x <- qread("Oligo/cells/deseq2/Aggregated_Summary_Data.qs")
  head(x$df_summary)   # per grid-point recall & precision
