## Supplementary Figures and Tables for "Single-Cell Study Designs Are Systematically Underpowered for Small-Effect Genes"

2026-09-17

#### Supplementary Figures

##### Supplementary Fig. 1. PCA gallery for unfiltered and filtered PsychAD cohorts

Principal component analysis (PCA) of pseudobulked donors showing the effect of filtering. Rows (top to bottom): $\log_2$ cell count, sex, age, brain bank source, disease status, and self-reported ethnicity. Columns: astrocytes (left), microglia (centre), and oligodendrocytes (right).

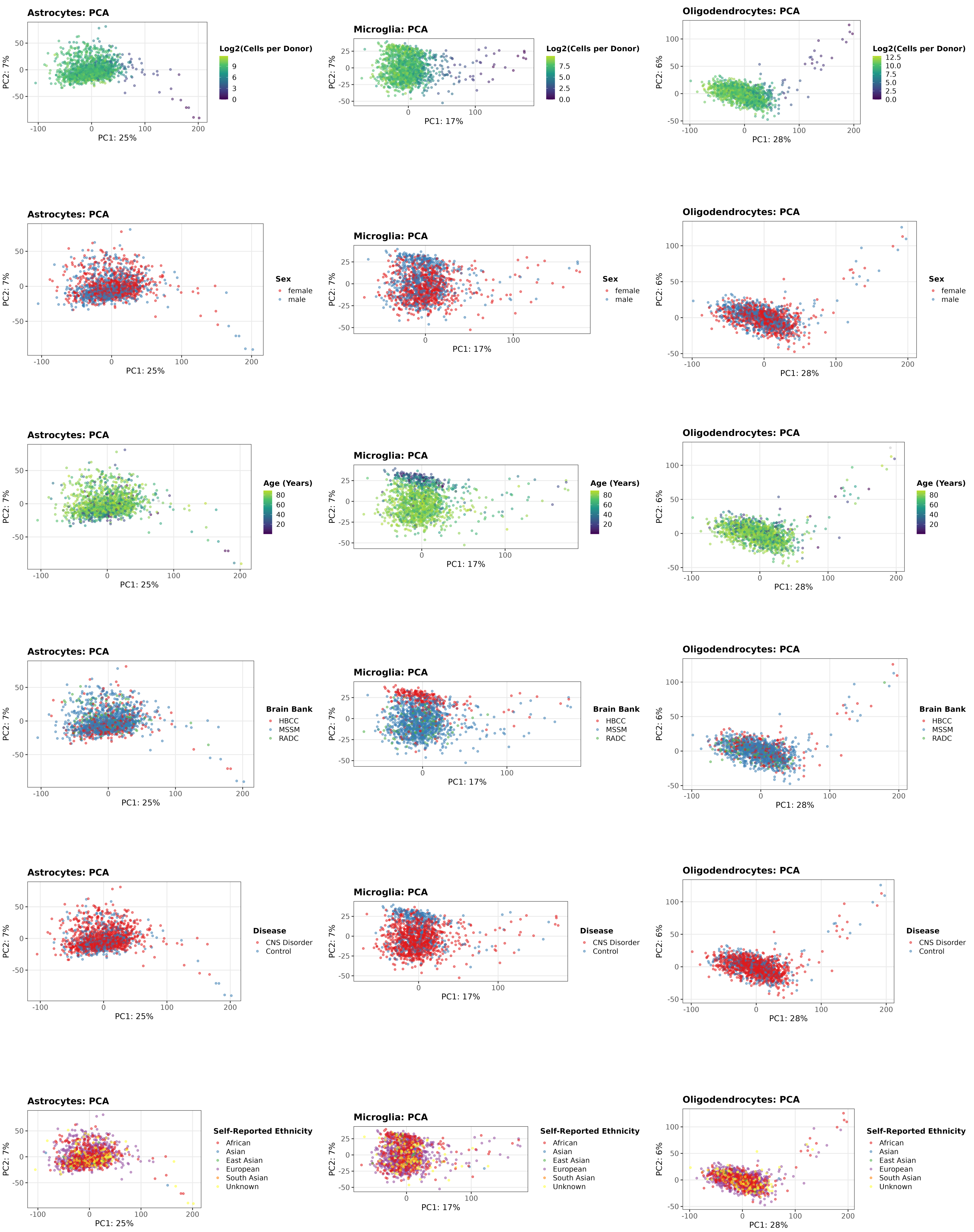

Figure 1: Complete PCA gallery for the unfiltered PsychAD dataset (N = 1,486/1,480/1,491 donors for Astro/Micro/Oligo). Each row shows one metadata covariate; columns show astrocytes (left), microglia (centre), and oligodendrocytes (right). Top to bottom: $\log_2$ cell count, sex, age, brain bank source, disease status, and self-reported ethnicity.

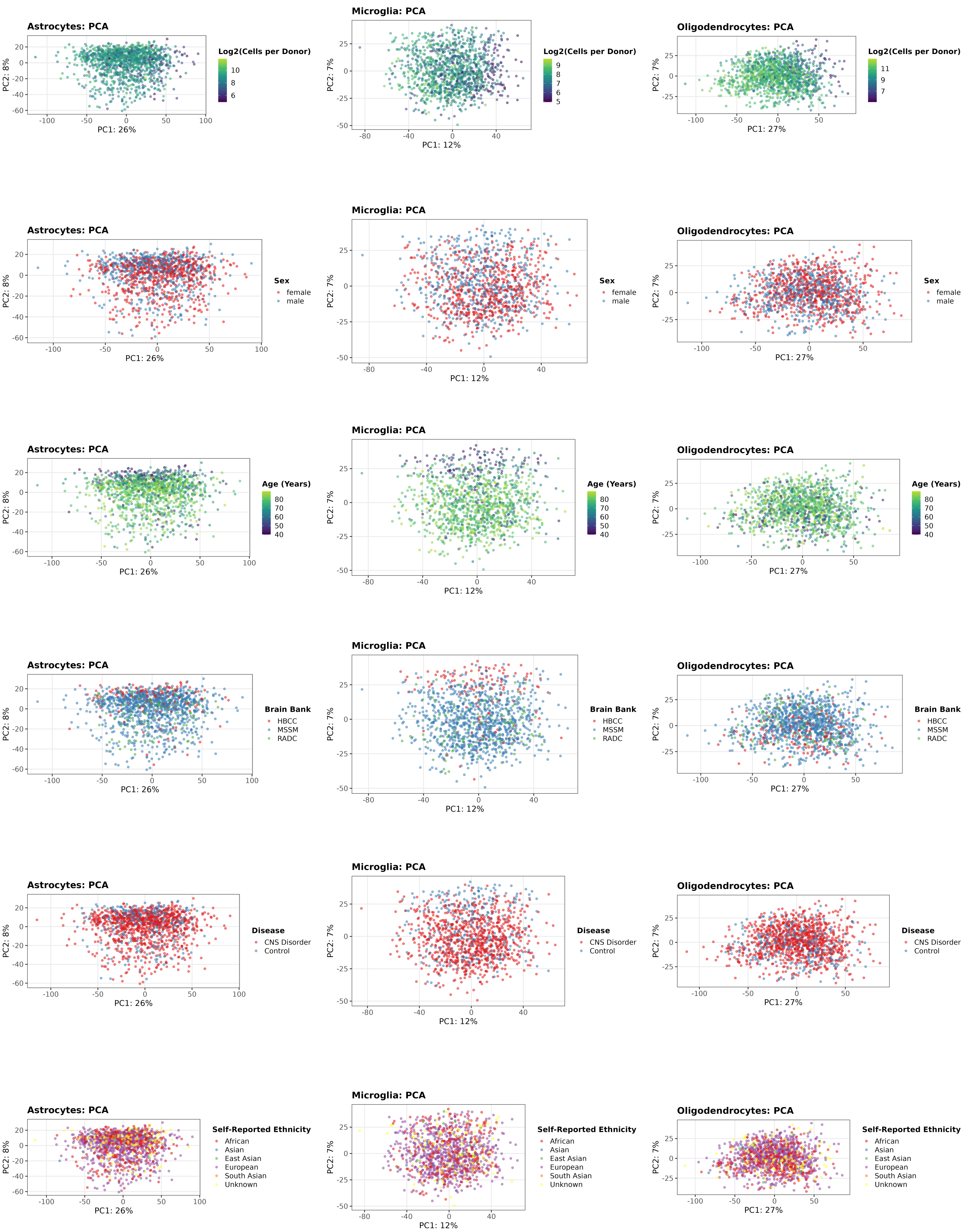


Figure 2: Complete PCA gallery for the filtered PsychAD analysis cohort (age $\geq$ 40, $\geq$ 30 cells; N = 1,293/1,214/1,303 donors for Astro/Micro/Oligo). Layout identical to the unfiltered gallery above. Compared with the unfiltered gallery, the cell-count gradient is attenuated, the outlier cluster in oligodendrocytes is absent, and the overall point cloud is more compact.

### Supplementary Figures: cross-method downsampling grids

### Supplementary Fig. 2. Cross-method recall grids (cells-based mode)

For each cell type, rows represent the DE method used to define ground-truth genes and columns represent the DE method used in the downsampled detection step. Each cell in the 3×3 grid shows recall within four LFC-magnitude bins (|LFC| 0.2–1, 1–2, 2–3, >3). High off-diagonal values indicate concordant power behaviour across methods; large between-row differences indicate that the choice of truth-definition method affects observed recall. Cells-based sampling mode.

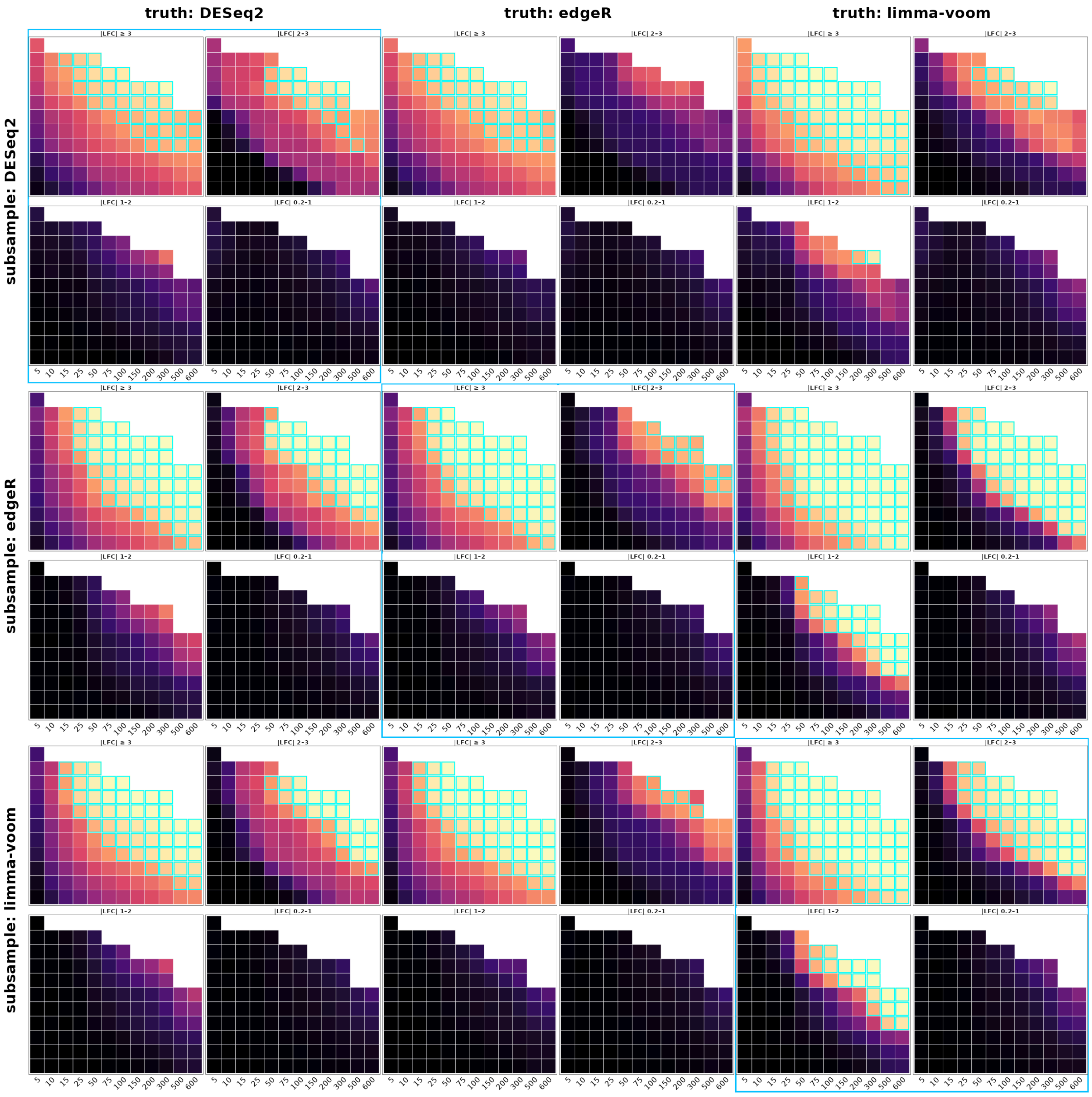

truth: DESeq2
truth: edgeR
truth: limma-voom
subsample: DESeq2
subsample: edgeR
subsample: limma-voom
|LFC| ≥ 3
|LFC| 2–3
|LFC| 1–2
|LFC| 0.2–1
5 10 15 25 50 75 100 150 200 300 500 600

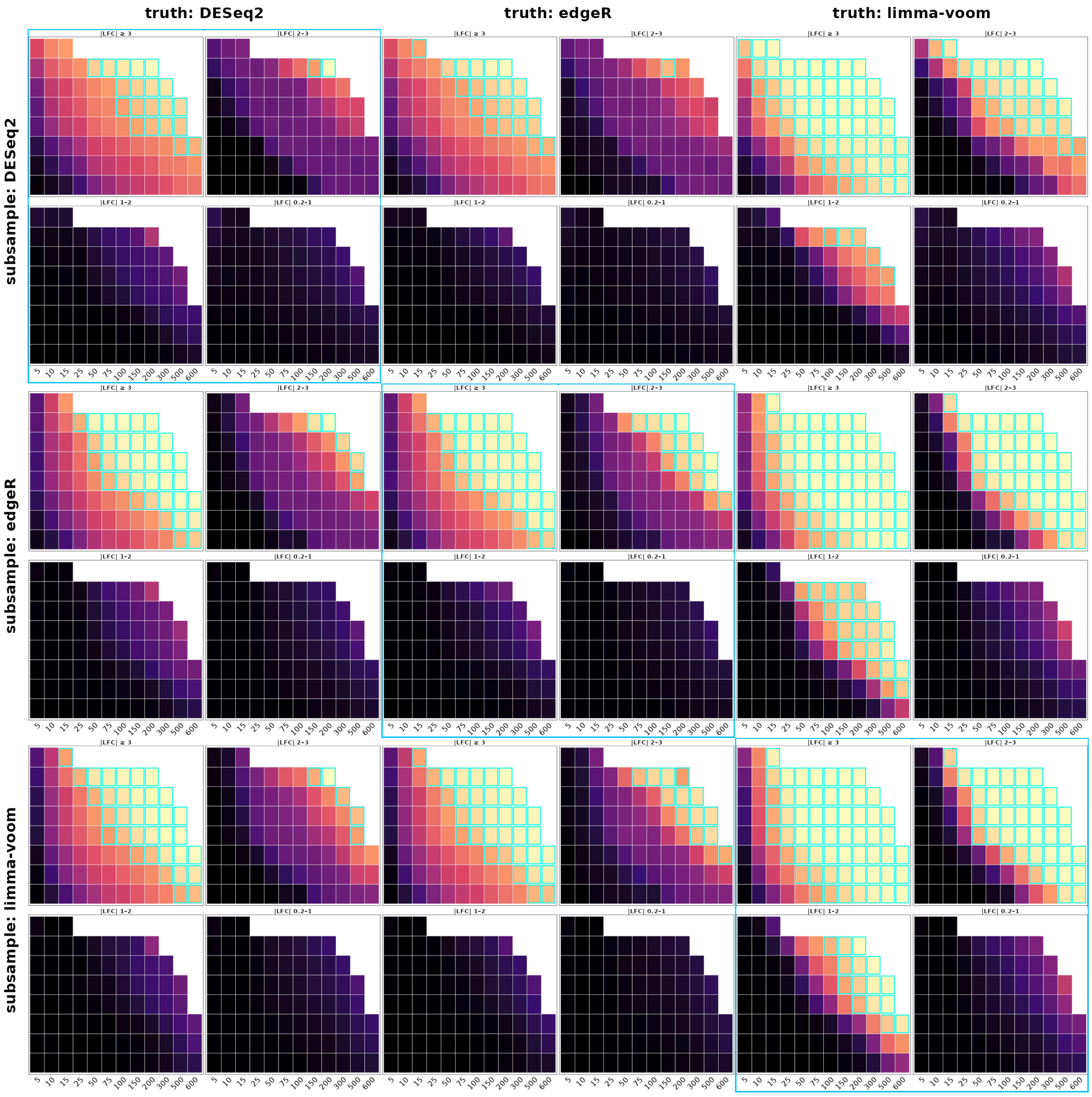

truth: DESeq2
truth: edgeR
truth: limma-voom
subsample: DESeq2
subsample: edgeR
subsample: limma-voom
|LFC| ≥ 3
|LFC| 2–3
|LFC| 1–2
|LFC| 0.2–1
5
10
15
25
50
75
100
150
200
300
500
600

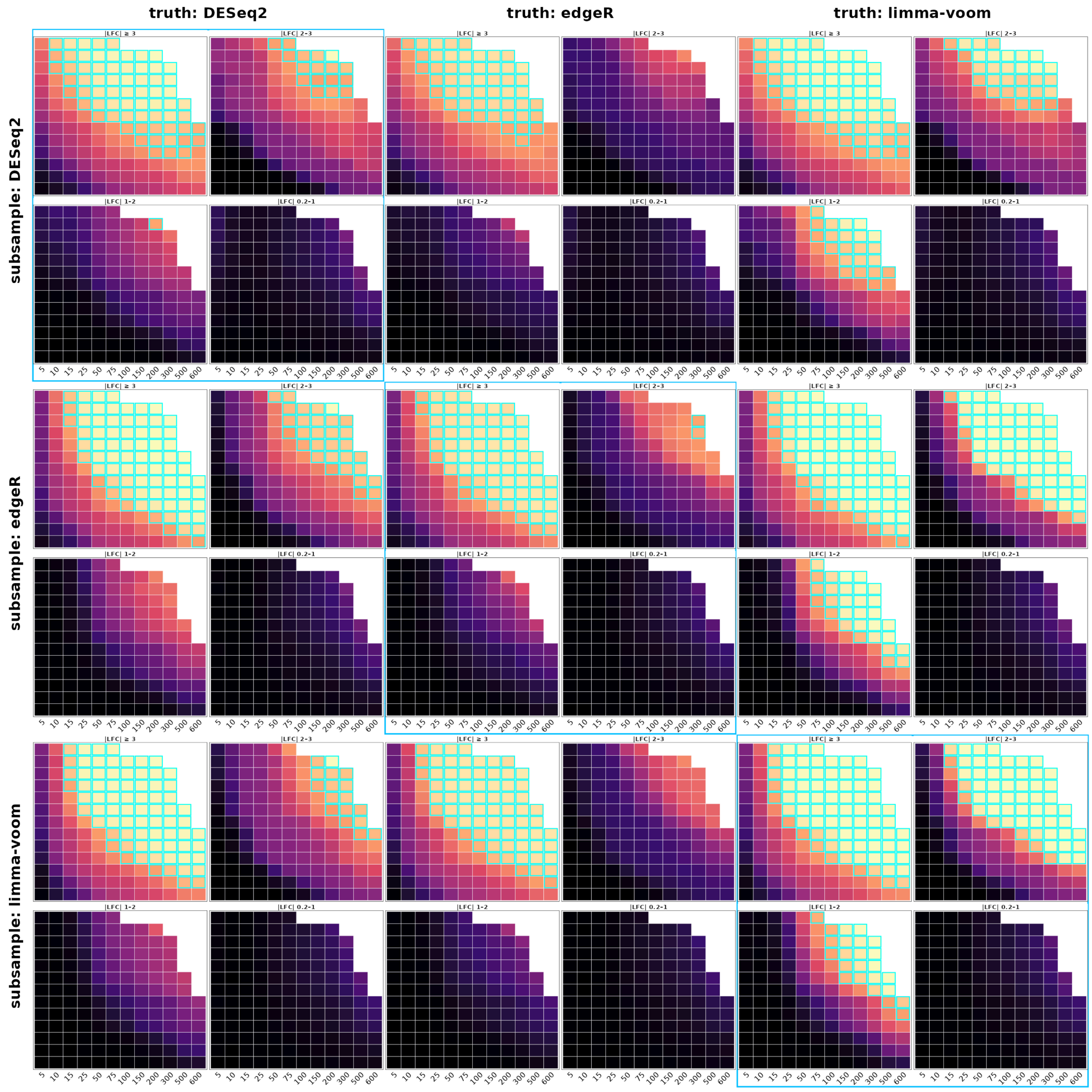


Figure 3: Cross-method recall grids (cells-based mode) for astrocytes (left), microglia (centre), and oligodendrocytes (right). Rows are the truth-defining DE method; columns are the downsampling DE method; panels show recall by LFC bin.

### Supplementary Fig. 3. Cross-method recall grids (counts-based mode)

Same layout as Supplementary Fig. 2, with counts per cell on the x-axis instead of cells per donor. Counts-based sampling mode.

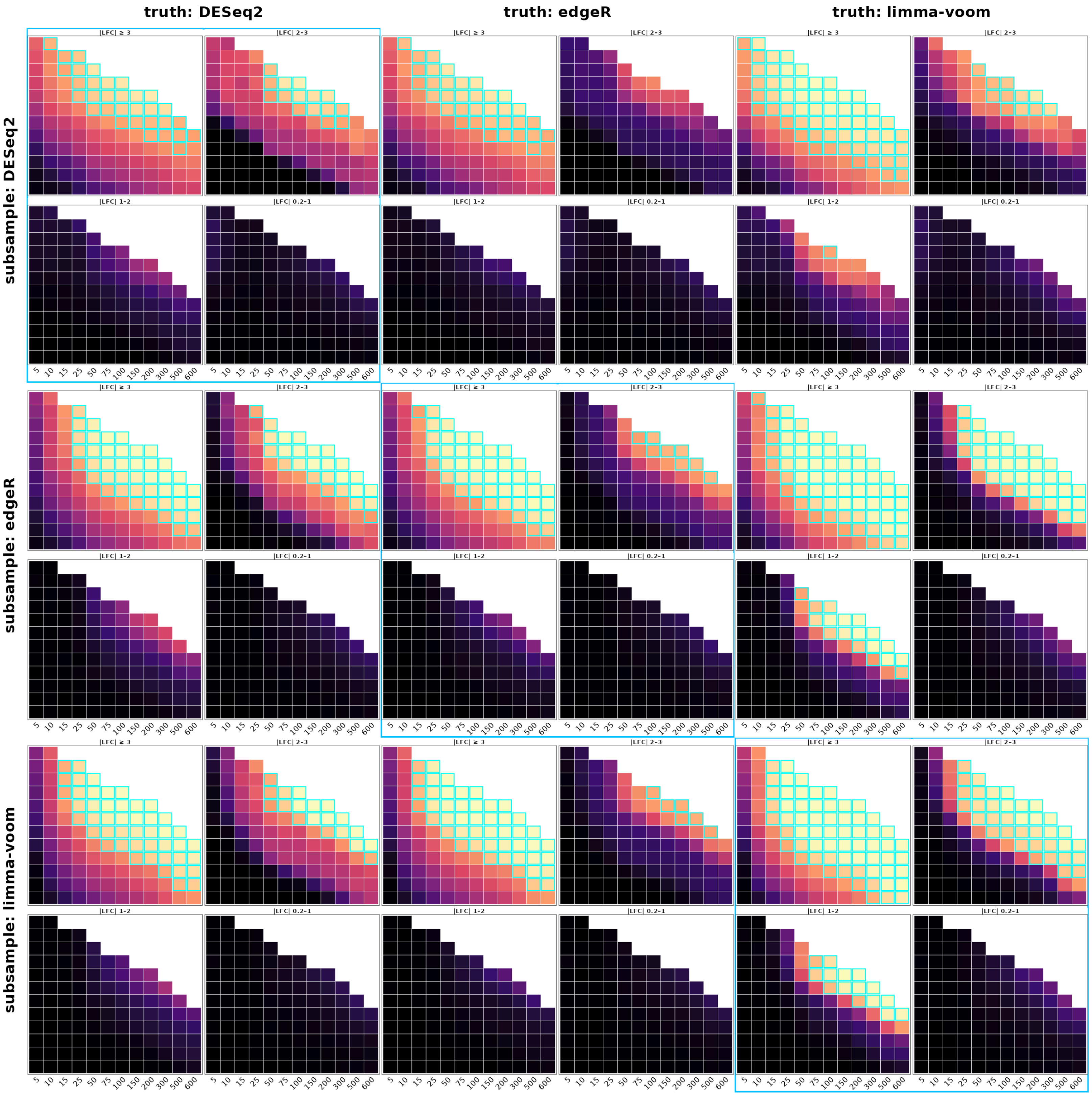

truth: DESeq2
truth: edgeR
truth: limma-voom
subsample: DESeq2
subsample: edgeR
subsample: limma-voom
|LFC| ≥ 3
|LFC| 2–3
|LFC| 1–2
|LFC| 0.2–1
5
10
15
25
50
75
100
150
200
300
500
600

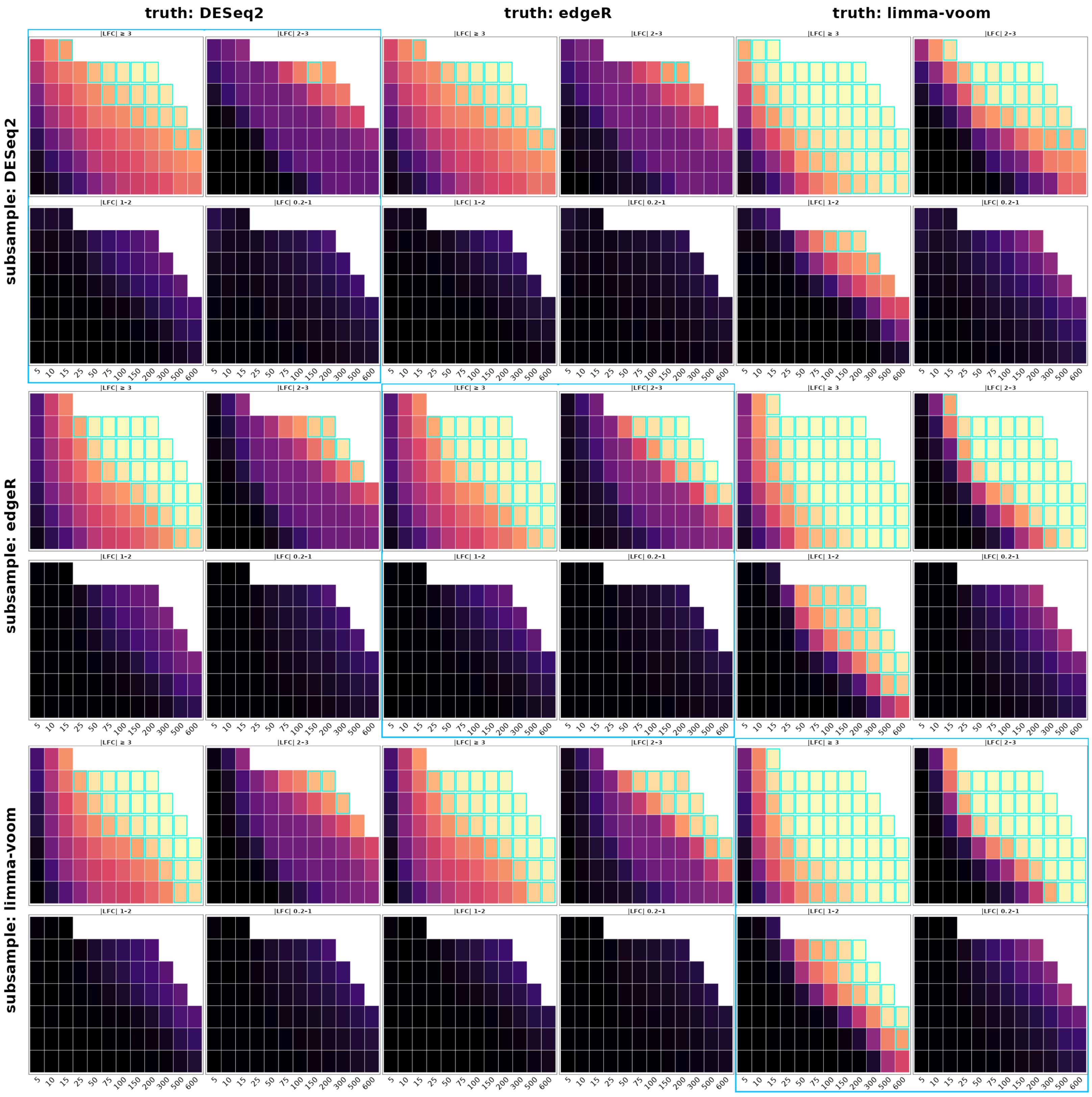

truth: DESeq2
truth: edgeR
truth: limma-voom
subsample: DESeq2
subsample: edgeR
subsample: limma-voom
|LFC| ≥ 3
|LFC| 2–3
|LFC| 1–2
|LFC| 0.2–1
5 10 15 25 50 75 100 150 200 300 500 600

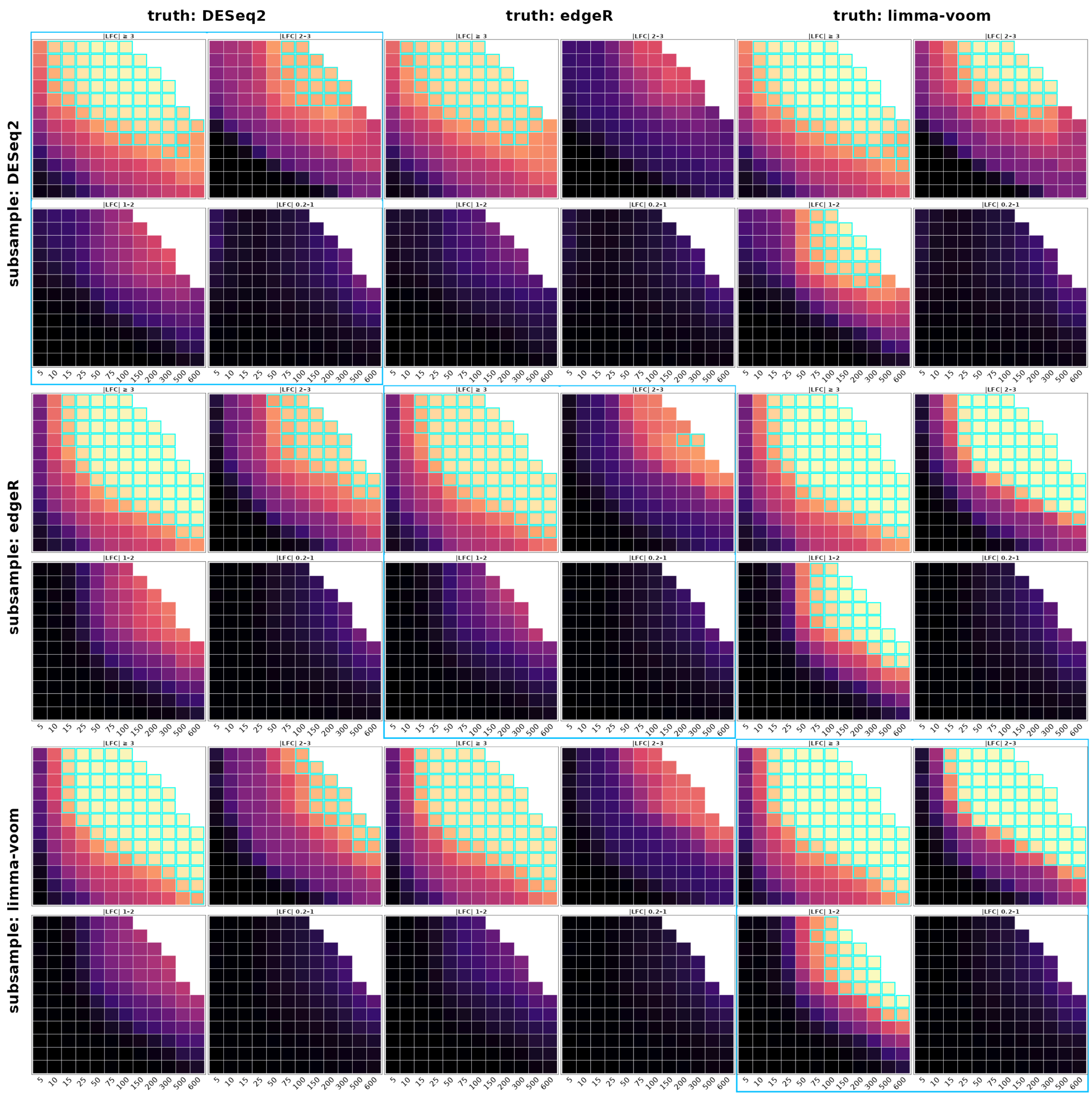


Figure 4: Cross-method recall grids (counts-based mode) for astrocytes (left), microglia (centre), and oligodendrocytes (right).

### Supplementary Fig. 4. Cross-method precision grids (cells-based mode)

Same 3×3 layout as Supplementary Fig. 2, showing precision rather than recall. Cells-based sampling mode.

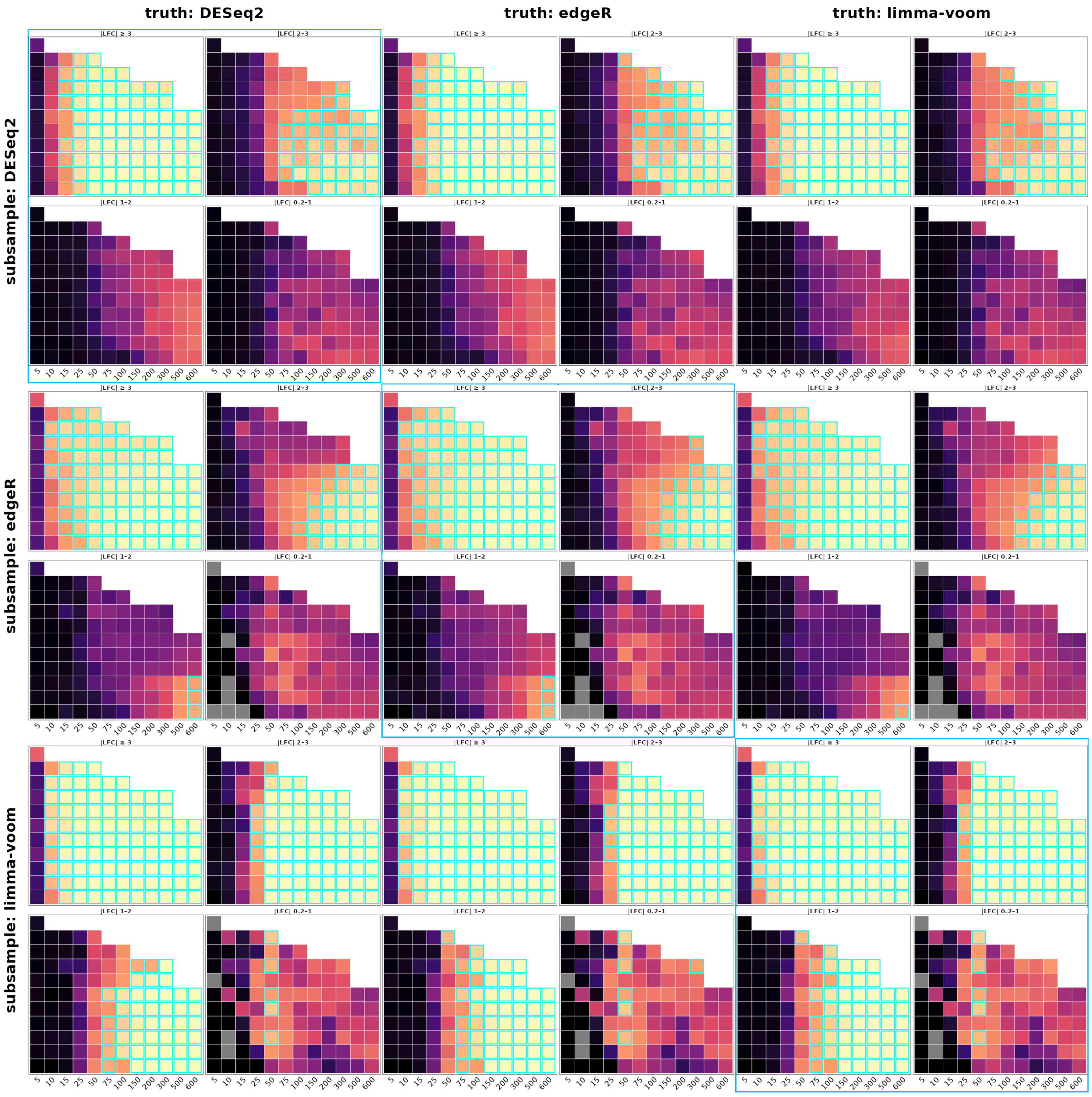
truth: DESeq2
truth: edgeR
truth: limma-voom
subsample: DESeq2
subsample: edgeR
subsample: limma-voom
|LFC| ≥ 3
|LFC| 2–3
|LFC| 1–2
|LFC| 0.2–1
5 10 15 25 50 75 100 150 200 300 500 600

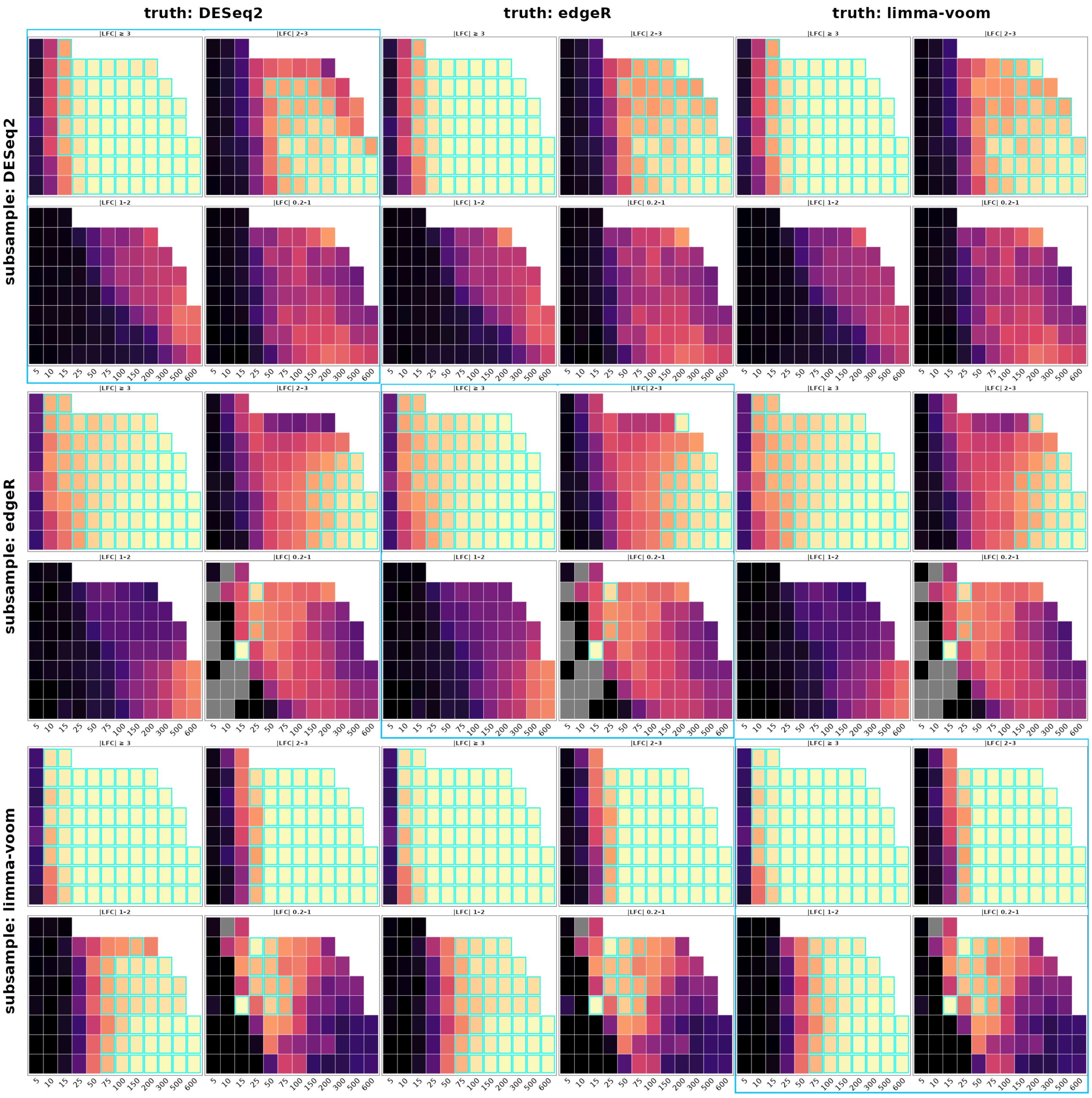
truth: DESeq2
truth: edgeR
truth: limma-voom
subsample: DESeq2
subsample: edgeR
subsample: limma-voom
|LFC| ≥ 3
|LFC| 2-3
|LFC| 1-2
|LFC| 0.2-1
5 10 15 25 50 75 100 150 200 300 500 600

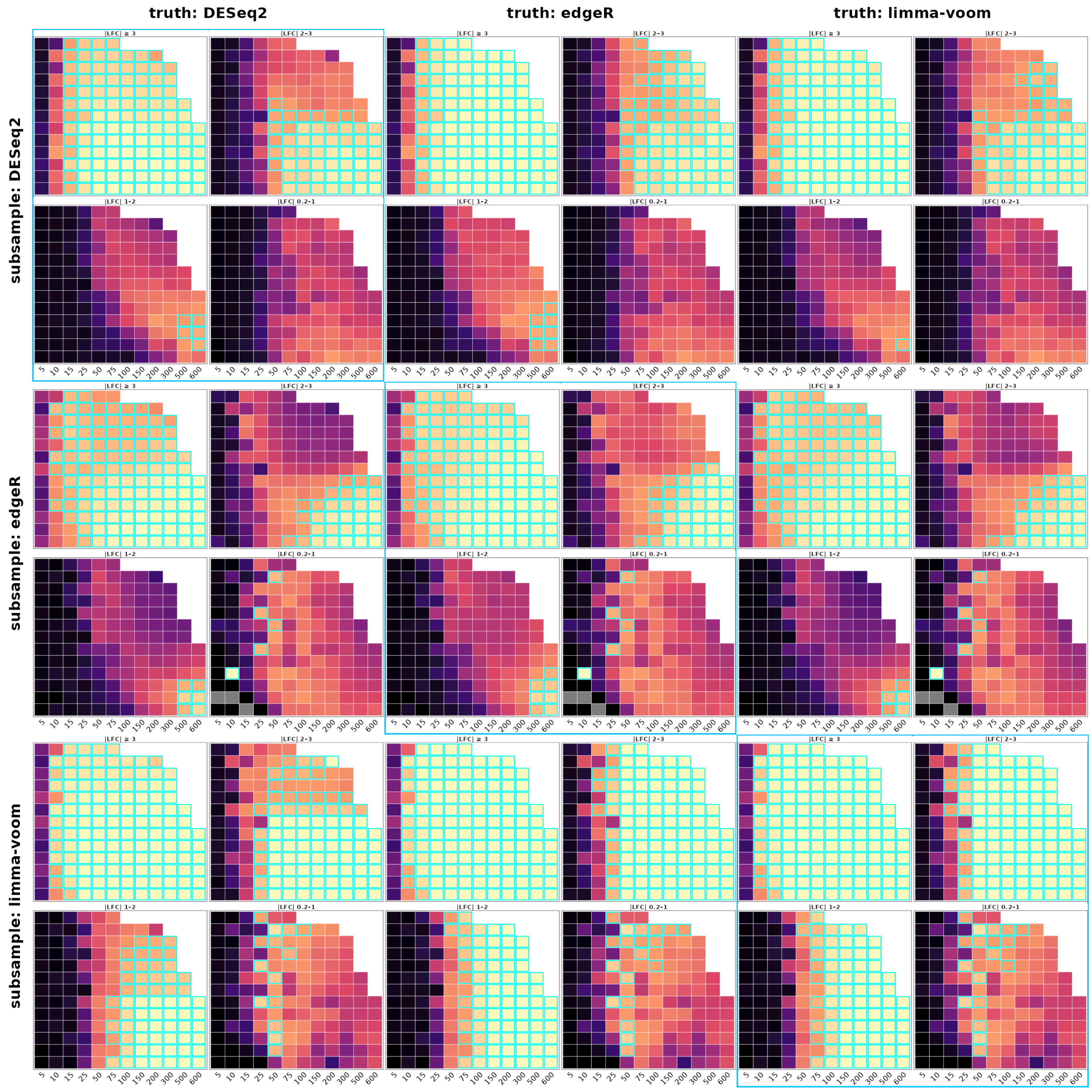


Figure 5: Cross-method precision grids (cells-based mode) for astrocytes (left), microglia (centre), and oligodendrocytes (right). Rows are the truth-defining DE method; columns are the downsampling DE method; panels show precision by LFC bin.

### Supplementary Fig. 5. Cross-method precision grids (counts-based mode)

Same layout as Supplementary Fig. 4, counts-based sampling mode.

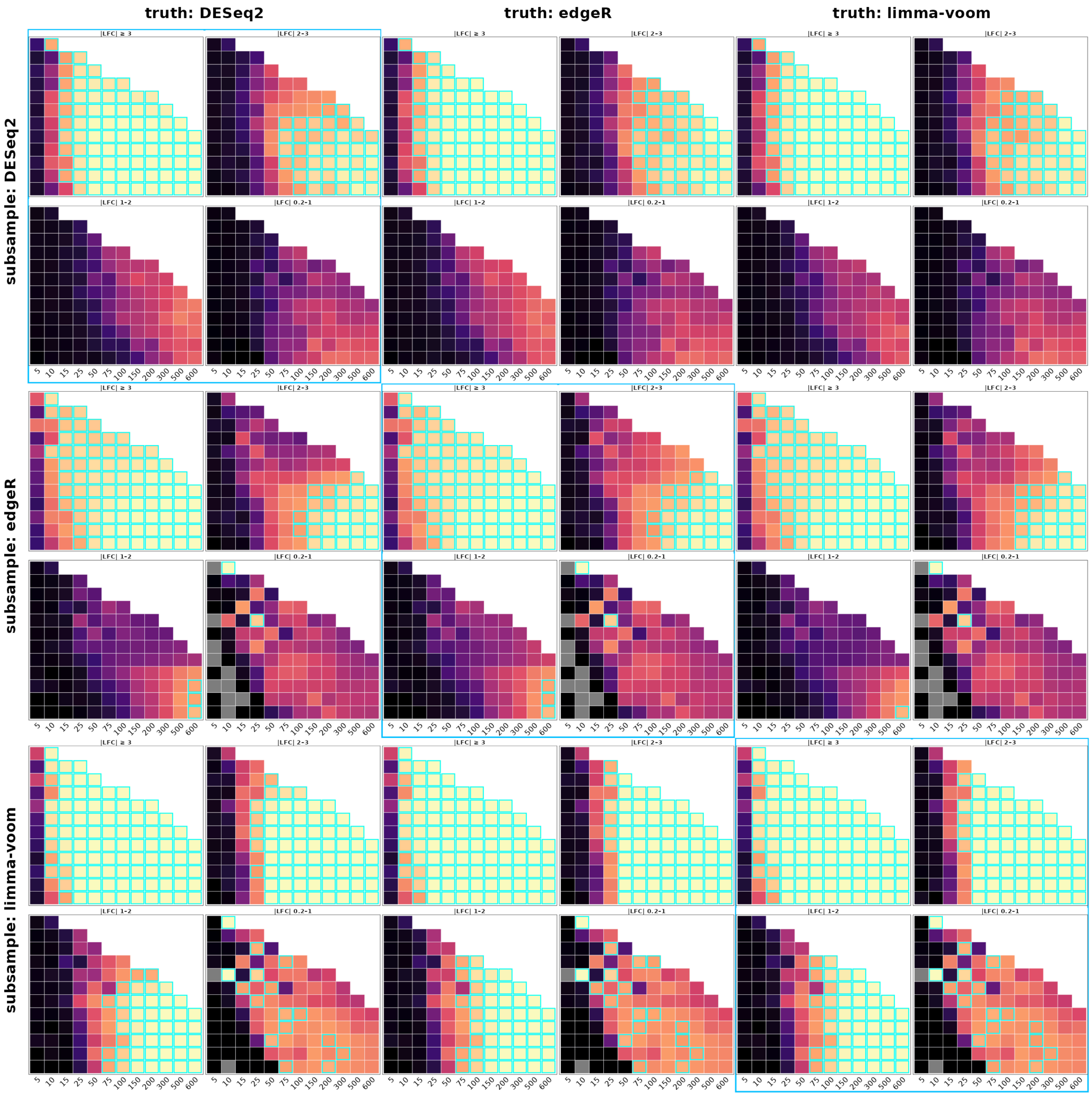
truth: DESeq2
truth: edgeR
truth: limma-voom
subsample: DESeq2
subsample: edgeR
subsample: limma-voom
|LFC| ≥ 3
|LFC| 2–3
|LFC| 1–2
|LFC| 0.2–1
5 10 15 25 50 75 100 150 200 300 500 600

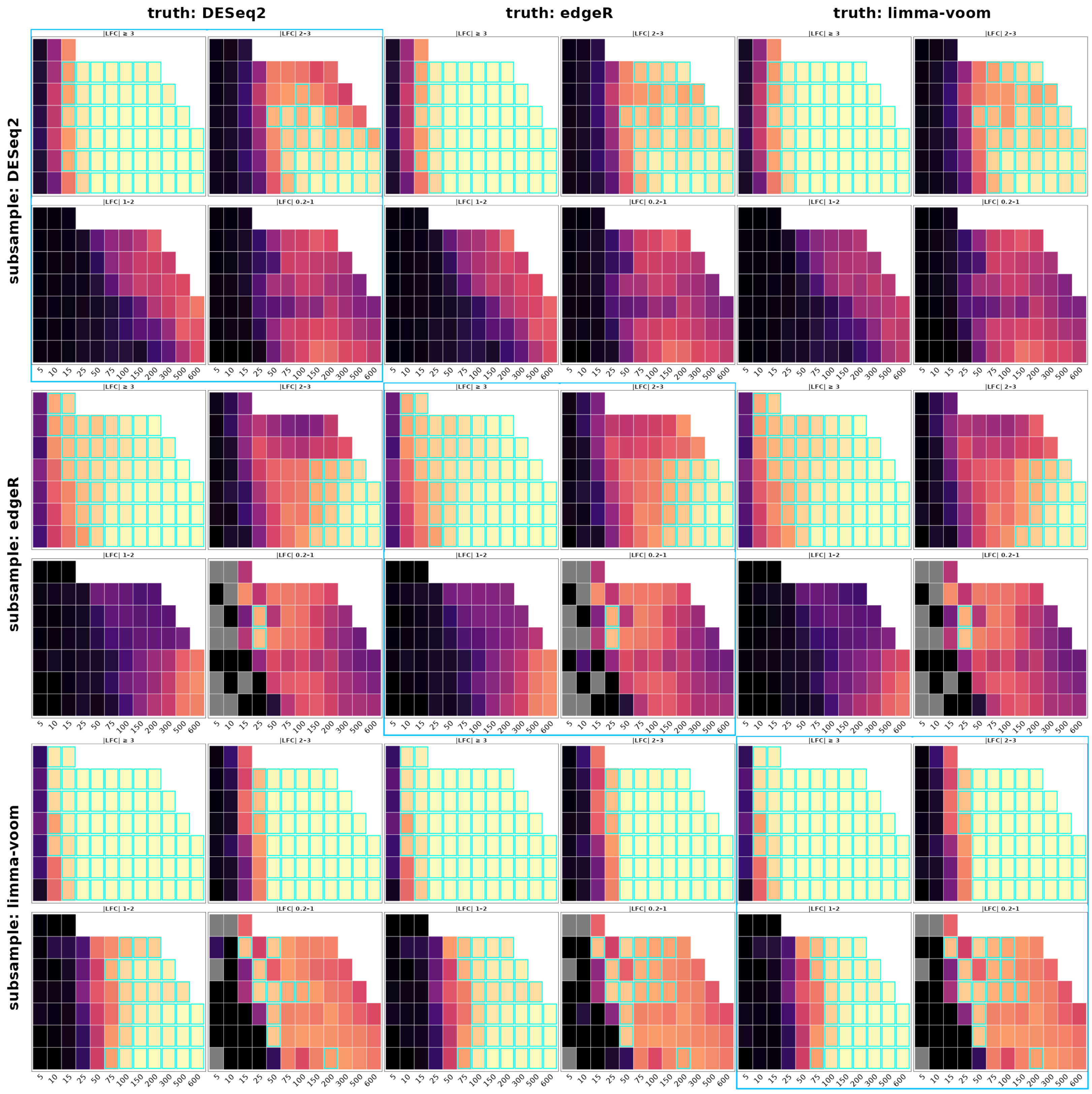
truth: DESeq2
truth: edgeR
truth: limma-voom
subsample: DESeq2
subsample: edgeR
subsample: limma-voom
|LFC| ≥ 3
|LFC| 2-3
|LFC| 1-2
|LFC| 0.2-1
5 10 15 25 50 75 100 150 200 300 500 600

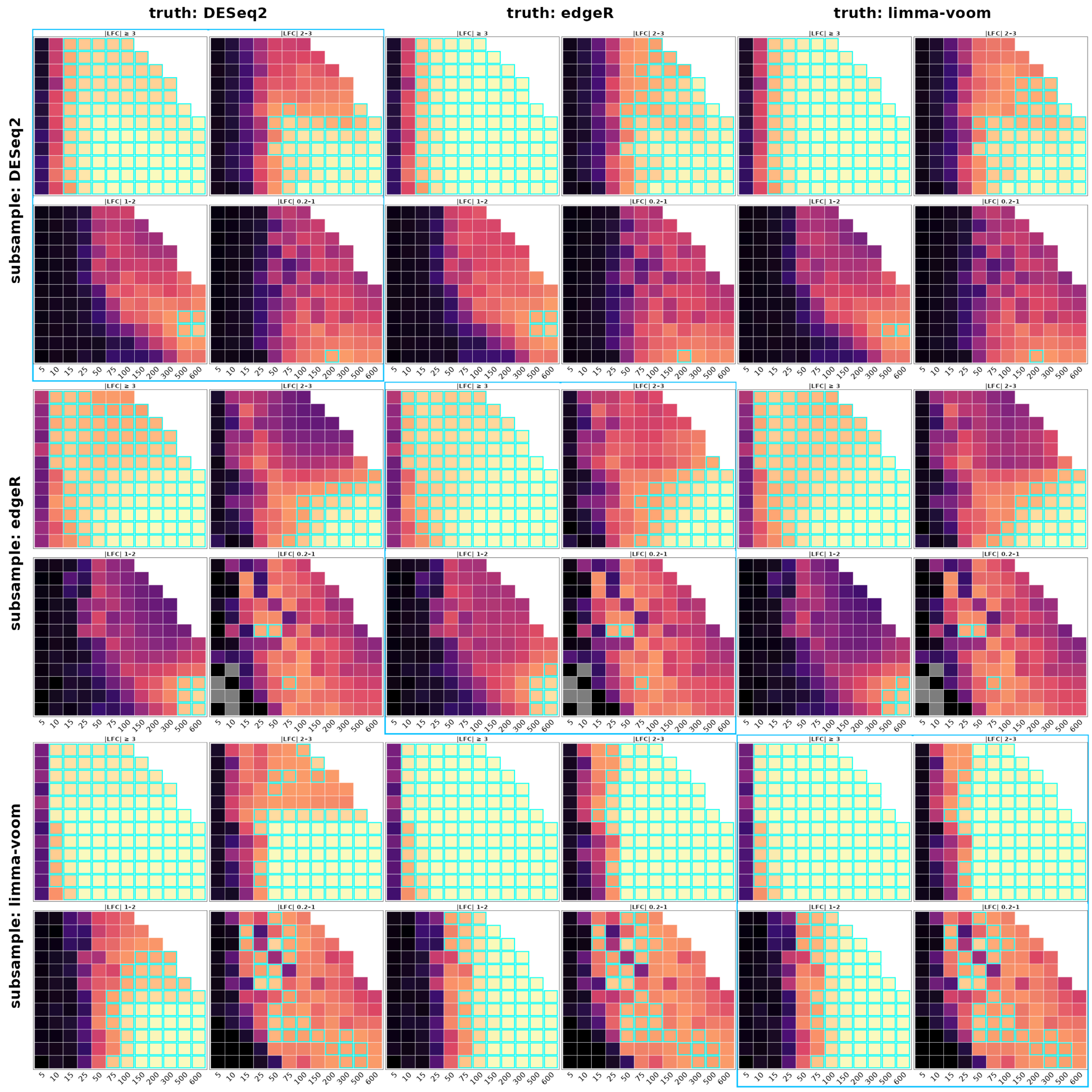

Figure 6: Cross-method precision grids (counts-based mode) for astrocytes (left), microglia (centre), and oligodendrocytes (right).

### Supplementary Figures: per-method downsampling grids

The following figures decompose the method-pooled Cartesian downsampling results (Supplementary Figs. 24-26) by the three differential-expression methods, showing DESeq2, edgeR, limma-voom, and the pooled all-methods result side by side (2×2 panel per figure) for a single cell type. They complement the cross-method comparison grids (Supplementary Figs. 2-5), which instead cross each method's truth set against each method's detection step. For every figure the cells-based sampling mode is shown on top and the counts-based mode below. Three views are provided per cell type and metric: the continuous semi-log surface across the full donor grid, and the recall/precision-versus-mean-expression curves within the tiny (|LFC| 0.2-1.0) and small (|LFC| 1.0-2.0) effect-size bins that the paper identifies as the underpowered regime.

#### Supplementary Fig. 6. Astrocytes: recall surface across the downsampling grid, by DE method

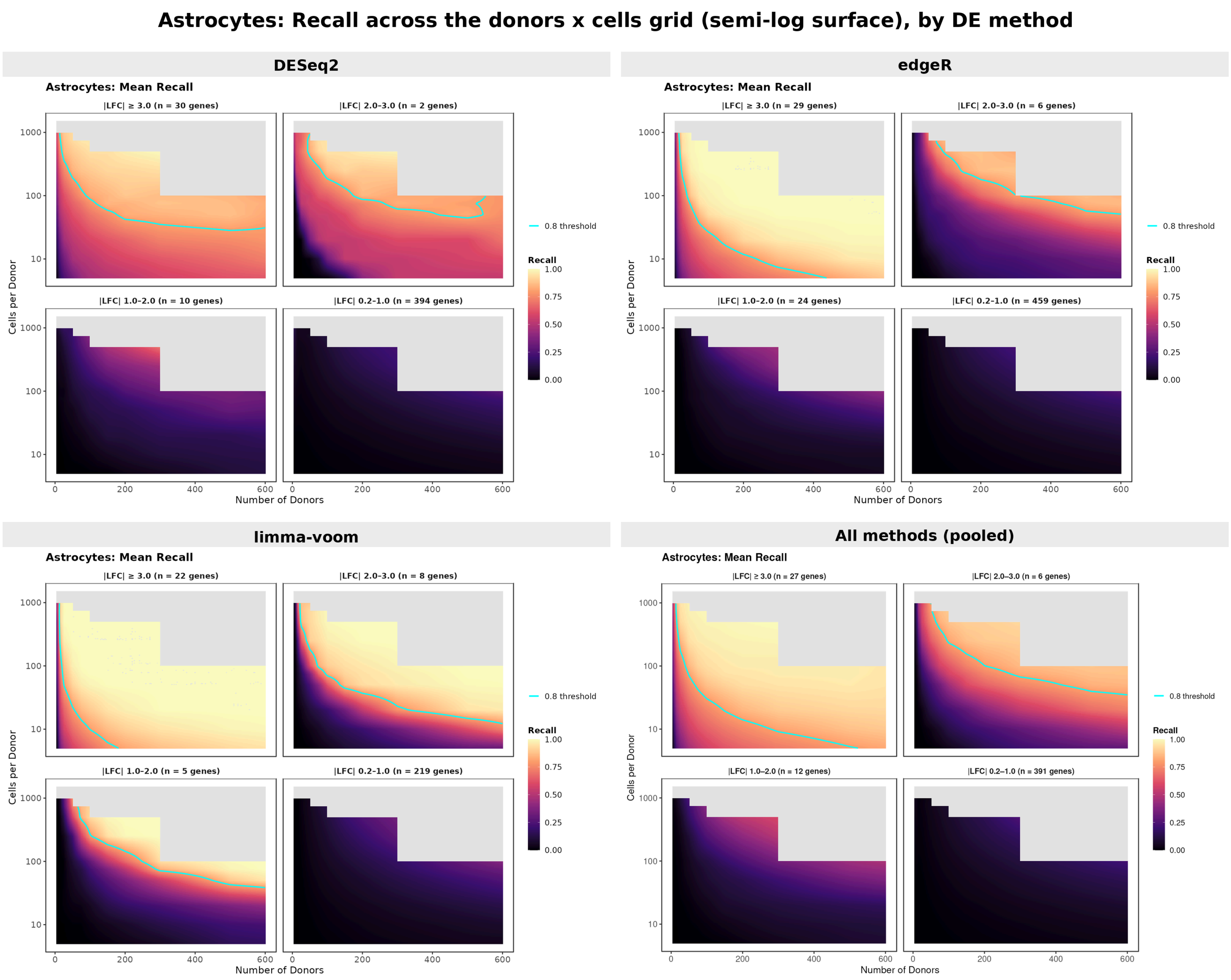


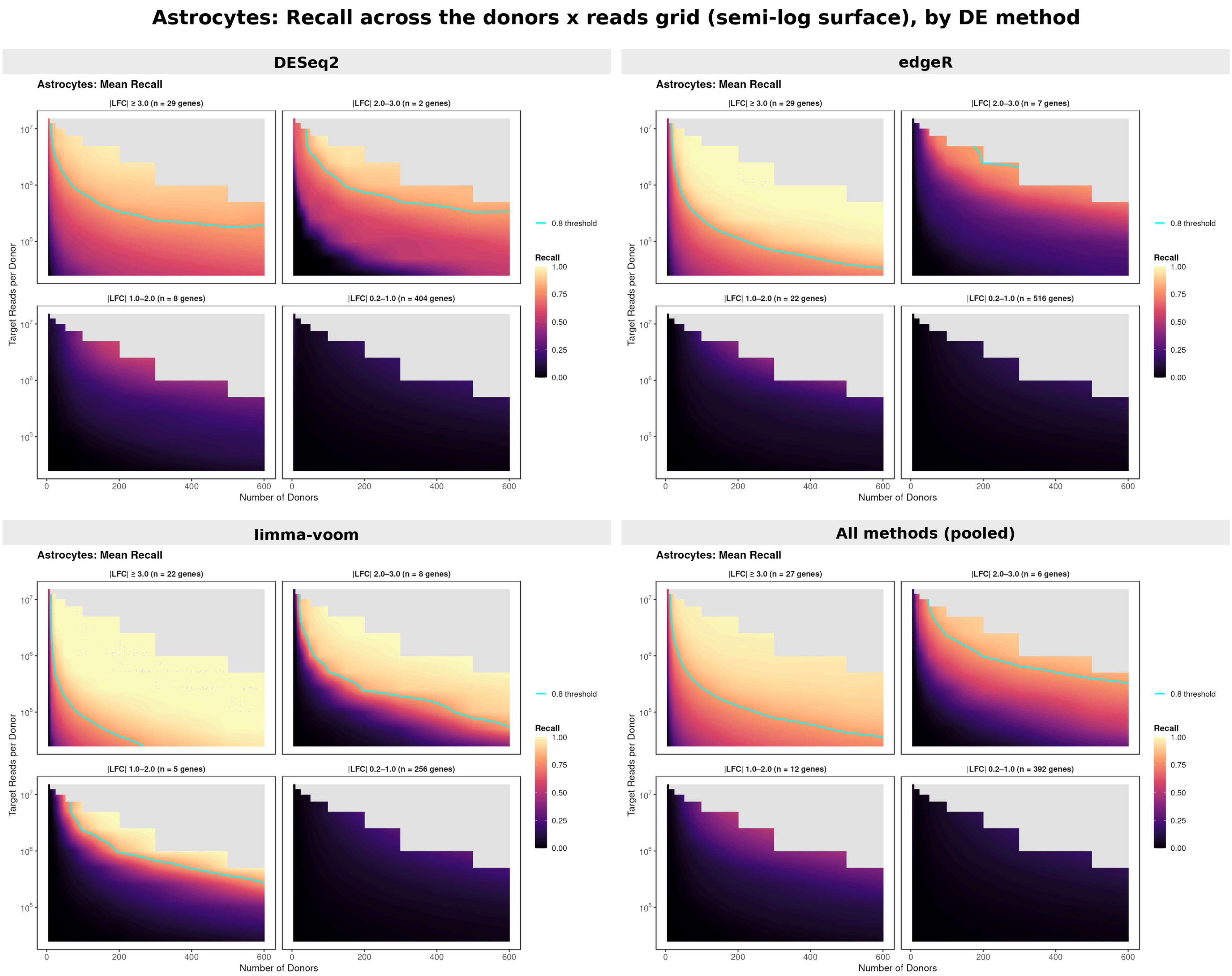


Figure 7: Astrocyte continuous recall surfaces (semi-log scale) split by DE method (DESeq2, edgeR, limma-voom, all-methods pooled). Cells-based sampling mode (top) and counts-based sampling mode (bottom). Each panel tiles the four methods in a 2×2 grid.

### Supplementary Fig. 7. Astrocytes: precision surface across the downsampling grid, by DE method

**Astrocytes: Precision across the donors x cells grid (semi-log surface), by DE method**

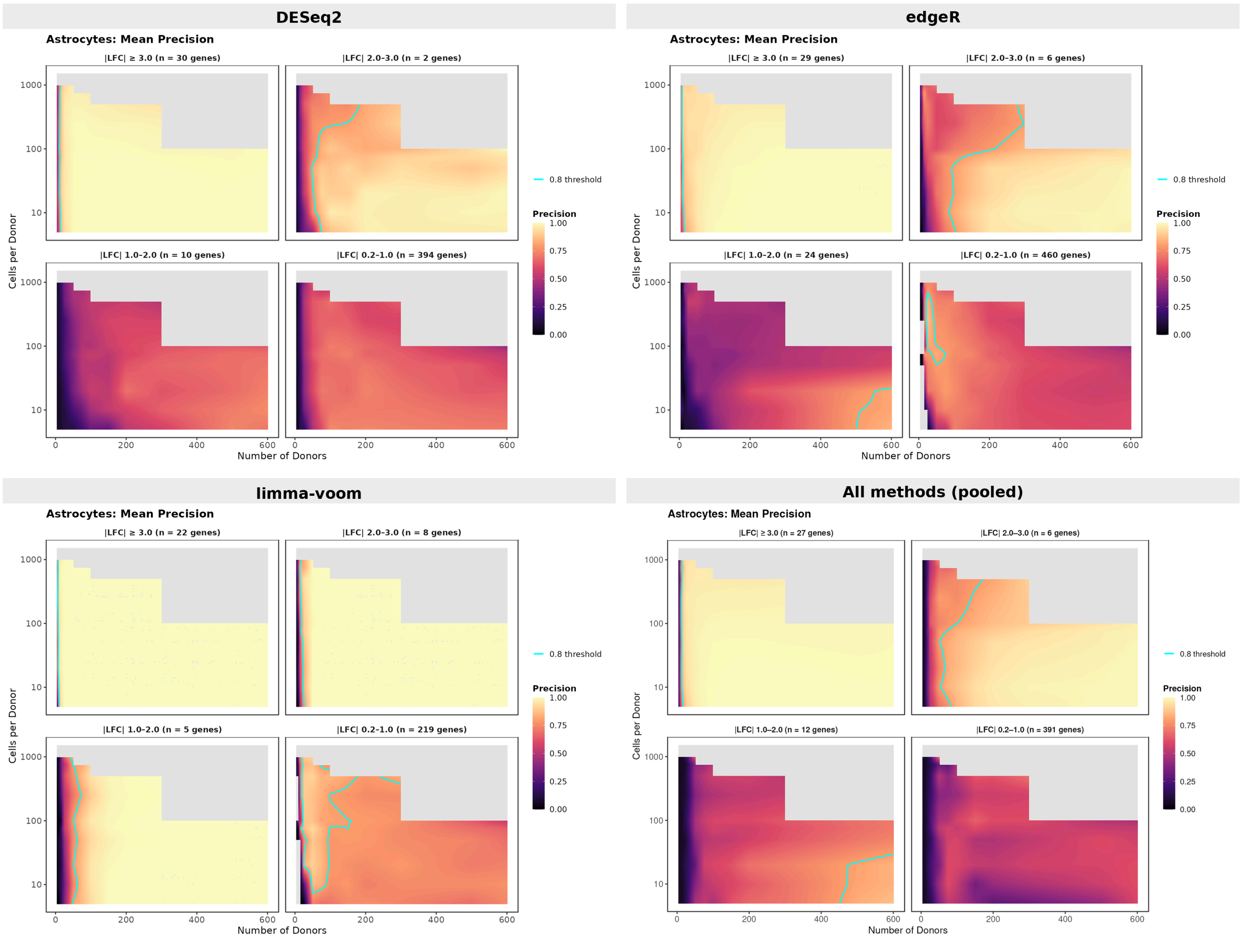


**Astrocytes: Precision across the donors x reads grid (semi-log surface), by DE method**

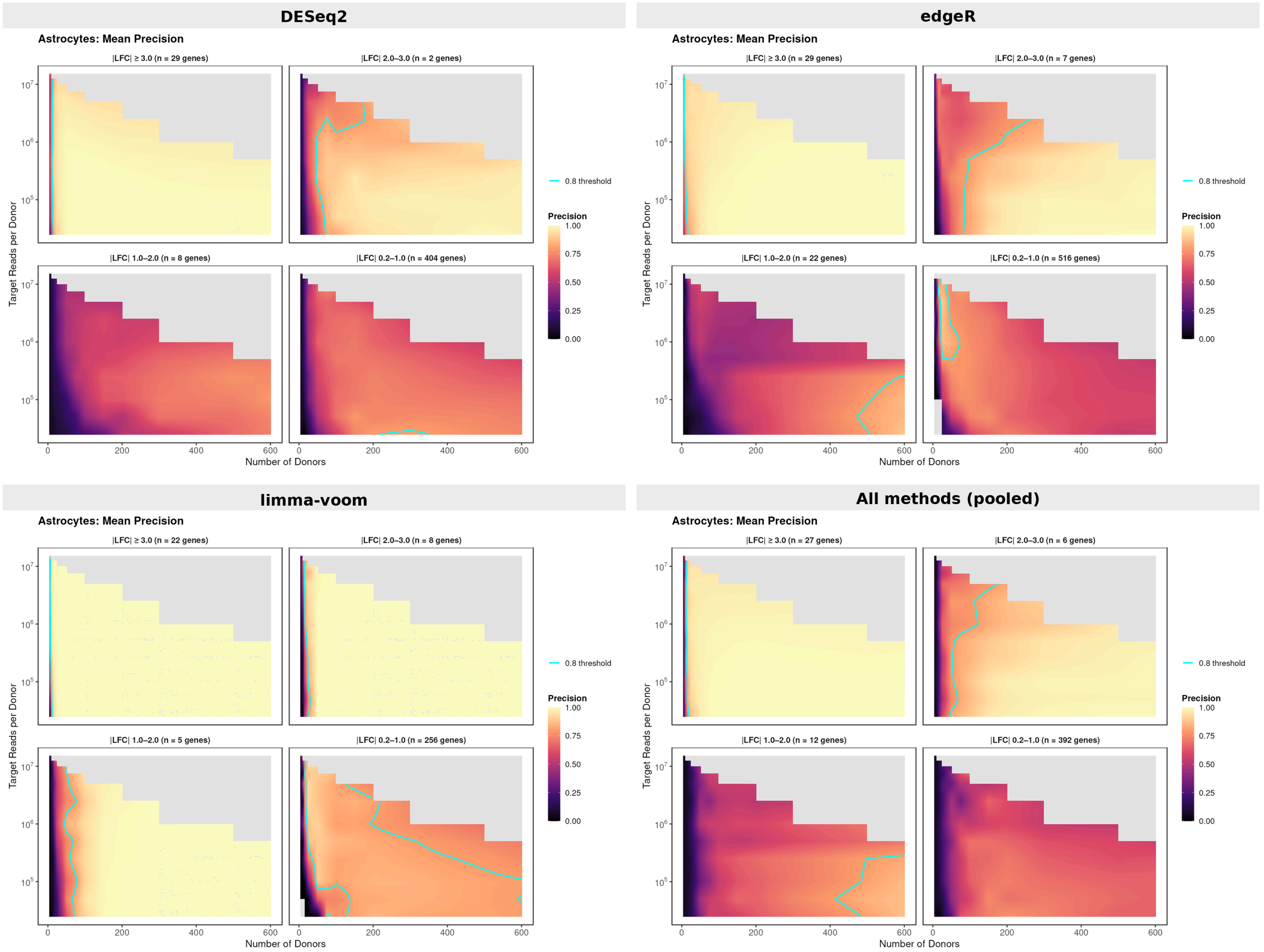

Figure 8: Astrocyte continuous precision surfaces (semi-log scale) split by DE method. Cells-based sampling mode (top) and counts-based sampling mode (bottom).

### Supplementary Fig. 8. Astrocytes: tiny-effect (|LFC| 0.2-1.0) recall versus mean expression, by DE method

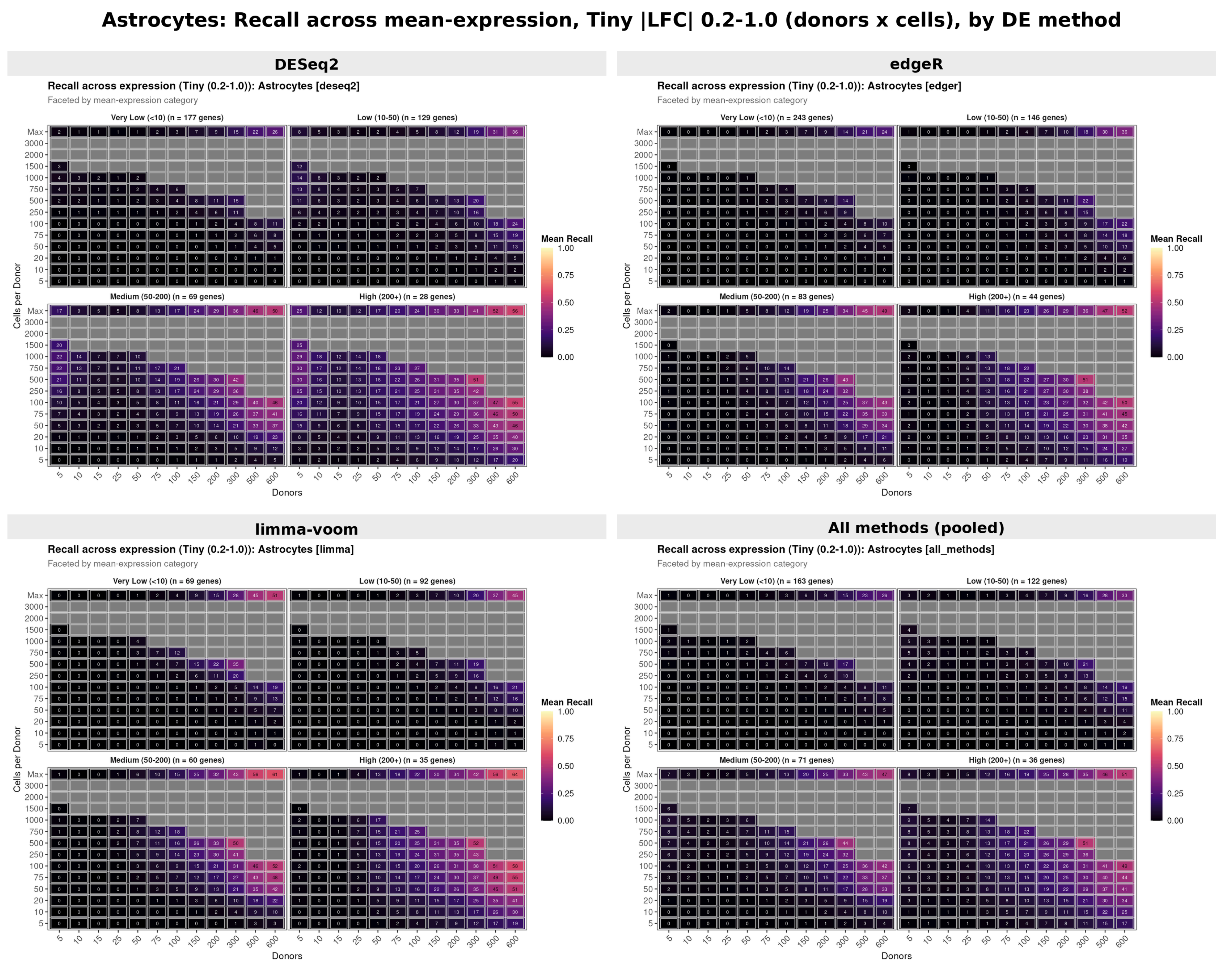


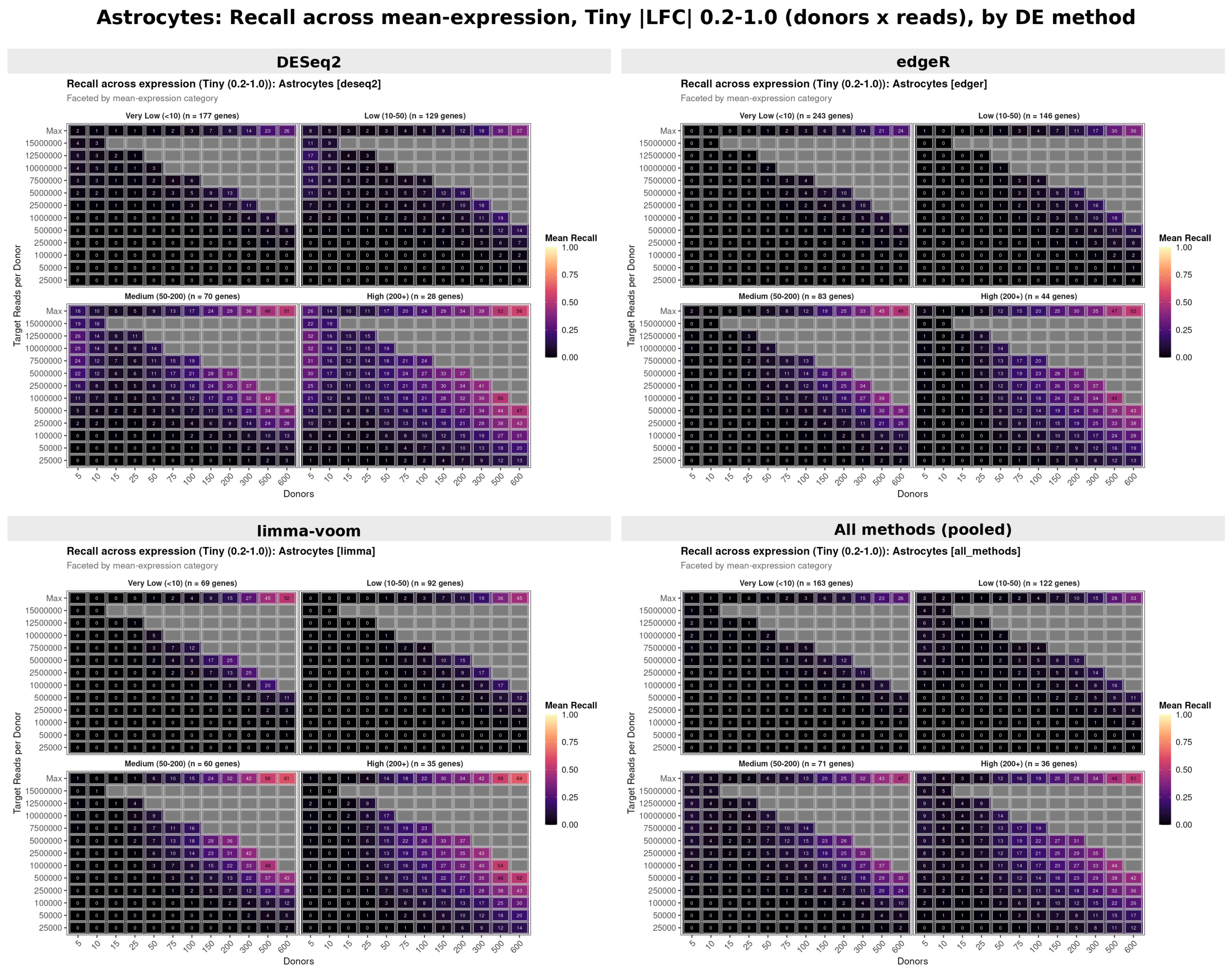


Figure 9: Astrocyte recall across mean expression within the tiny effect-size bin (|LFC| 0.2-1.0), split by DE method. Cells-based sampling mode (top) and counts-based sampling mode (bottom).

### Supplementary Fig. 9. Astrocytes: tiny-effect (|LFC| 0.2-1.0) precision versus mean expression, by DE method

**Astrocytes: Precision across mean-expression, Tiny |LFC| 0.2-1.0 (donors x cells), by DE method**

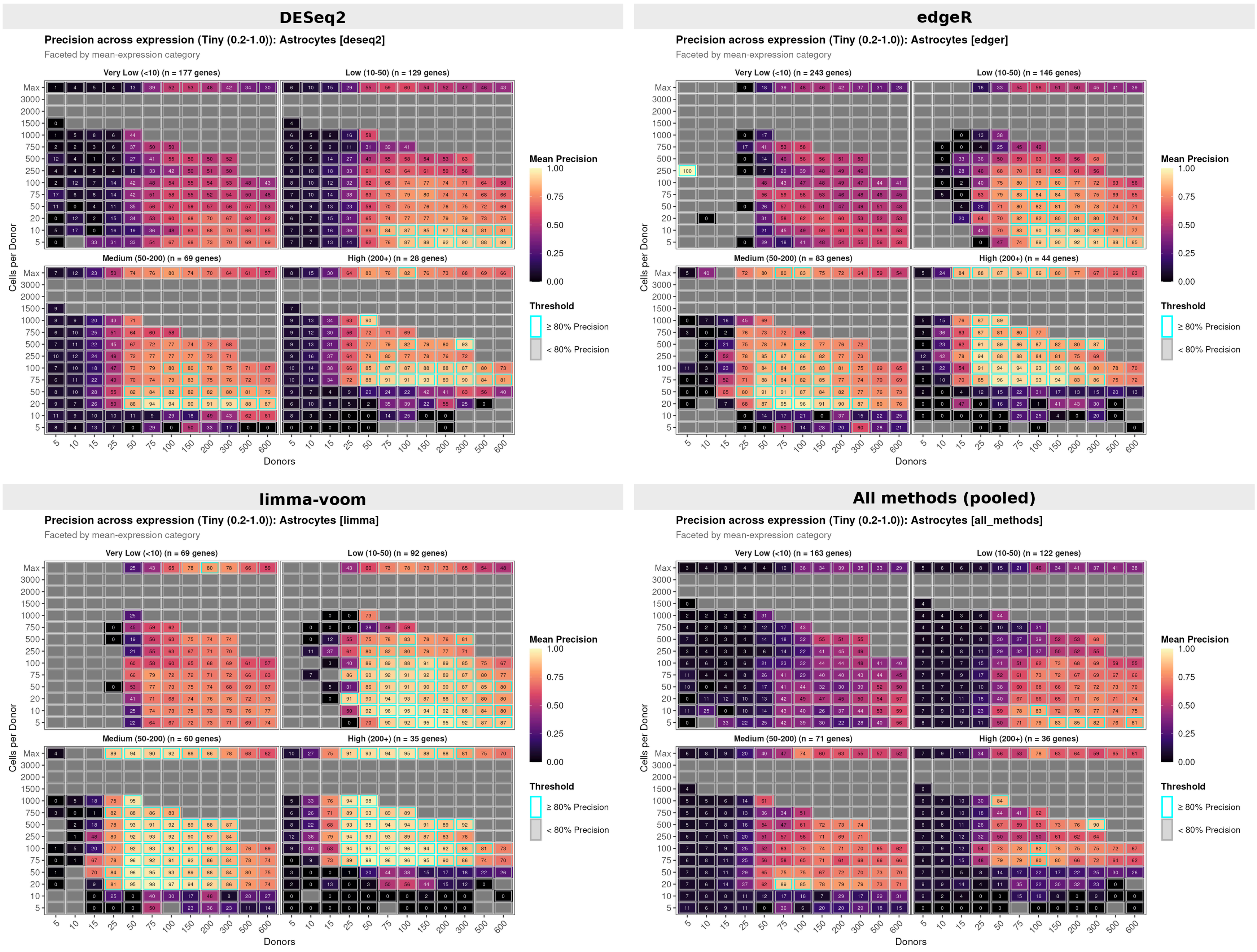


**Astrocytes: Precision across mean-expression, Tiny |LFC| 0.2-1.0 (donors x reads), by DE method**

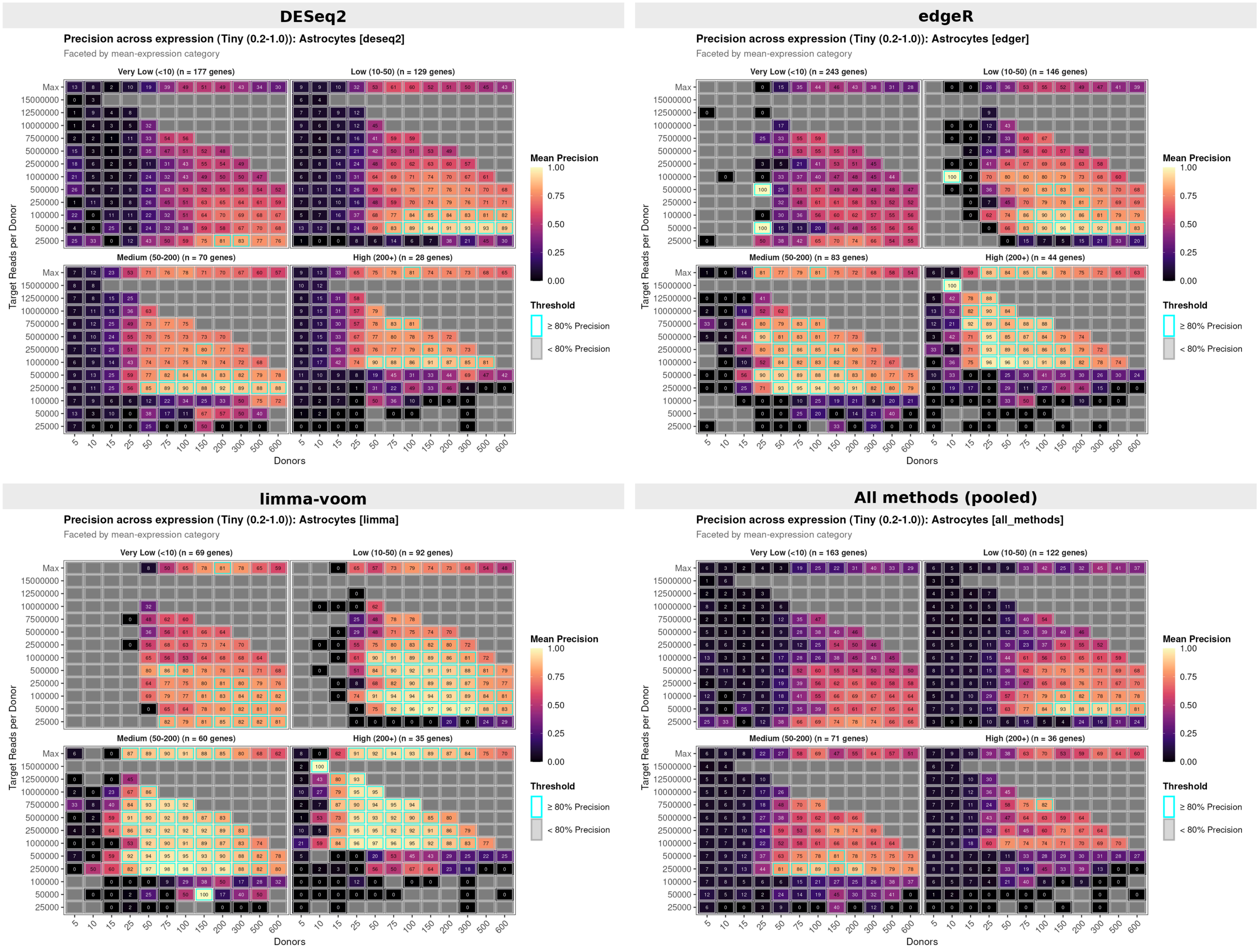

Figure 10: Astrocyte precision across mean expression within the tiny effect-size bin (|LFC| 0.2-1.0), split by DE method. Cells-based sampling mode (top) and counts-based sampling mode (bottom).

### Supplementary Fig. 10. Astrocytes: small-effect (|LFC| 1.0-2.0) recall versus mean expression, by DE method

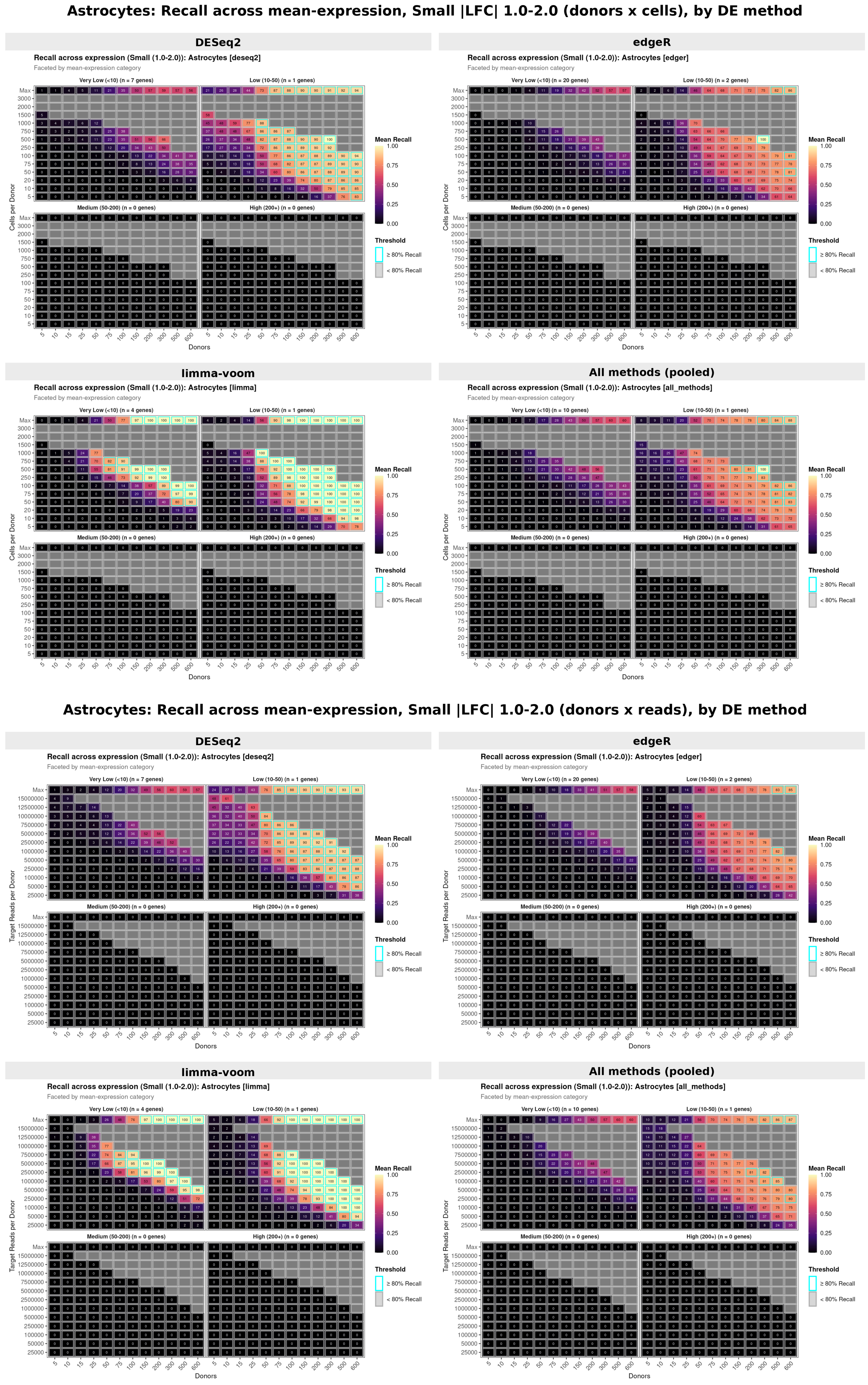


Figure 11: Astrocyte recall across mean expression within the small effect-size bin (|LFC| 1.0-2.0), split by DE method. Cells-based sampling mode (top) and counts-based sampling mode (bottom).

### Supplementary Fig. 11. Astrocytes: small-effect (|LFC| 1.0-2.0) precision versus mean expression, by DE method

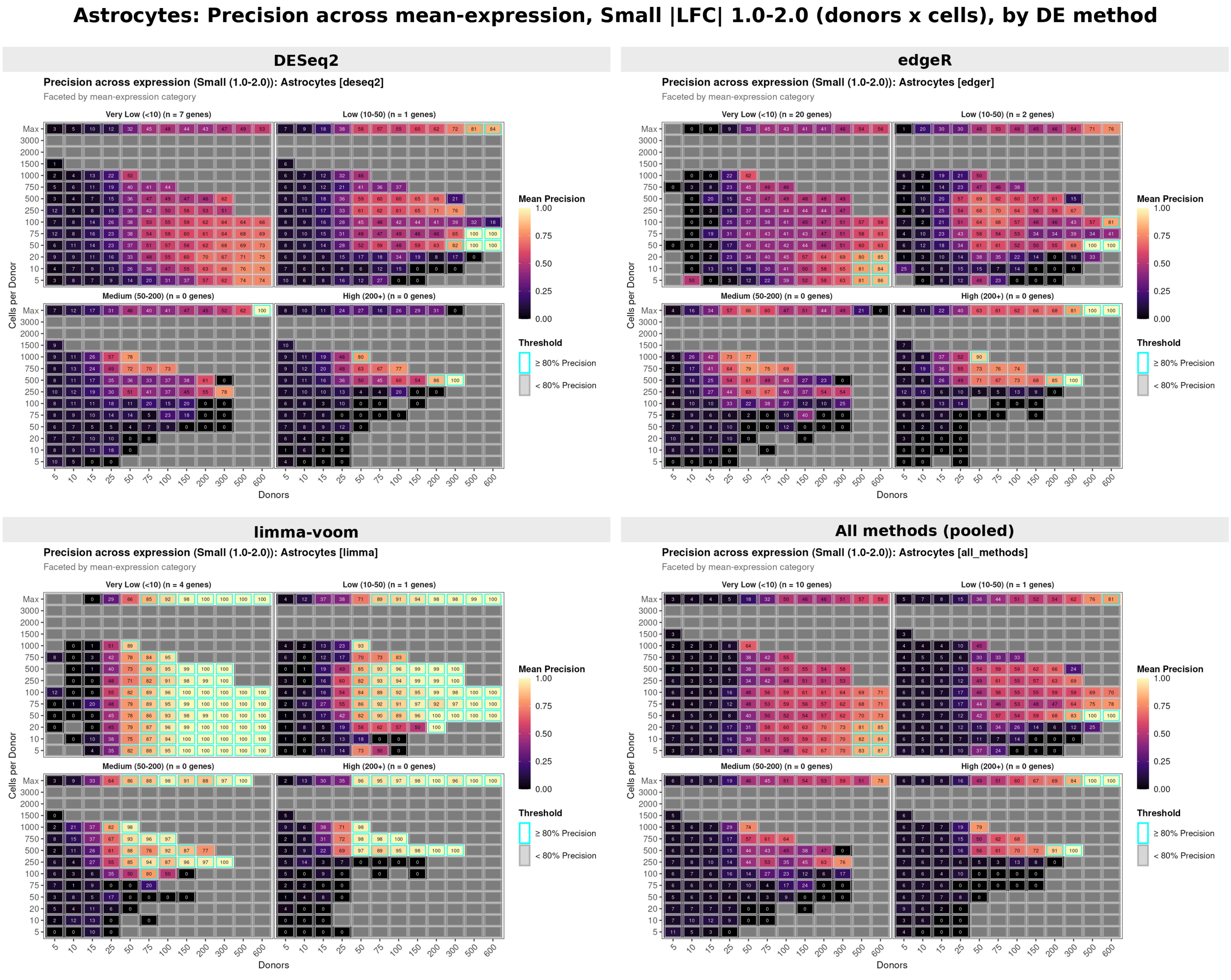


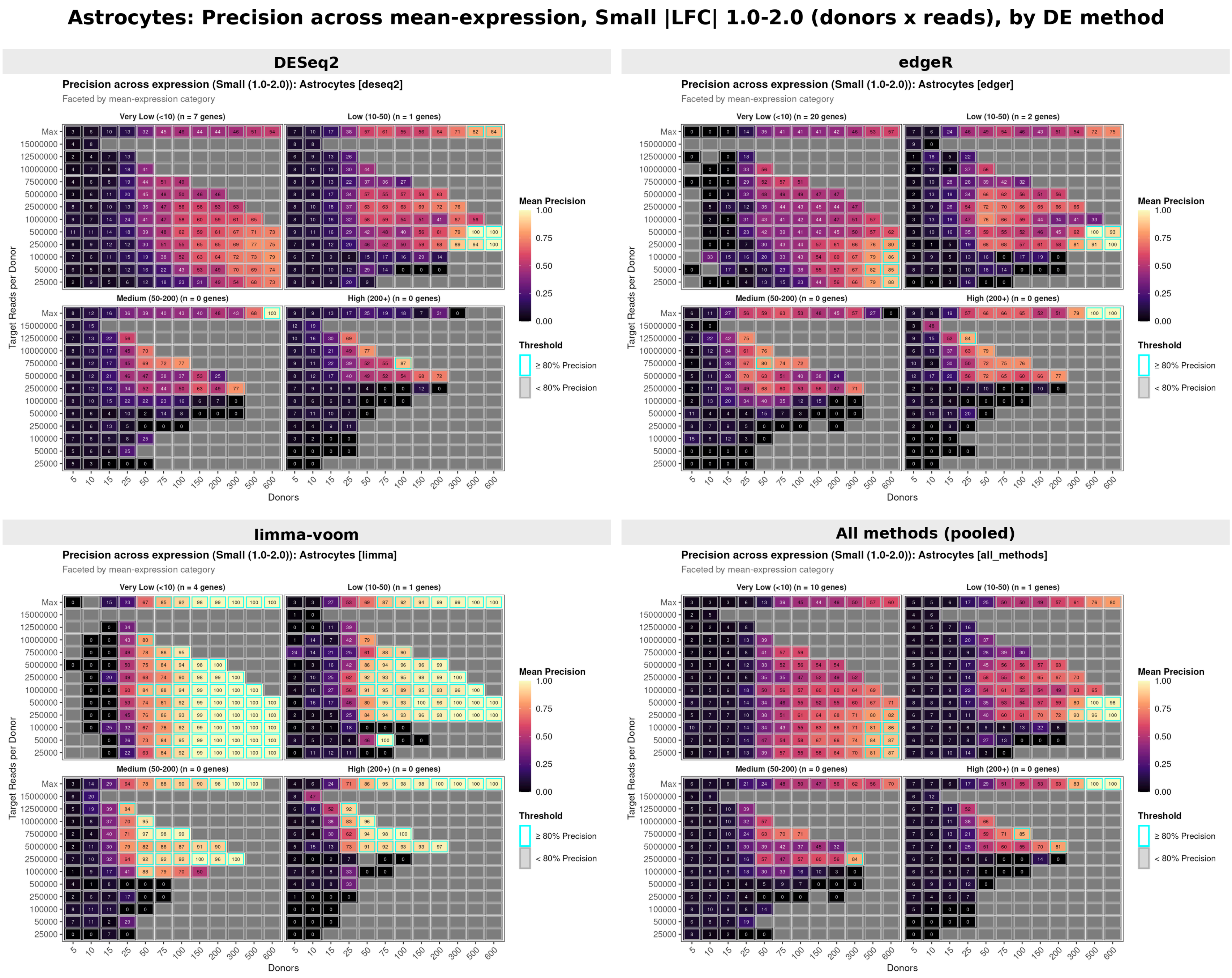

Figure 12: Astrocyte precision across mean expression within the small effect-size bin (|LFC| 1.0-2.0), split by DE method. Cells-based sampling mode (top) and counts-based sampling mode (bottom).

### Supplementary Fig. 12. Microglia: recall surface across the downsampling grid, by DE method

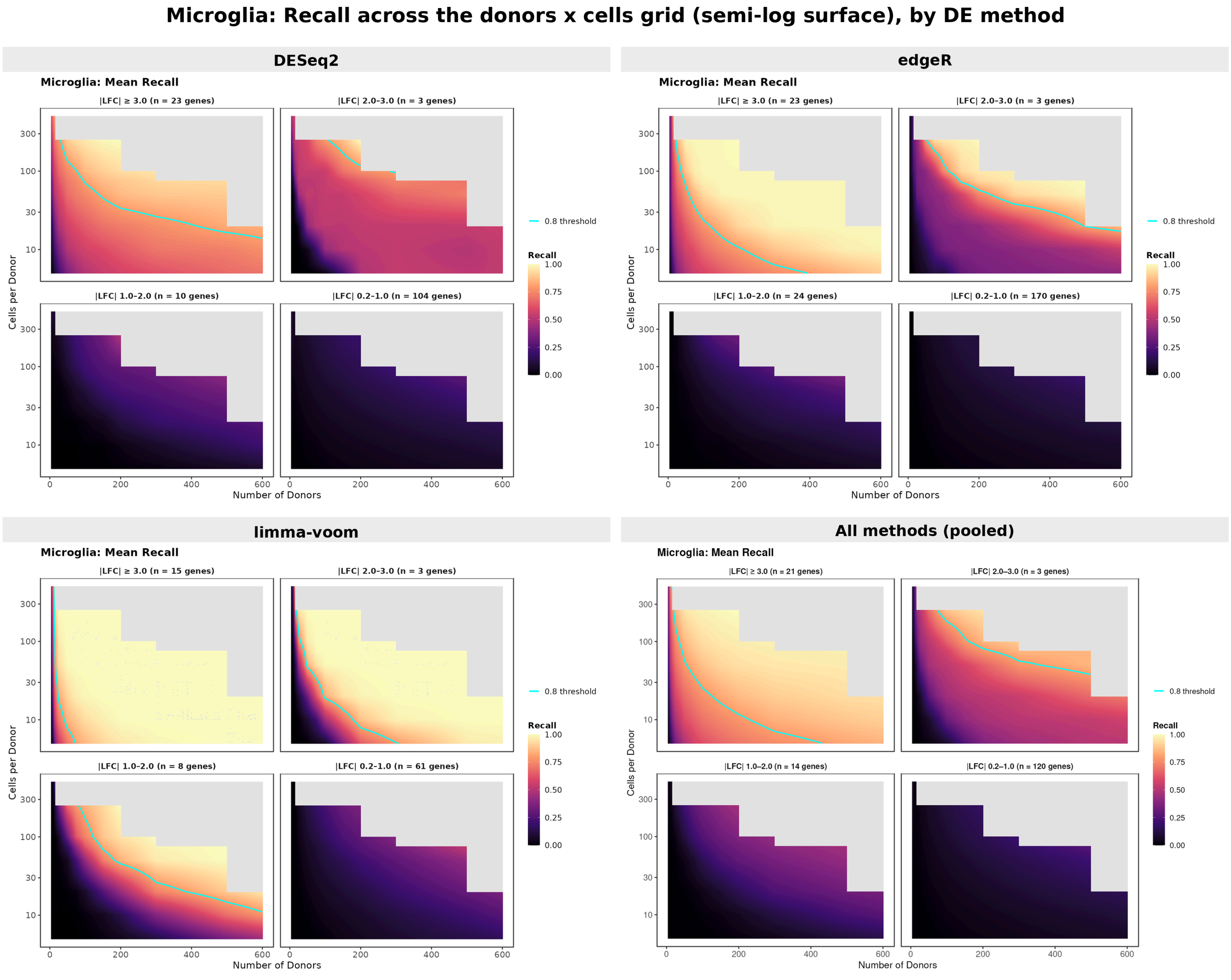


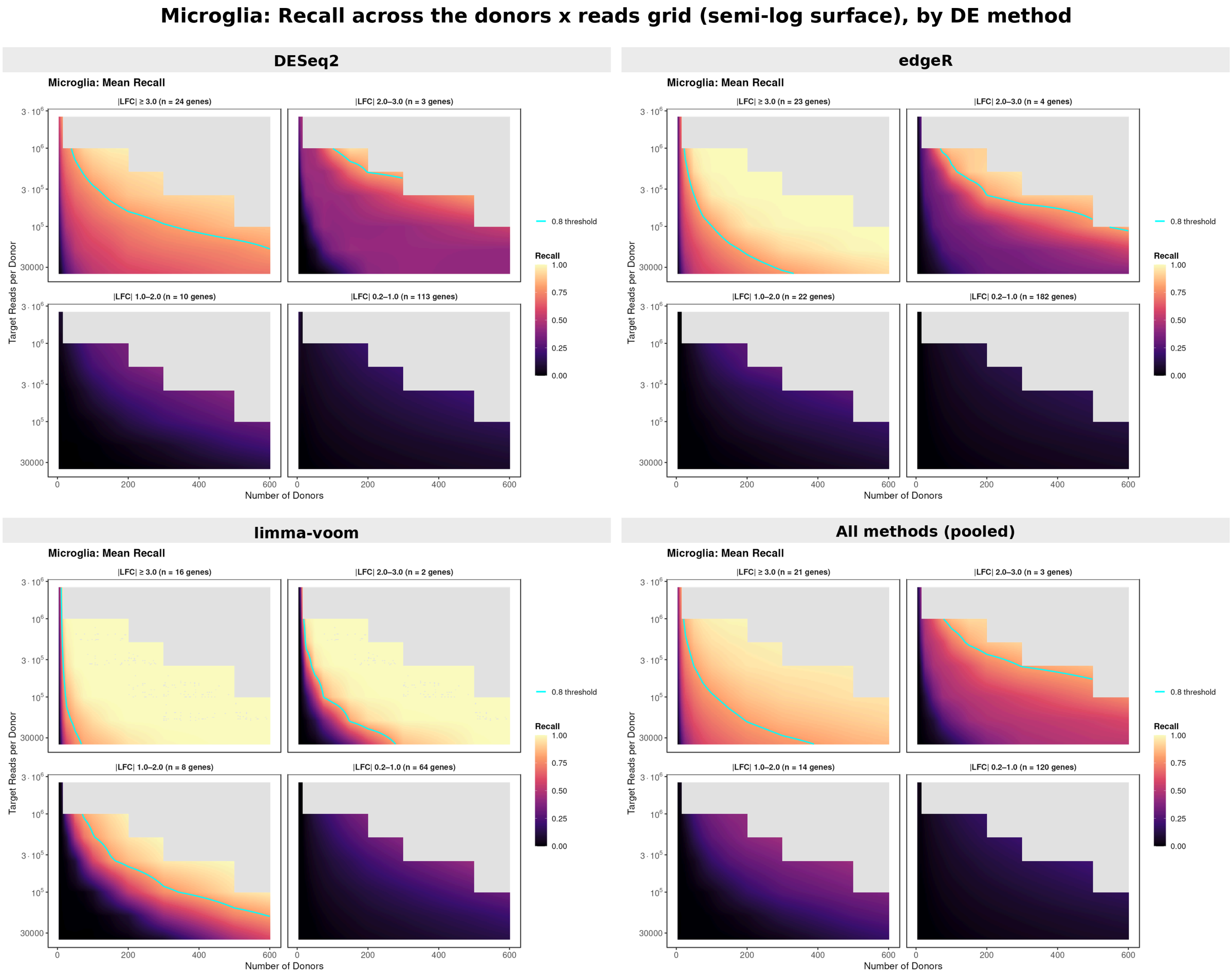


Figure 13: Microglia continuous recall surfaces (semi-log scale) split by DE method. Cells-based sampling mode (top) and counts-based sampling mode (bottom).

### Supplementary Fig. 13. Microglia: precision surface across the downsampling grid, by DE method

**Microglia: Precision across the donors x cells grid (semi-log surface), by DE method**

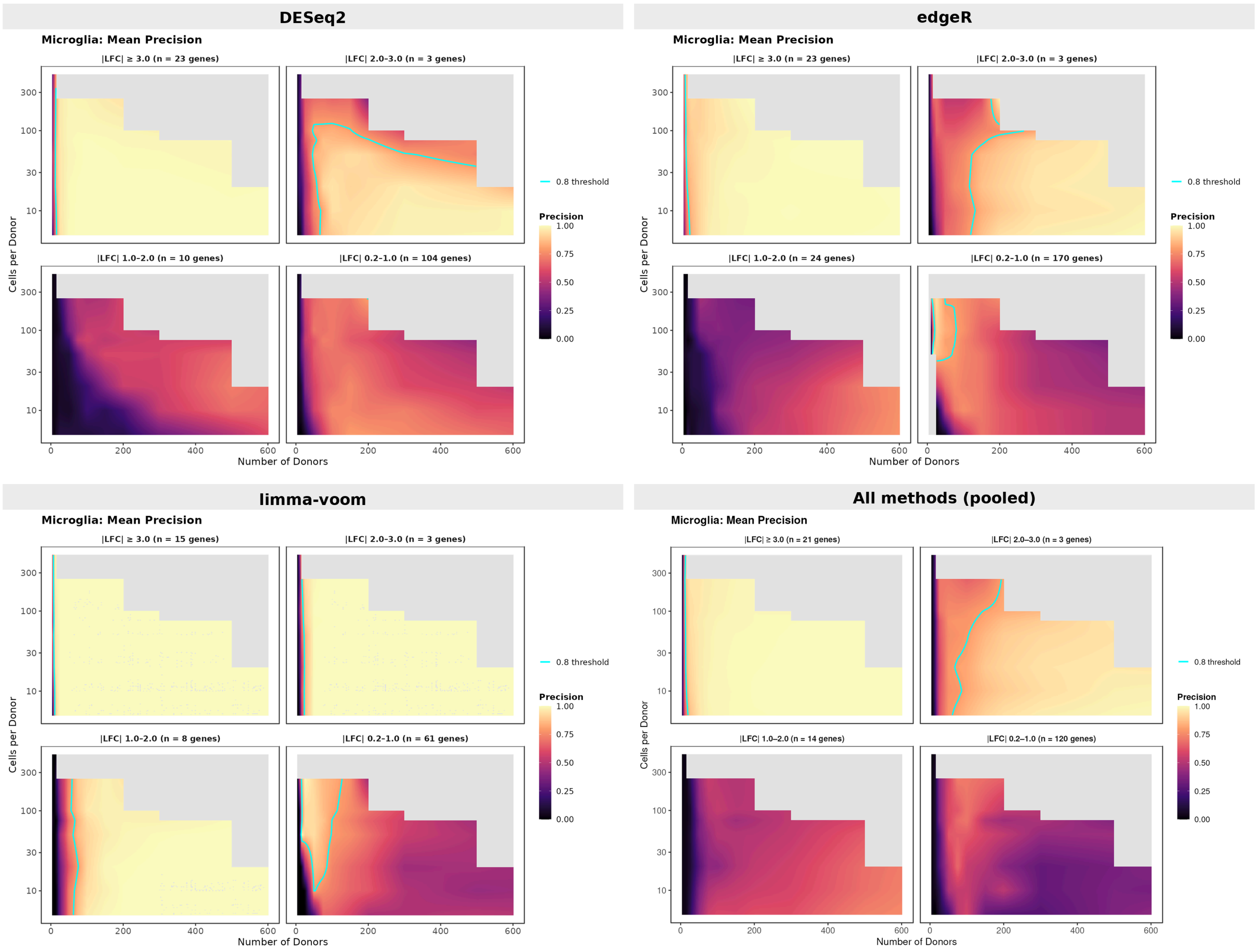


**Microglia: Precision across the donors x reads grid (semi-log surface), by DE method**

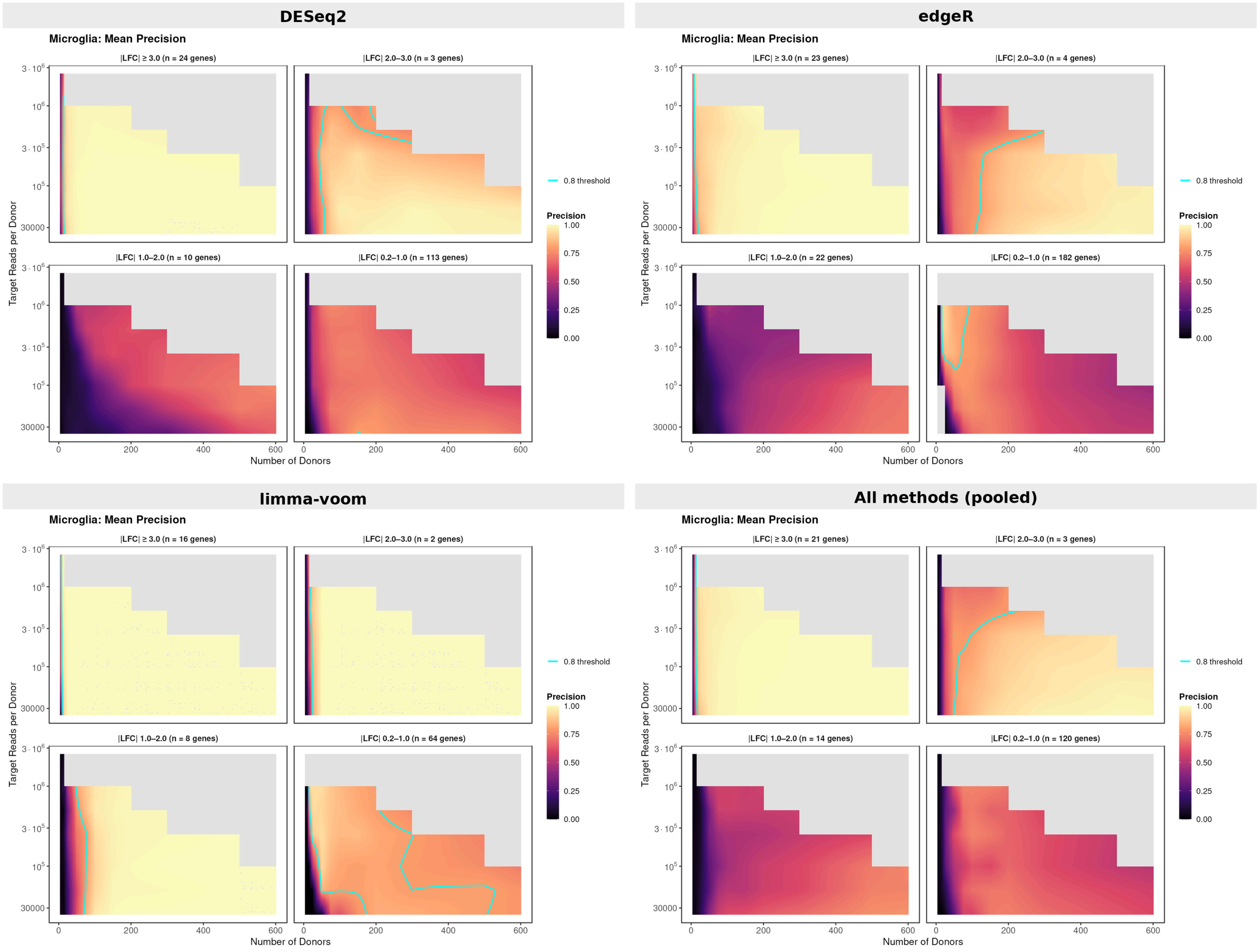

Figure 14: Microglia continuous precision surfaces (semi-log scale) split by DE method. Cells-based sampling mode (top) and counts-based sampling mode (bottom).

### Supplementary Fig. 14. Microglia: tiny-effect (|LFC| 0.2-1.0) recall versus mean expression, by DE method

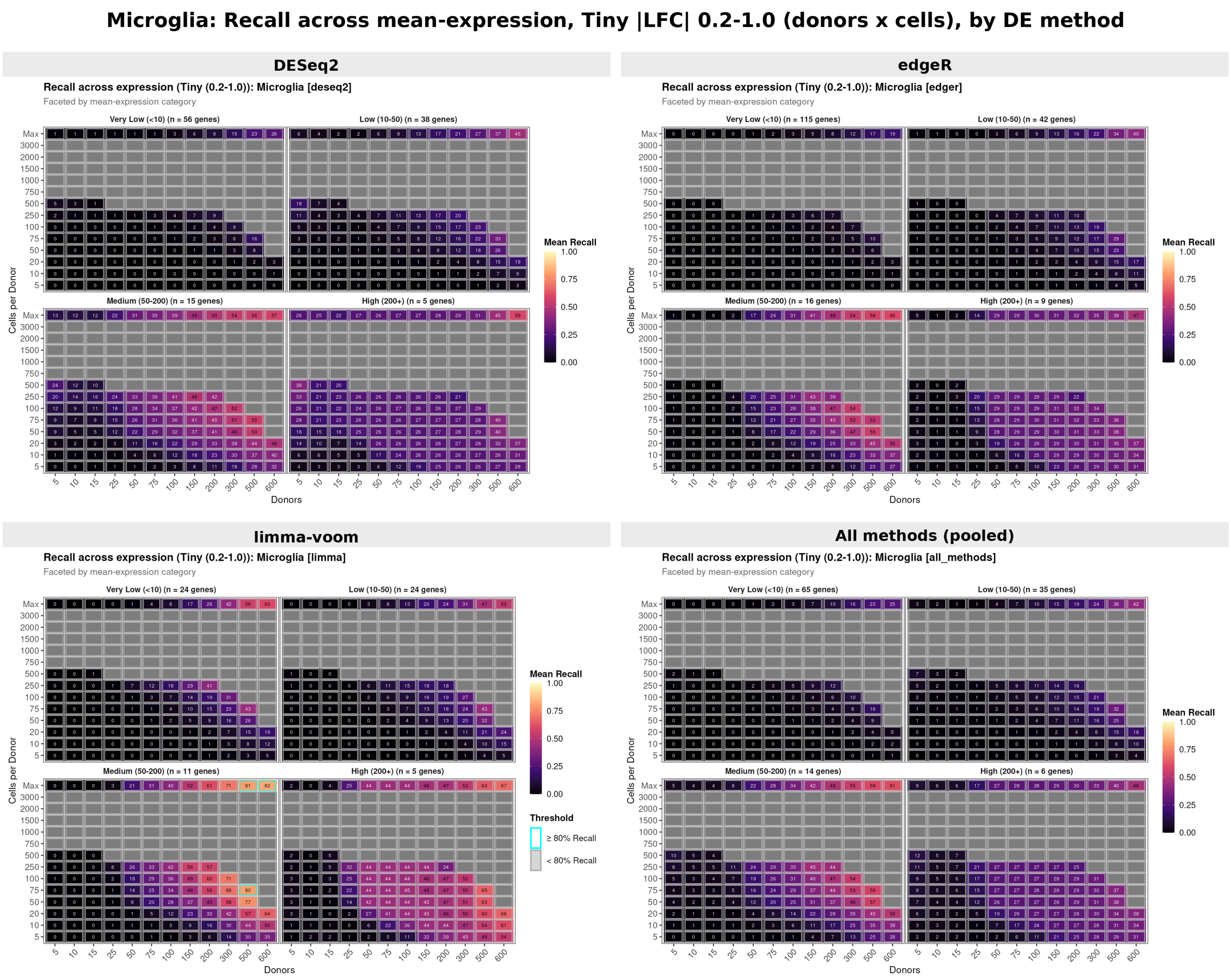


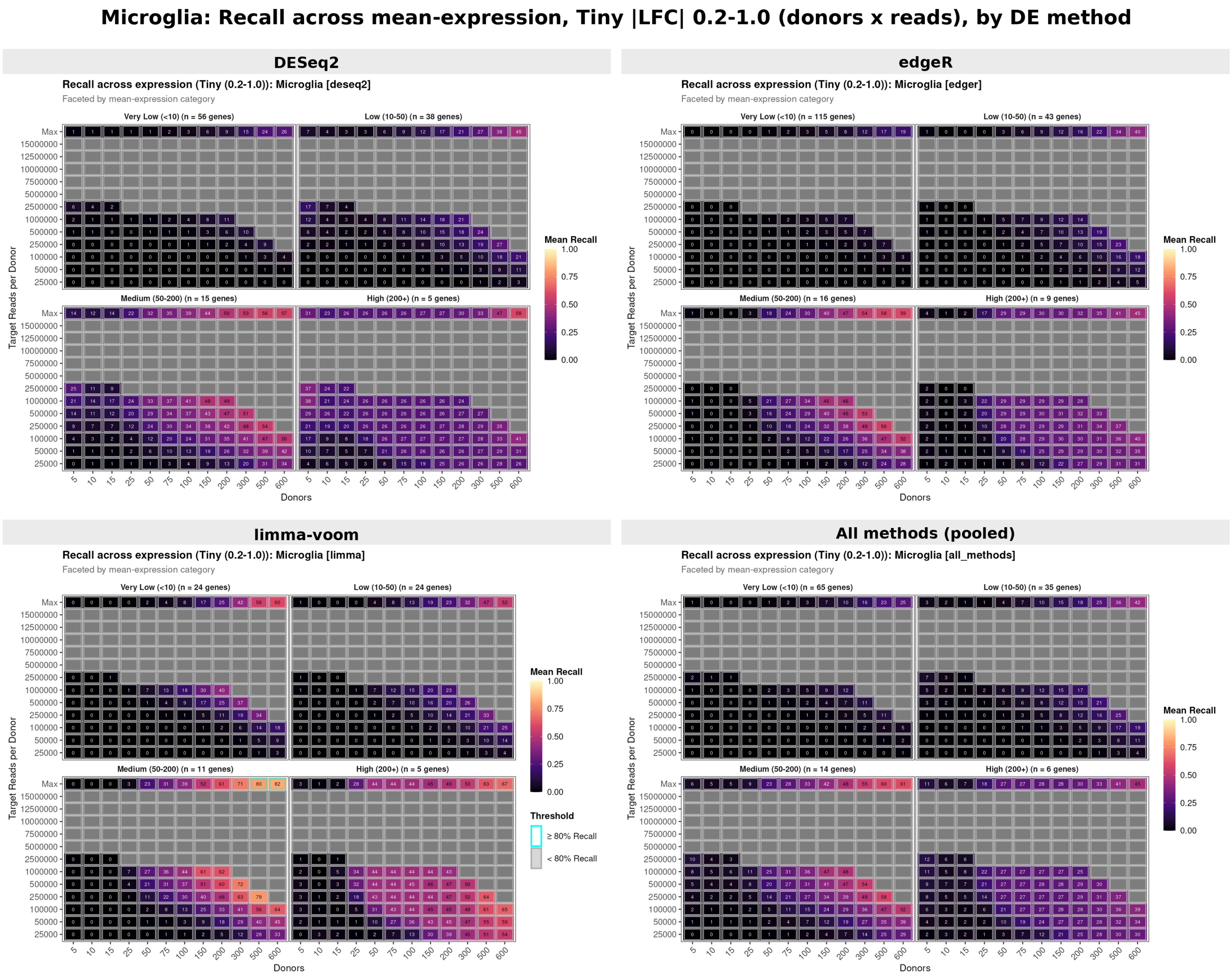


Figure 15: Microglia recall across mean expression within the tiny effect-size bin (|LFC| 0.2-1.0), split by DE method. Cells-based sampling mode (top) and counts-based sampling mode (bottom).

### Supplementary Fig. 15. Microglia: tiny-effect (|LFC| 0.2-1.0) precision versus mean expression, by DE method

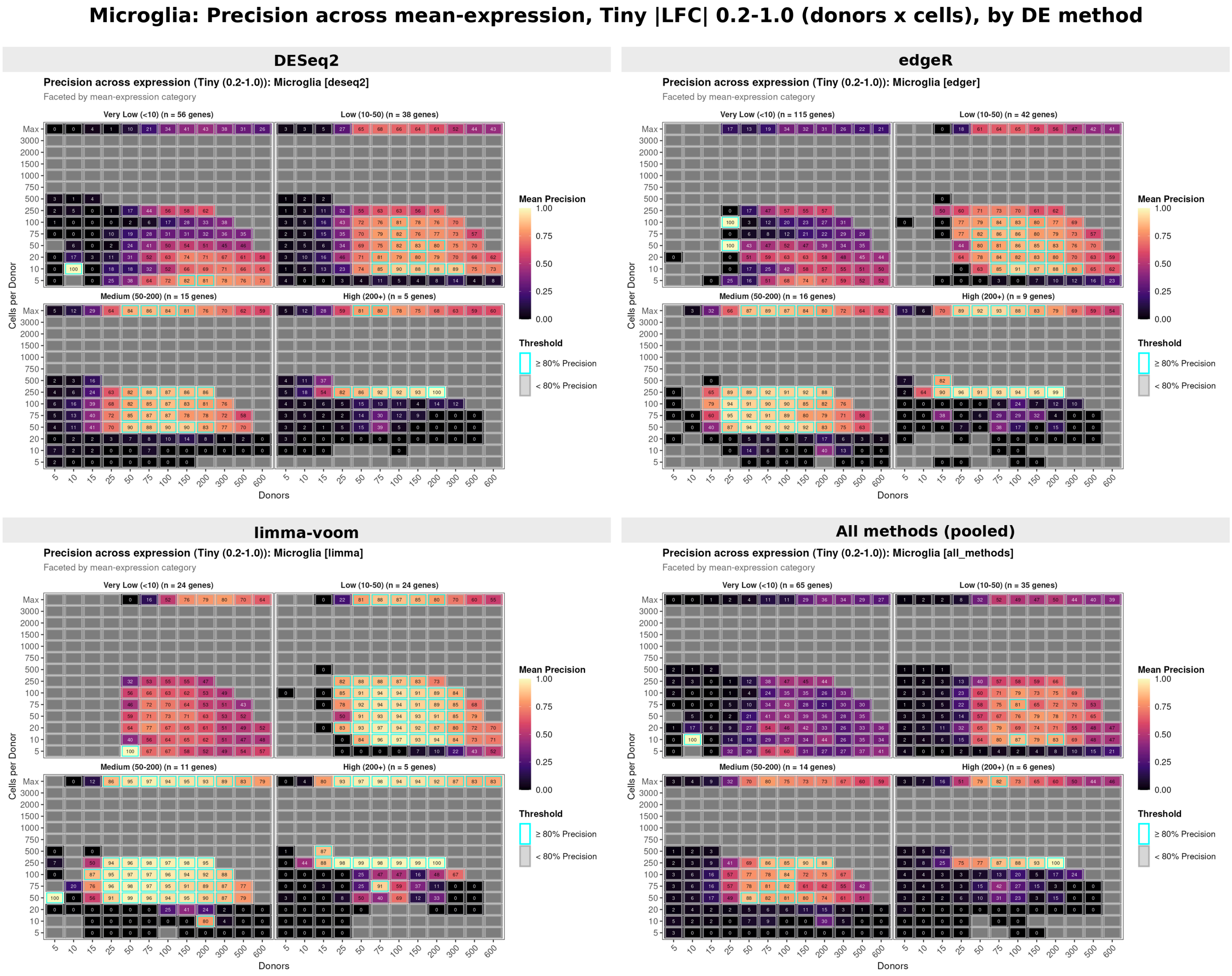


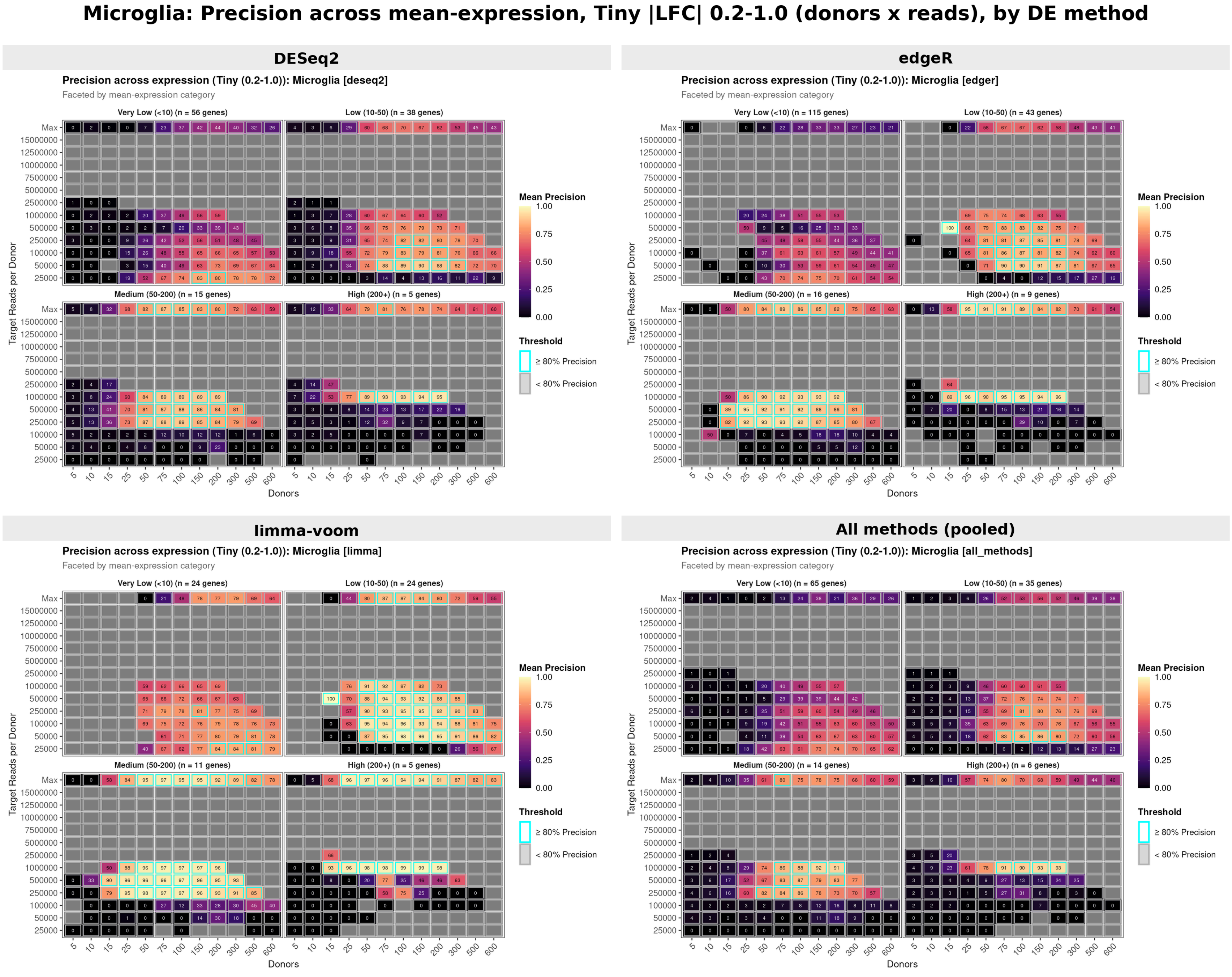

Figure 16: Microglia precision across mean expression within the tiny effect-size bin (|LFC| 0.2-1.0), split by DE method. Cells-based sampling mode (top) and counts-based sampling mode (bottom).

### Supplementary Fig. 16. Microglia: small-effect (|LFC| 1.0-2.0) recall versus mean expression, by DE method

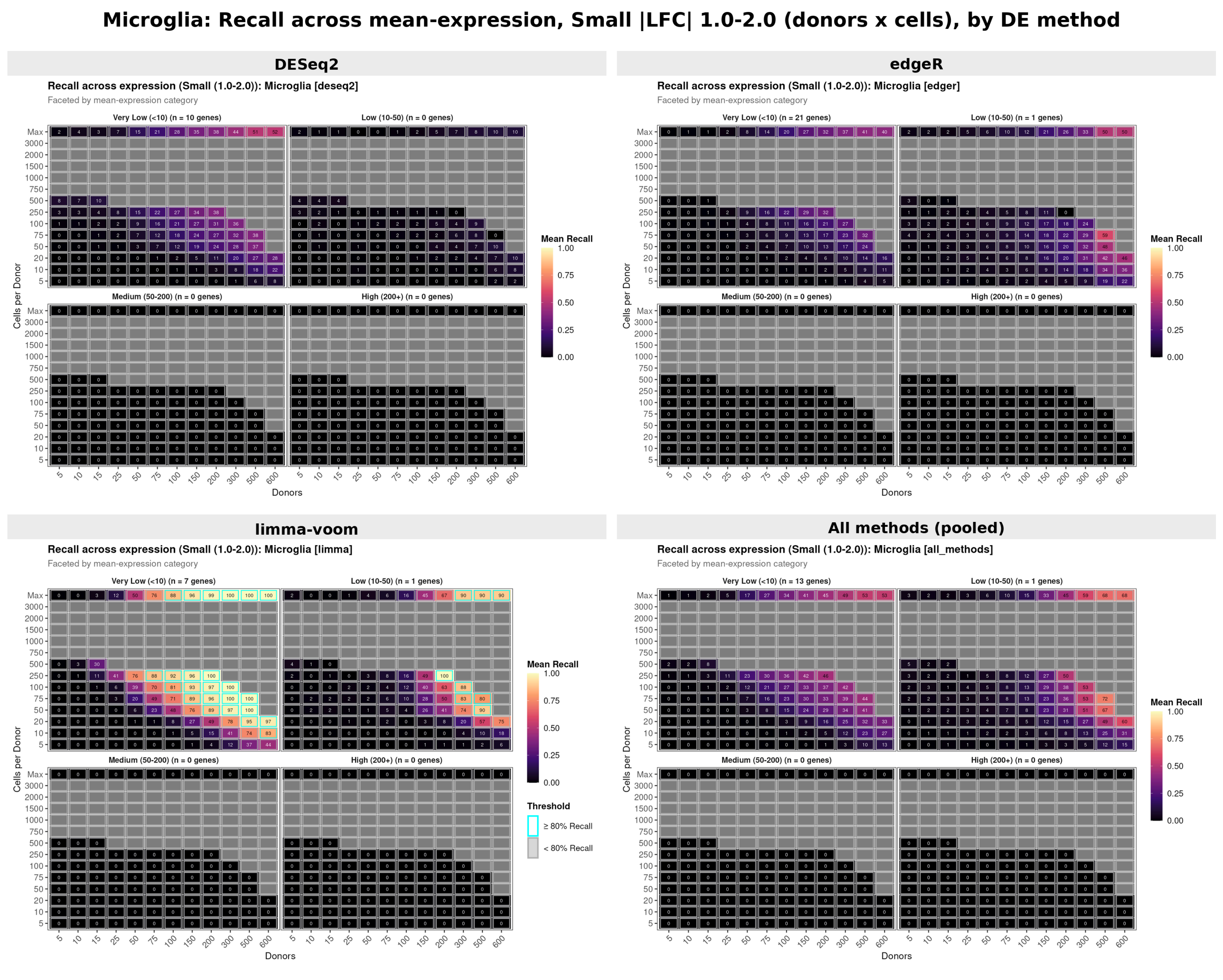


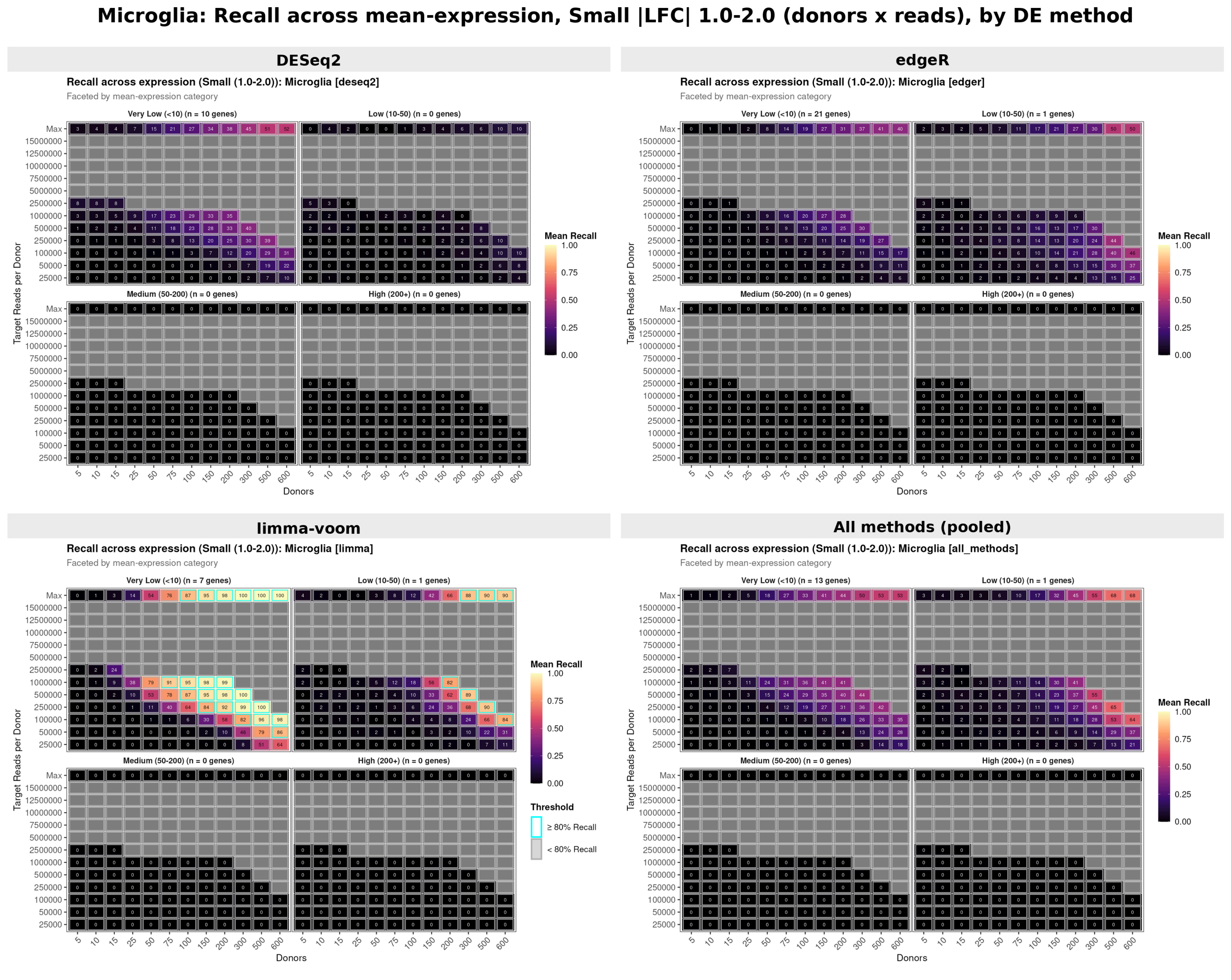


Figure 17: Microglia recall across mean expression within the small effect-size bin (|LFC| 1.0-2.0), split by DE method. Cells-based sampling mode (top) and counts-based sampling mode (bottom).

### Supplementary Fig. 17. Microglia: small-effect (|LFC| 1.0-2.0) precision versus mean expression, by DE method

**Microglia: Precision across mean-expression, Small |LFC| 1.0-2.0 (donors x cells), by DE method**

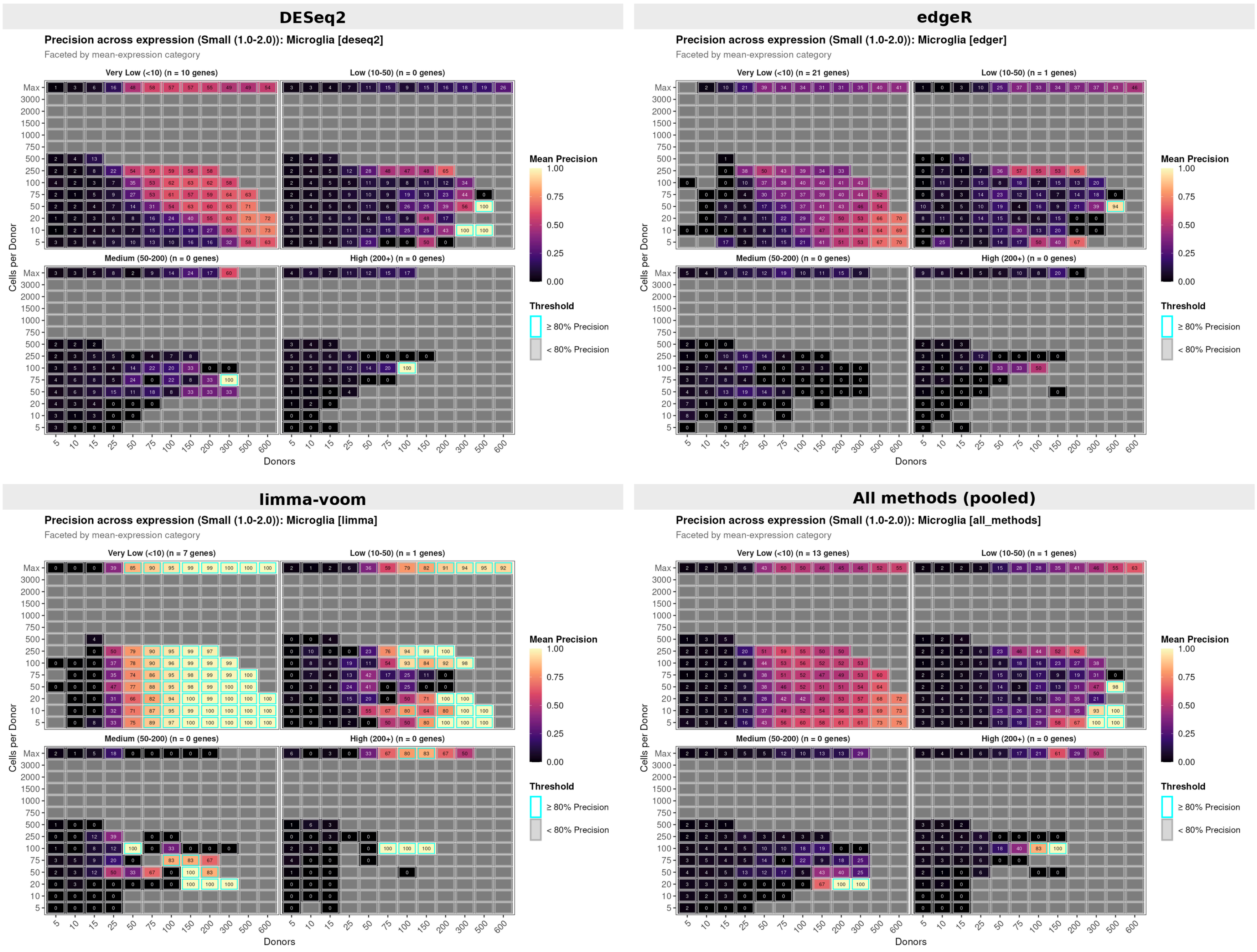


**Microglia: Precision across mean-expression, Small |LFC| 1.0-2.0 (donors x reads), by DE method**

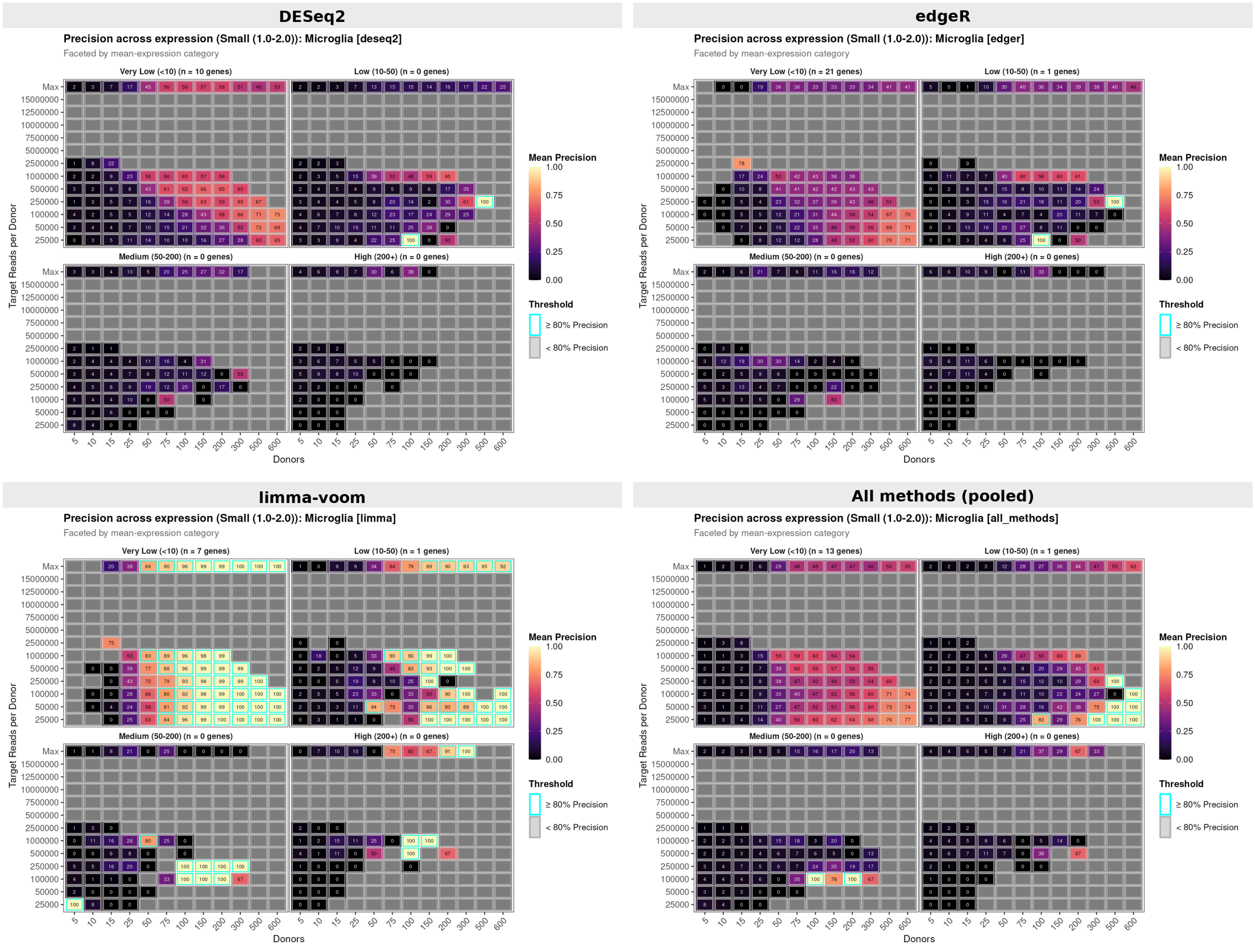

Figure 18: Microglia precision across mean expression within the small effect-size bin (|LFC| 1.0-2.0), split by DE method. Cells-based sampling mode (top) and counts-based sampling mode (bottom).

#### Supplementary Fig. 18. Oligodendrocytes: recall surface across the downsampling grid, by DE method

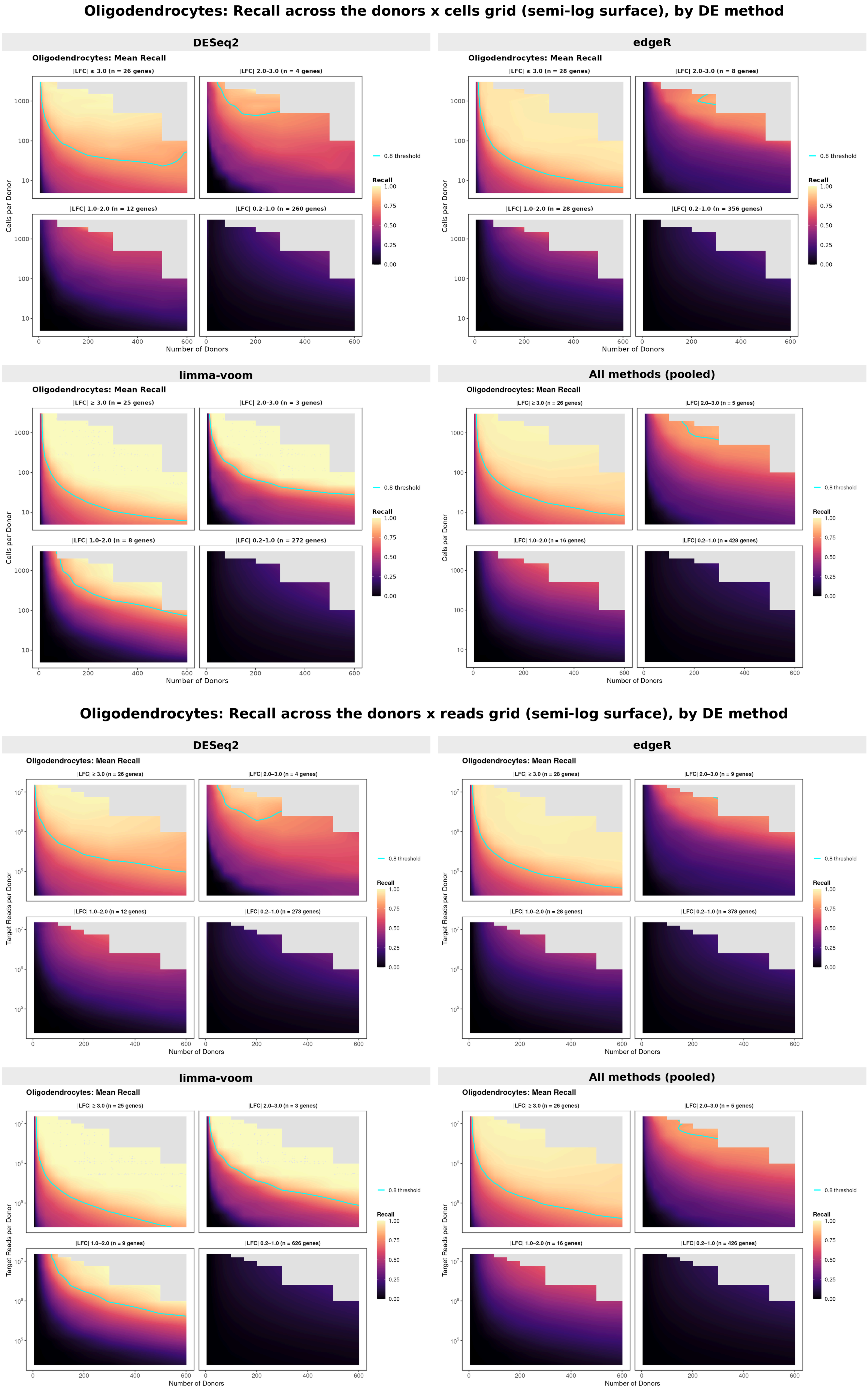


Figure 19: Oligodendrocyte continuous recall surfaces (semi-log scale) split by DE method. Cells-based sampling mode (top) and counts-based sampling mode (bottom).

### Supplementary Fig. 19. Oligodendrocytes: precision surface across the downsampling grid, by DE method

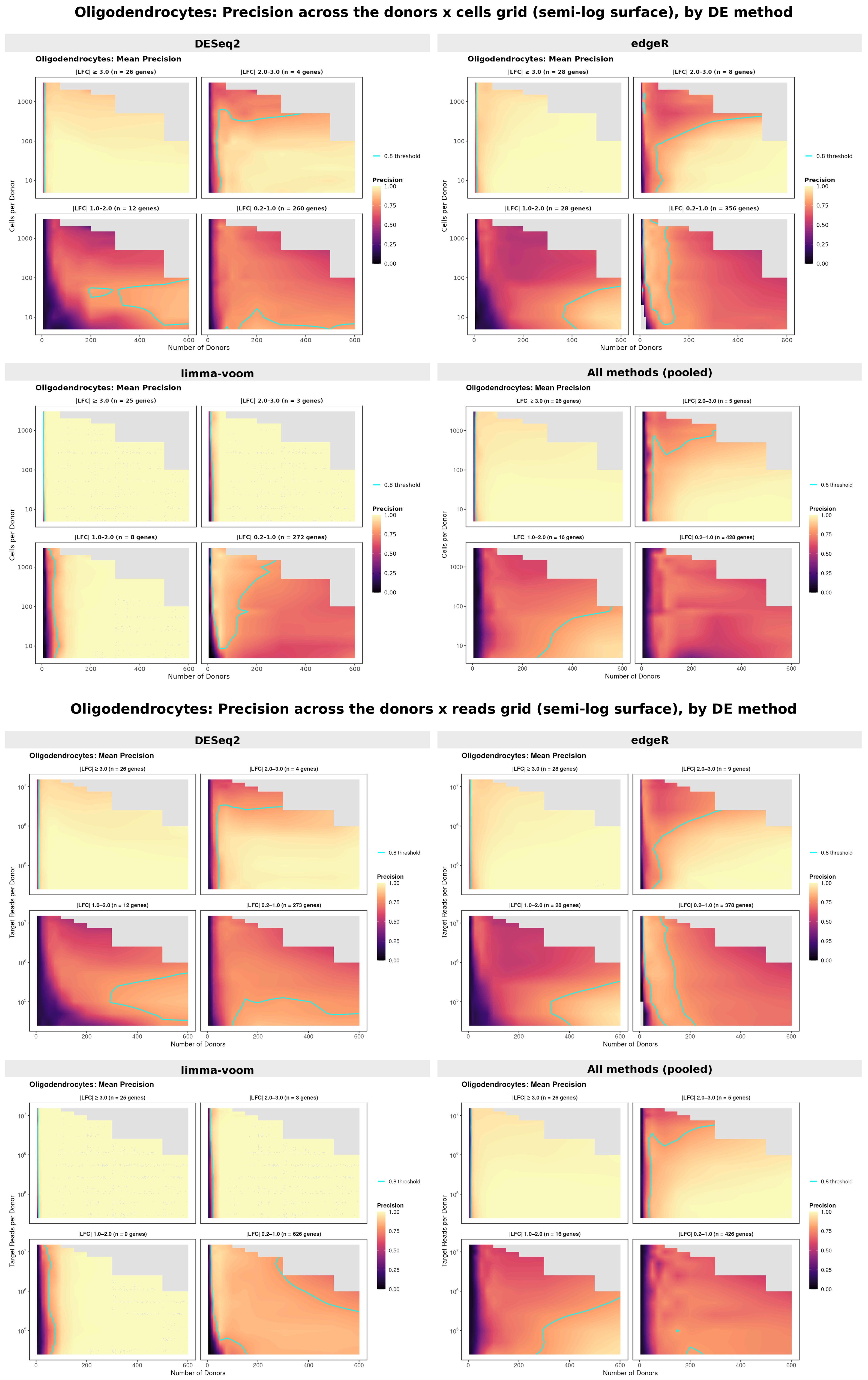

Figure 20: Oligodendrocyte continuous precision surfaces (semi-log scale) split by DE method. Cells-based sampling mode (top) and counts-based sampling mode (bottom).

### Supplementary Fig. 20. Oligodendrocytes: tiny-effect (|LFC| 0.2-1.0) recall versus mean expression, by DE method

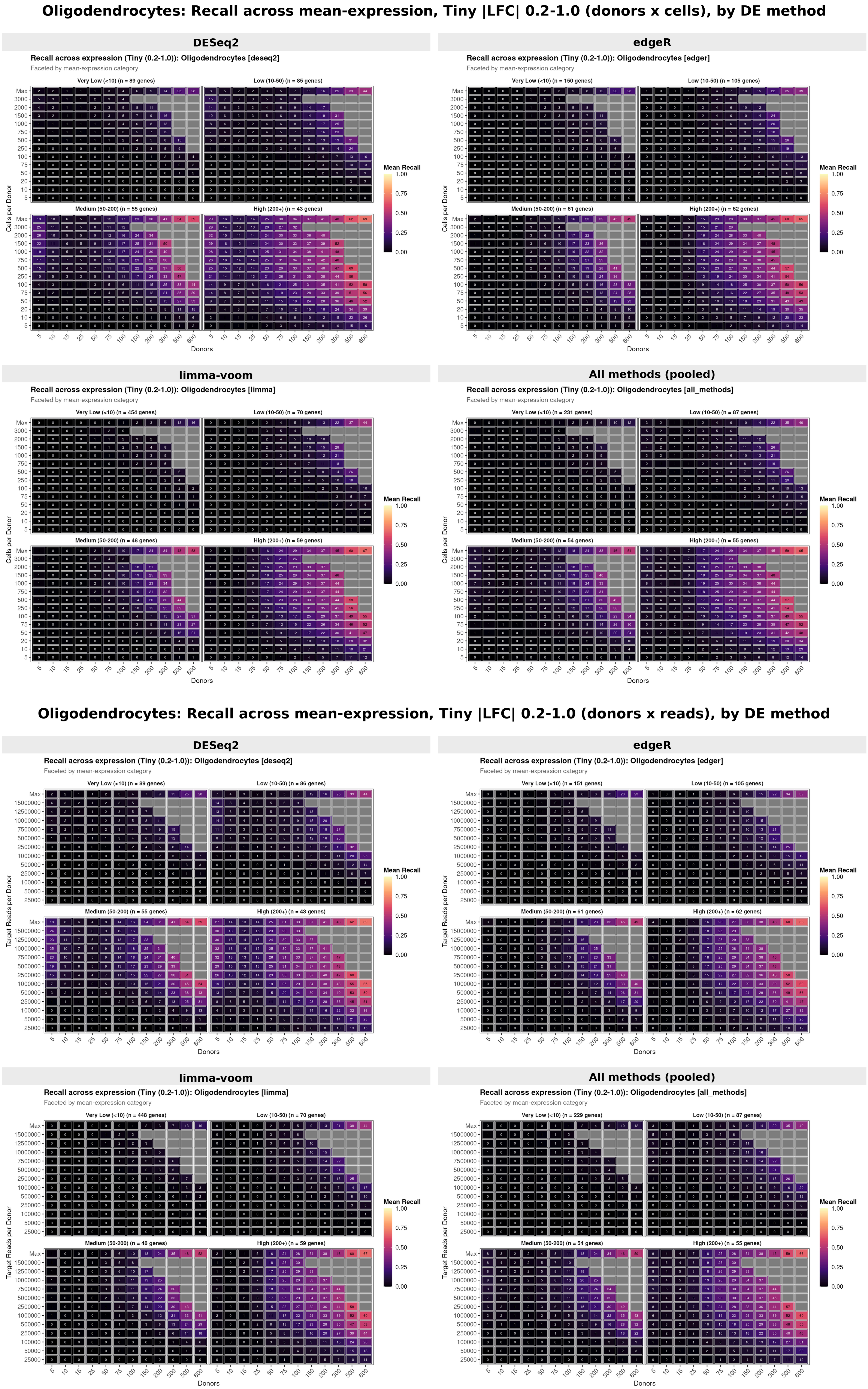


Figure 21: Oligodendrocyte recall across mean expression within the tiny effect-size bin (|LFC| 0.2-1.0), split by DE method. Cells-based sampling mode (top) and counts-based sampling mode (bottom).

### Supplementary Fig. 21. Oligodendrocytes: tiny-effect (|LFC| 0.2-1.0) precision versus mean expression, by DE method

**Oligodendrocytes: Precision across mean-expression, Tiny |LFC| 0.2-1.0 (donors x cells), by DE method**

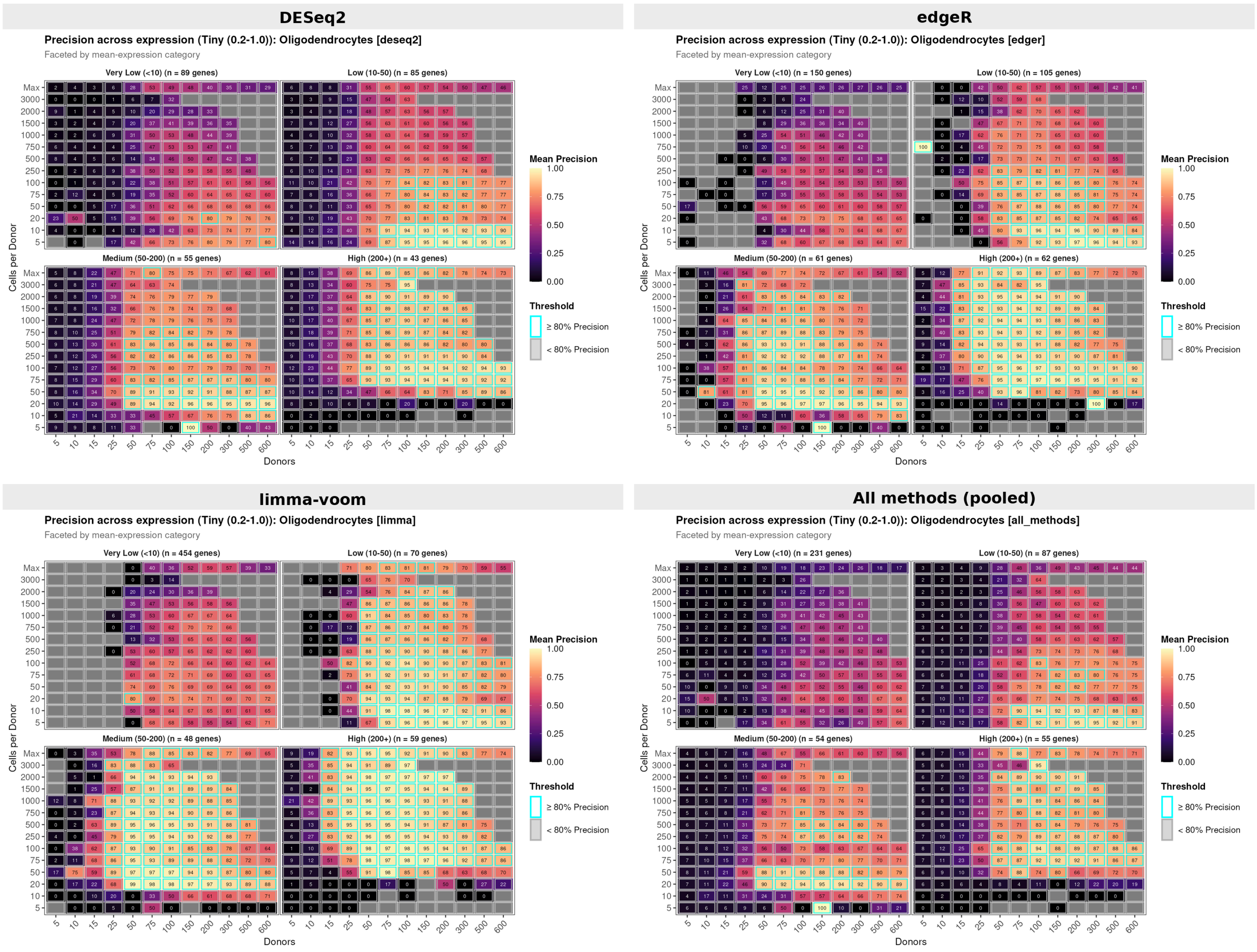


**Oligodendrocytes: Precision across mean-expression, Tiny |LFC| 0.2-1.0 (donors x reads), by DE method**

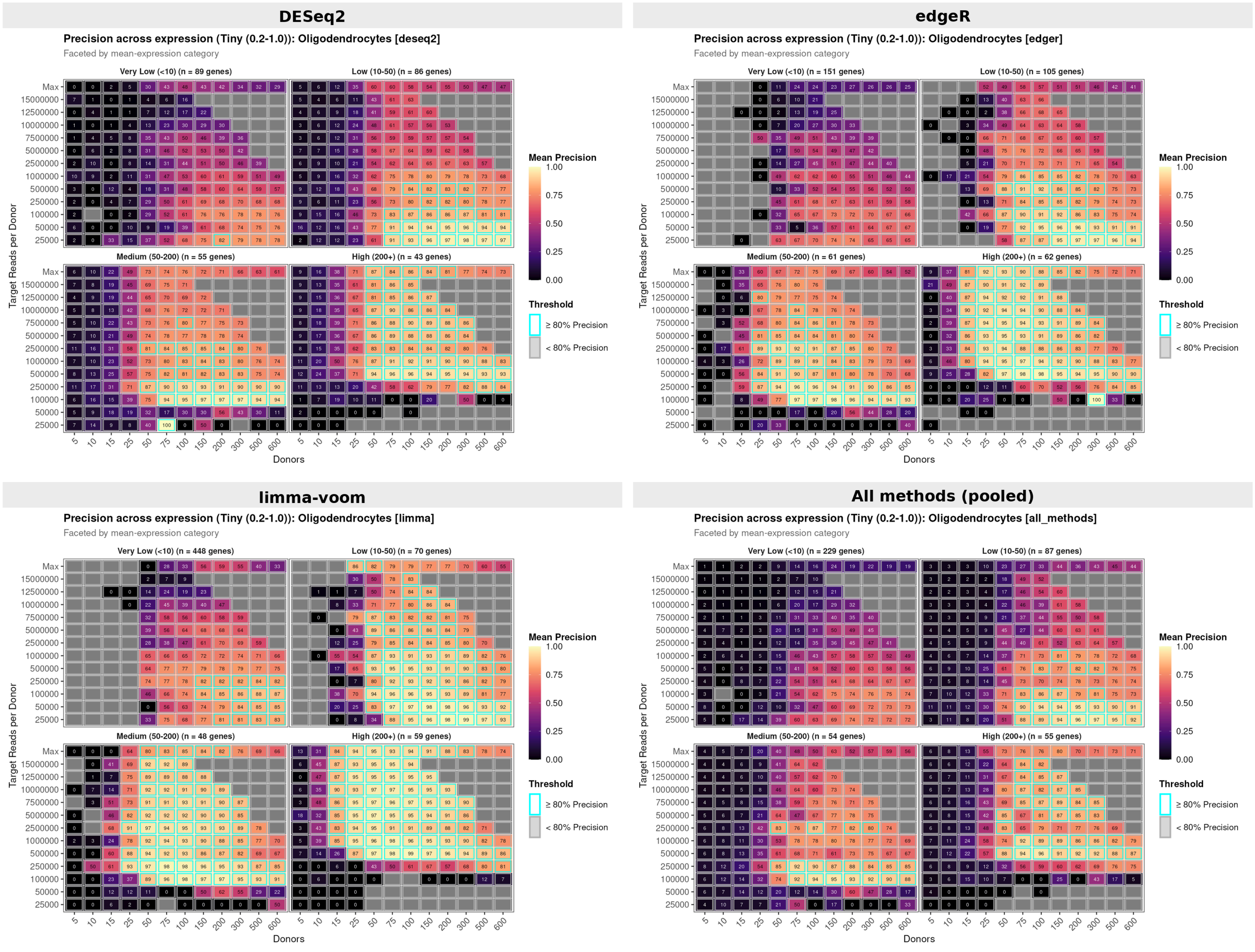

Figure 22: Oligodendrocyte precision across mean expression within the tiny effect-size bin (|LFC| 0.2-1.0), split by DE method. Cells-based sampling mode (top) and counts-based sampling mode (bottom).

### Supplementary Fig. 22. Oligodendrocytes: small-effect (|LFC| 1.0-2.0) recall versus mean expression, by DE method

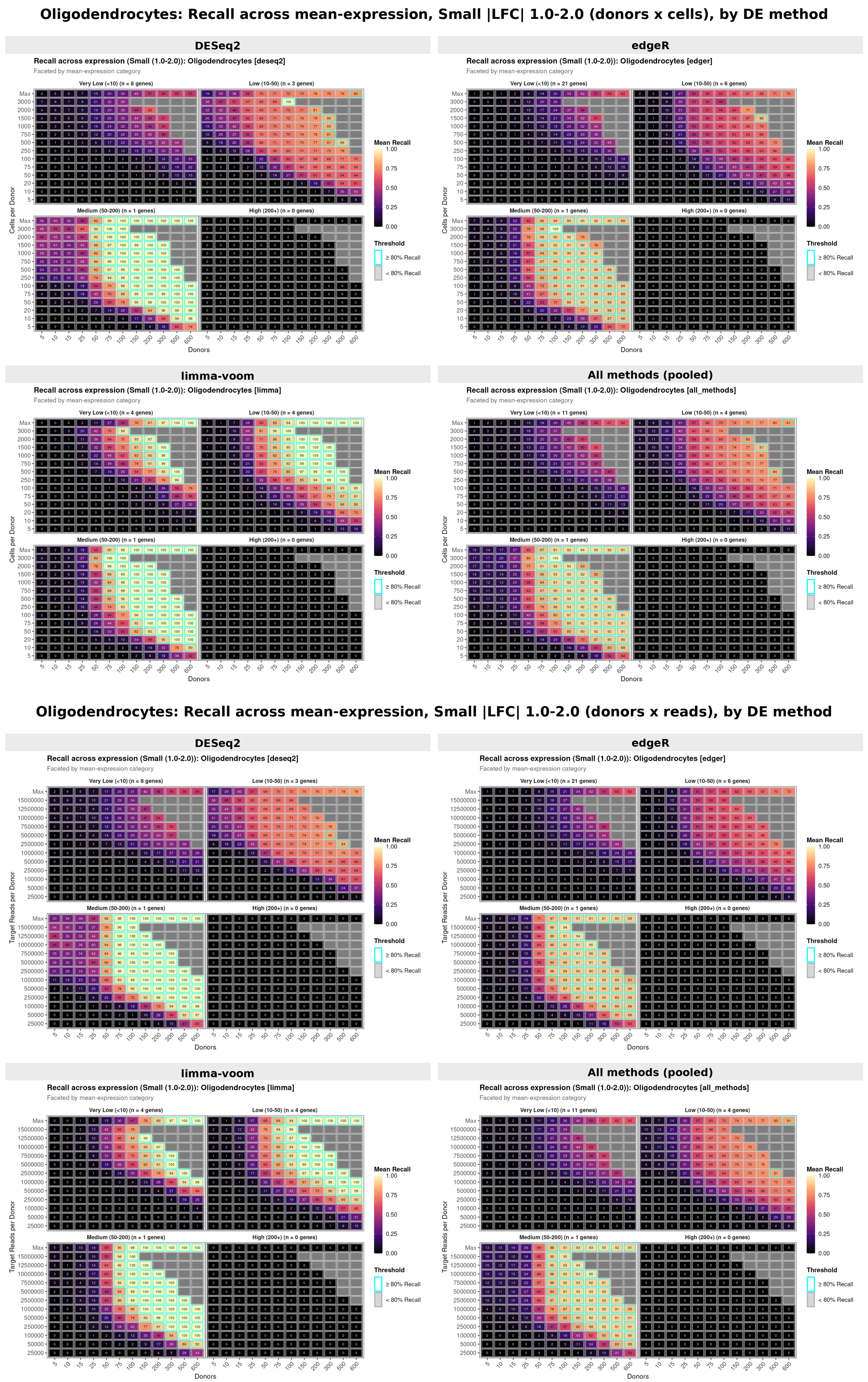


Figure 23: Oligodendrocyte recall across mean expression within the small effect-size bin (|LFC| 1.0-2.0), split by DE method. Cells-based sampling mode (top) and counts-based sampling mode (bottom).

### Supplementary Fig. 23. Oligodendrocytes: small-effect (|LFC| 1.0-2.0) precision versus mean expression, by DE method

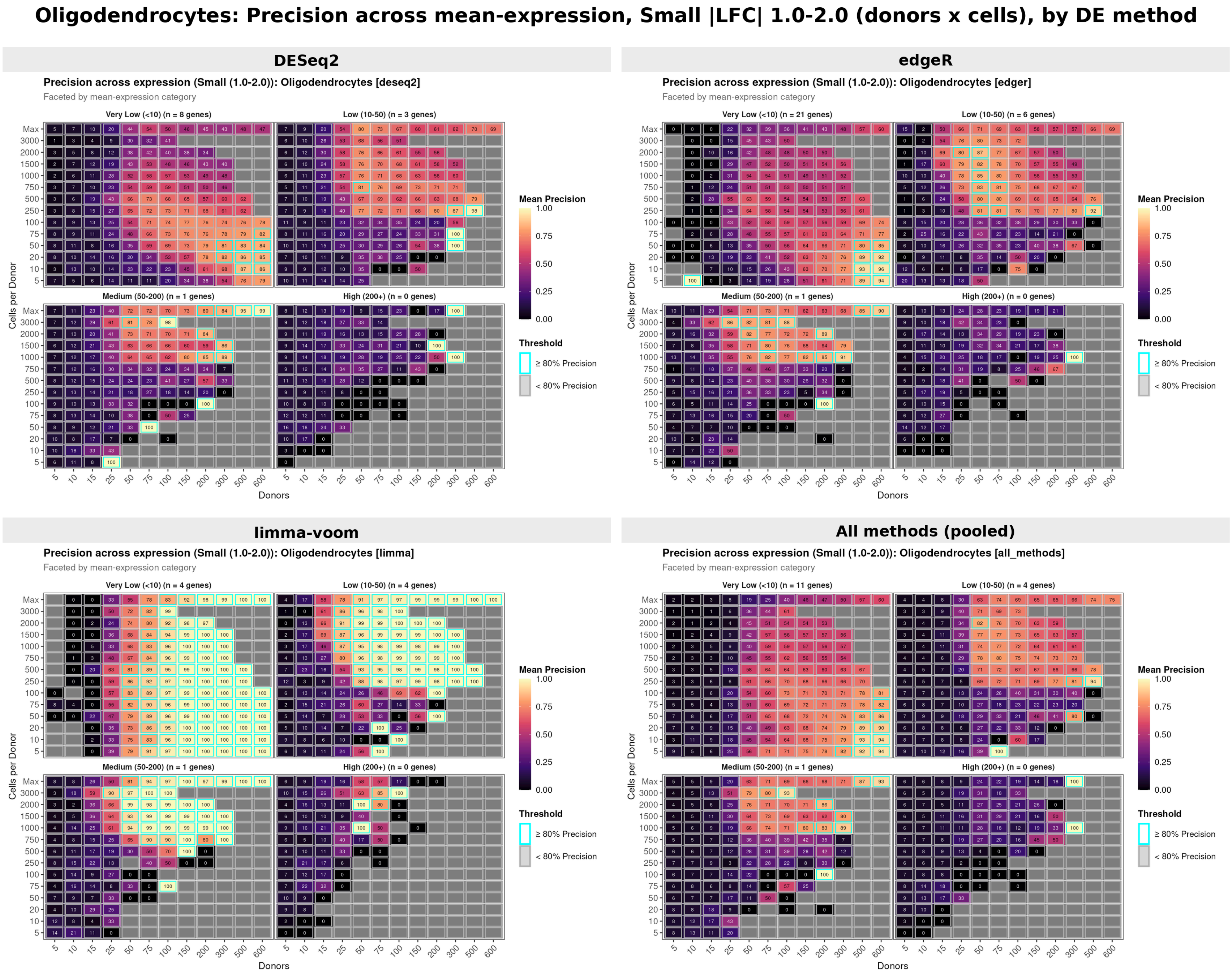


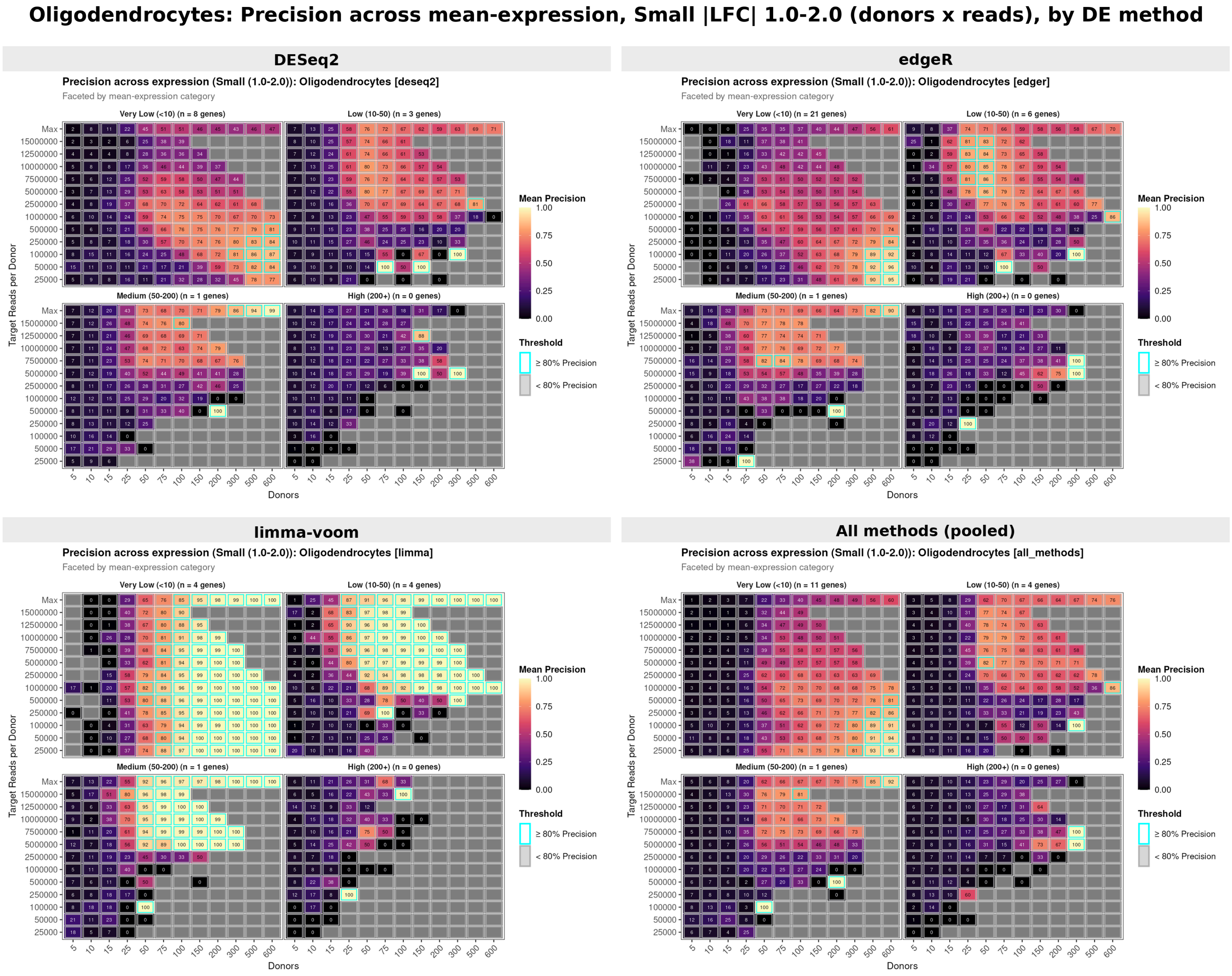

Figure 24: Oligodendrocyte precision across mean expression within the small effect-size bin (|LFC| 1.0-2.0), split by DE method. Cells-based sampling mode (top) and counts-based sampling mode (bottom).

### Supplementary Figures: method-pooled downsampling surfaces

#### Supplementary Fig. 24. Continuous recall surfaces (cells-based and counts-based modes)

Continuous recall surfaces for astrocytes, microglia, and oligodendrocytes under both cells-based sampling, and counts-based sampling. The same qualitative patterns are observed across all three cell types: large-effect genes are readily detected at modest sample sizes, while small-effect genes remain below 80% recall across the entire experimental space. The apparent power ordering (microglia > astrocytes > oligodendrocytes) is an artefact of ground-truth censorship, as explained in the main text.

#### Astrocytes: Mean Recall

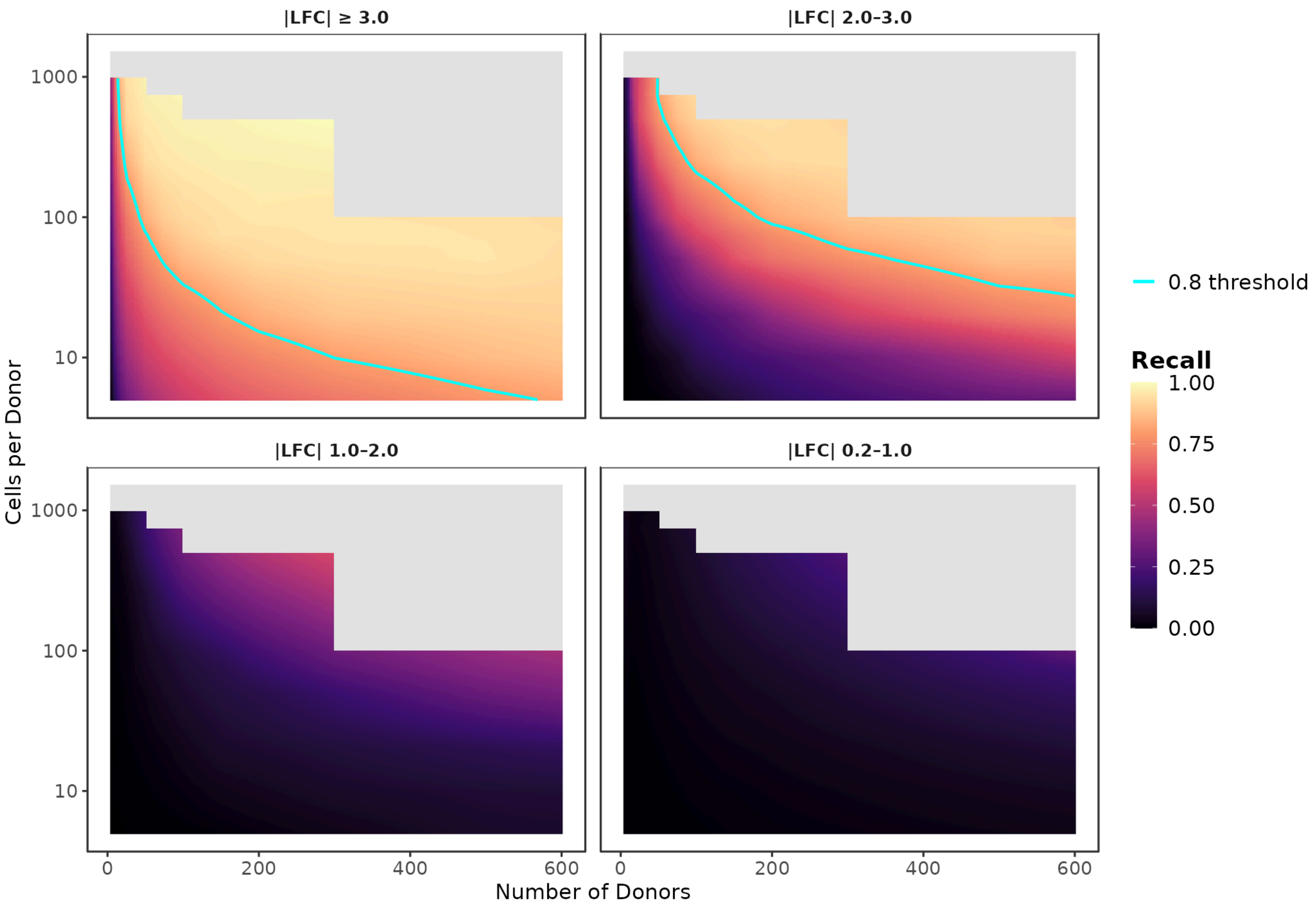


#### Microglia: Mean Recall

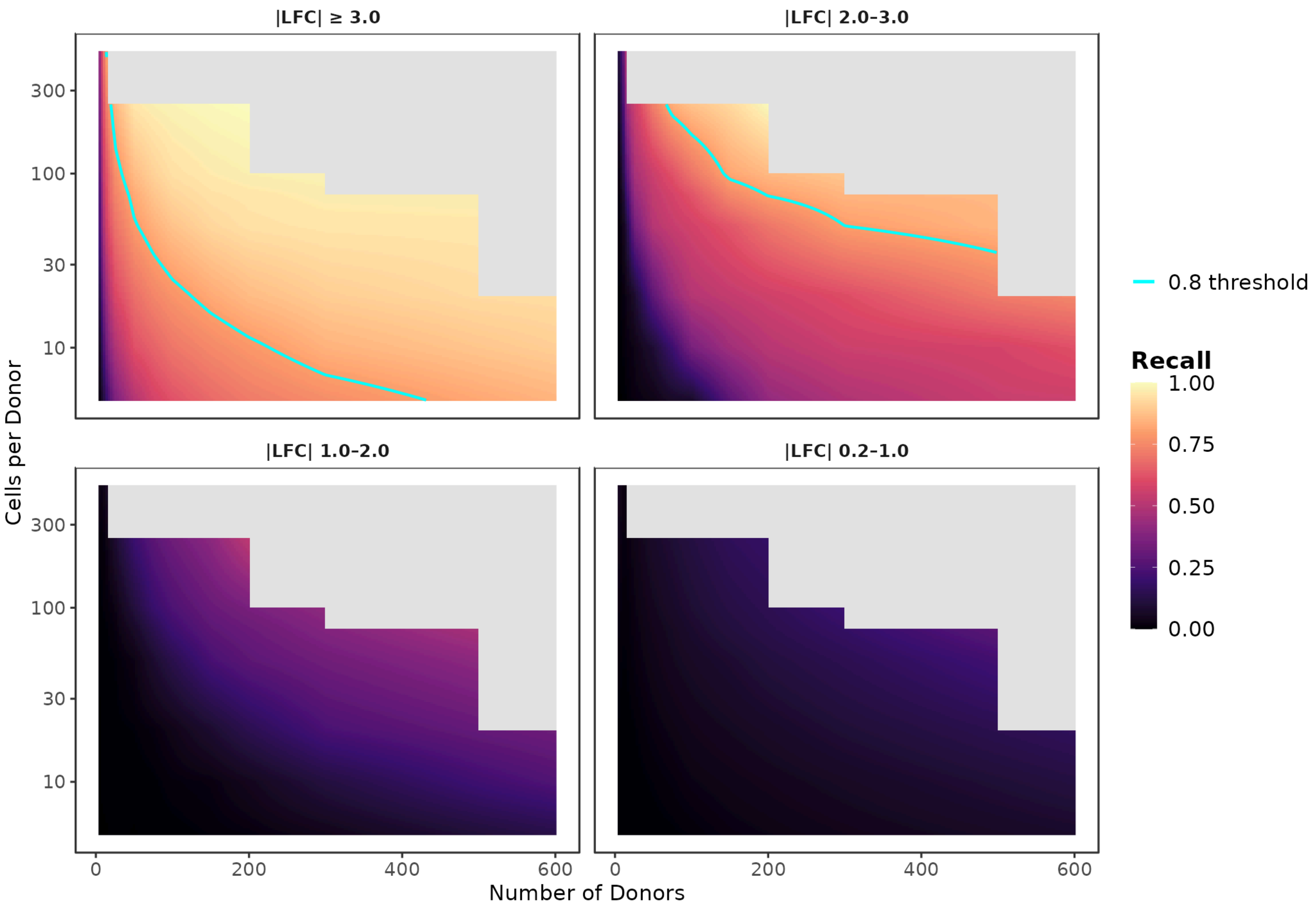

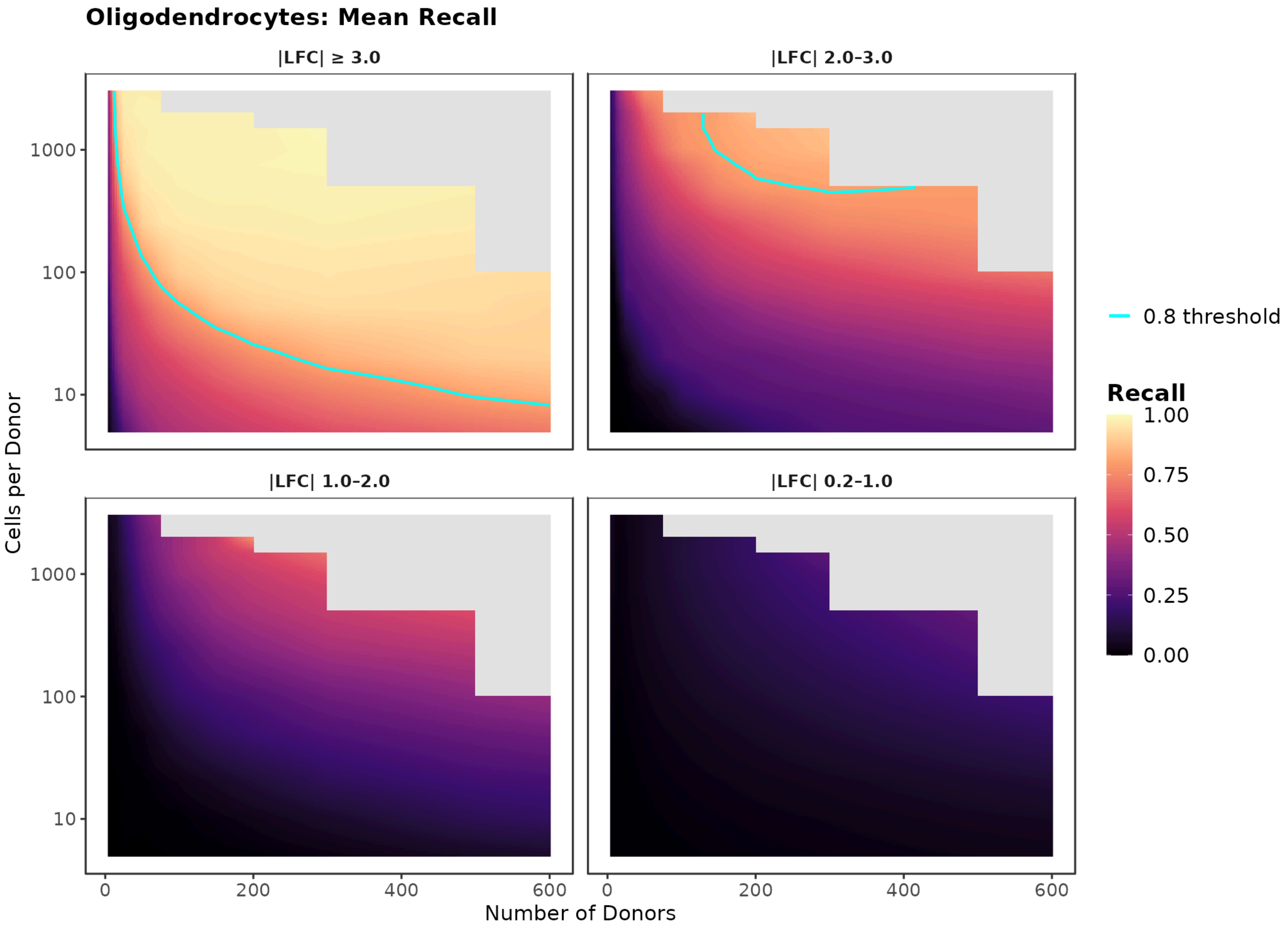


Figure 25: Continuous recall surfaces (semi-log scale, cells-based mode) for astrocytes (top), microglia (middle), and oligodendrocytes (bottom).

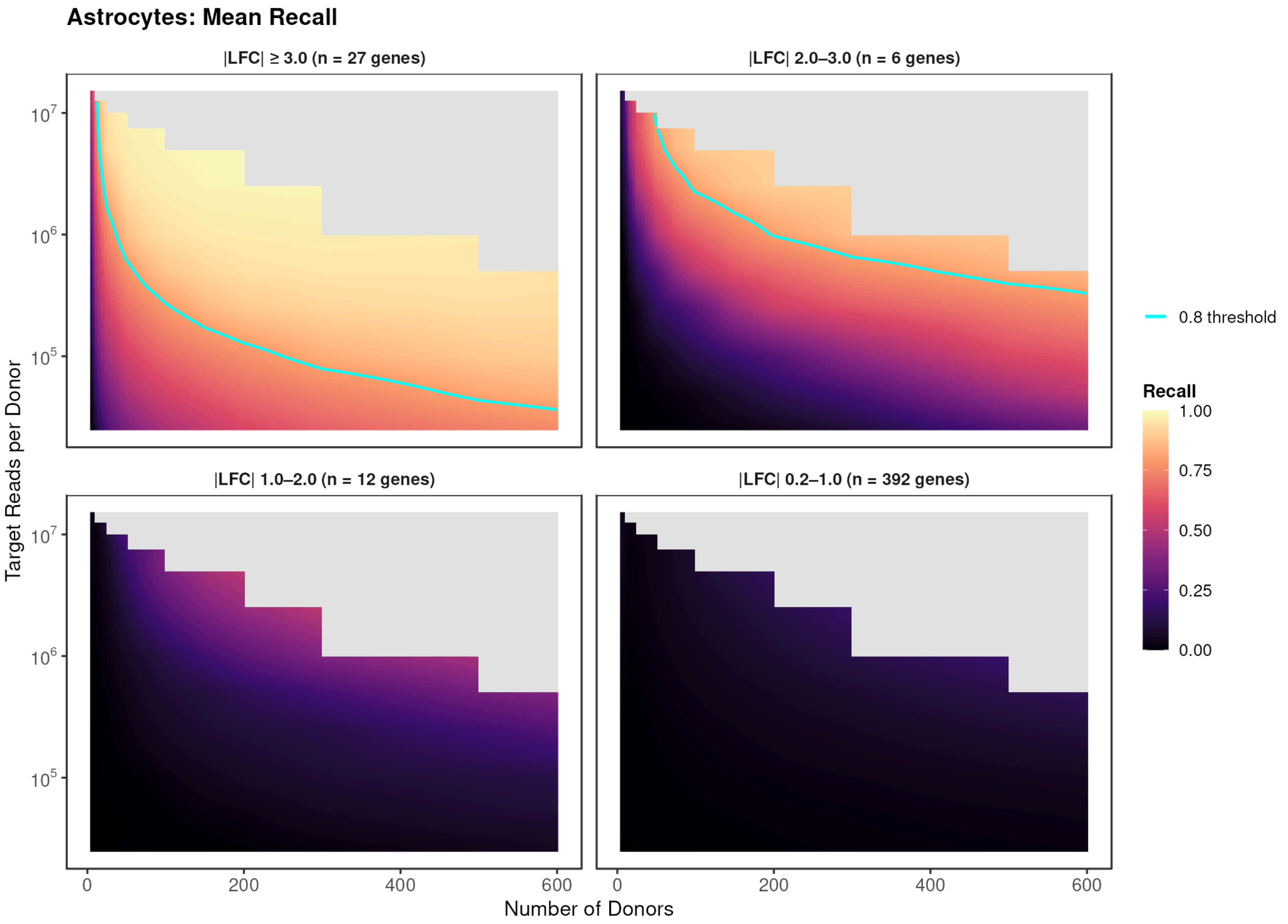
Astrocytes: Mean Recall
|LFC| ≥ 3.0 (n = 27 genes)
|LFC| 2.0–3.0 (n = 6 genes)
|LFC| 1.0–2.0 (n = 12 genes)
|LFC| 0.2–1.0 (n = 392 genes)
Target Reads per Donor
Number of Donors
0.8 threshold
Recall
1.00
0.75
0.50
0.25
0.00

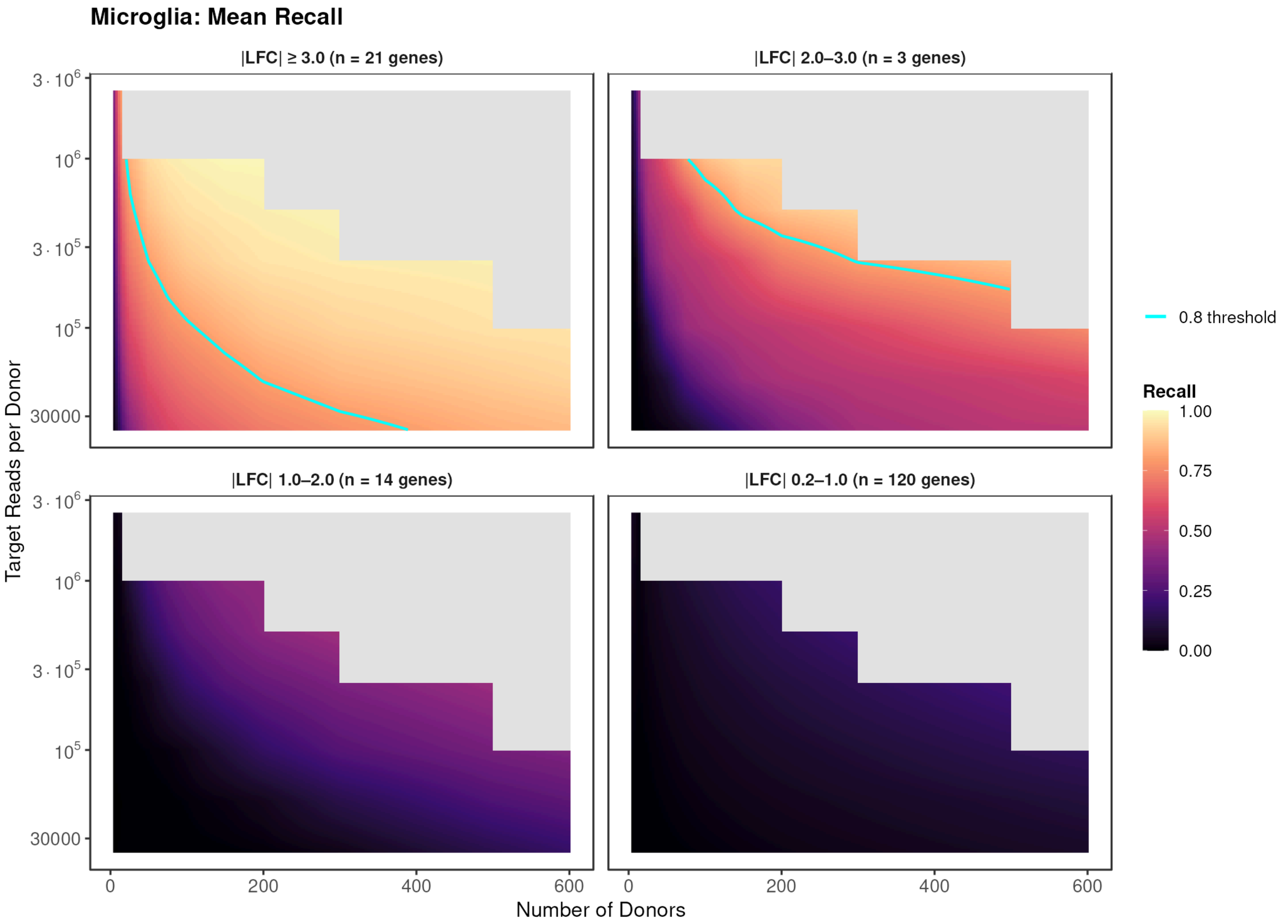
Microglia: Mean Recall
|LFC| ≥ 3.0 (n = 21 genes)
|LFC| 2.0–3.0 (n = 3 genes)
|LFC| 1.0–2.0 (n = 14 genes)
|LFC| 0.2–1.0 (n = 120 genes)
Target Reads per Donor
Number of Donors
0.8 threshold
Recall
1.00
0.75
0.50
0.25
0.00

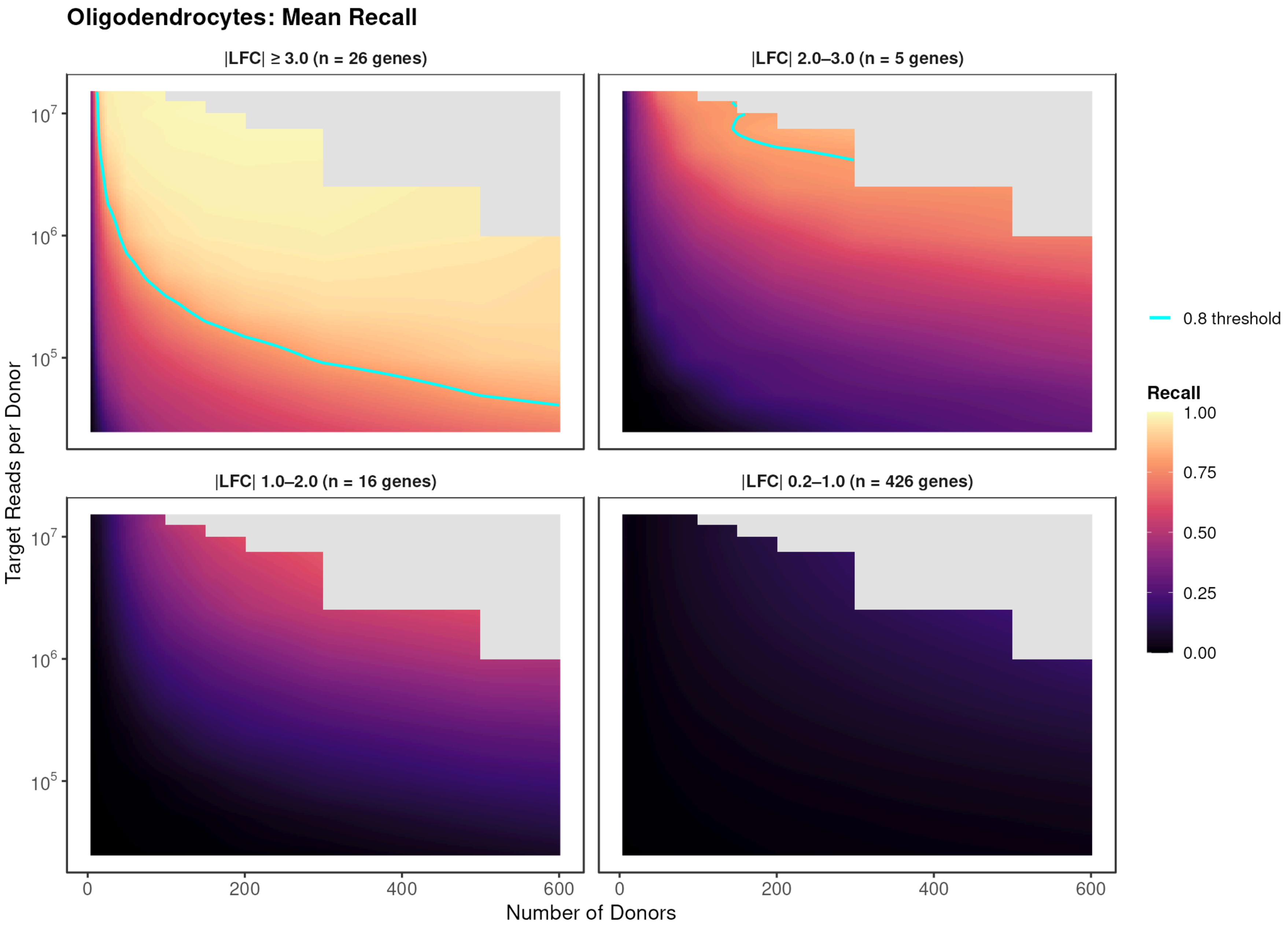


Figure 26: Continuous recall surfaces (semi-log scale, counts-based mode) for astrocytes (top), microglia (middle), and oligodendrocytes (bottom). Same layout as cells-based version above, with counts per cell on the x-axis.

### Supplementary Fig. 25. Precision heatmaps (cells-based and counts-based modes)

Astrocyte, microglial, and oligodendrocyte precision heatmaps.

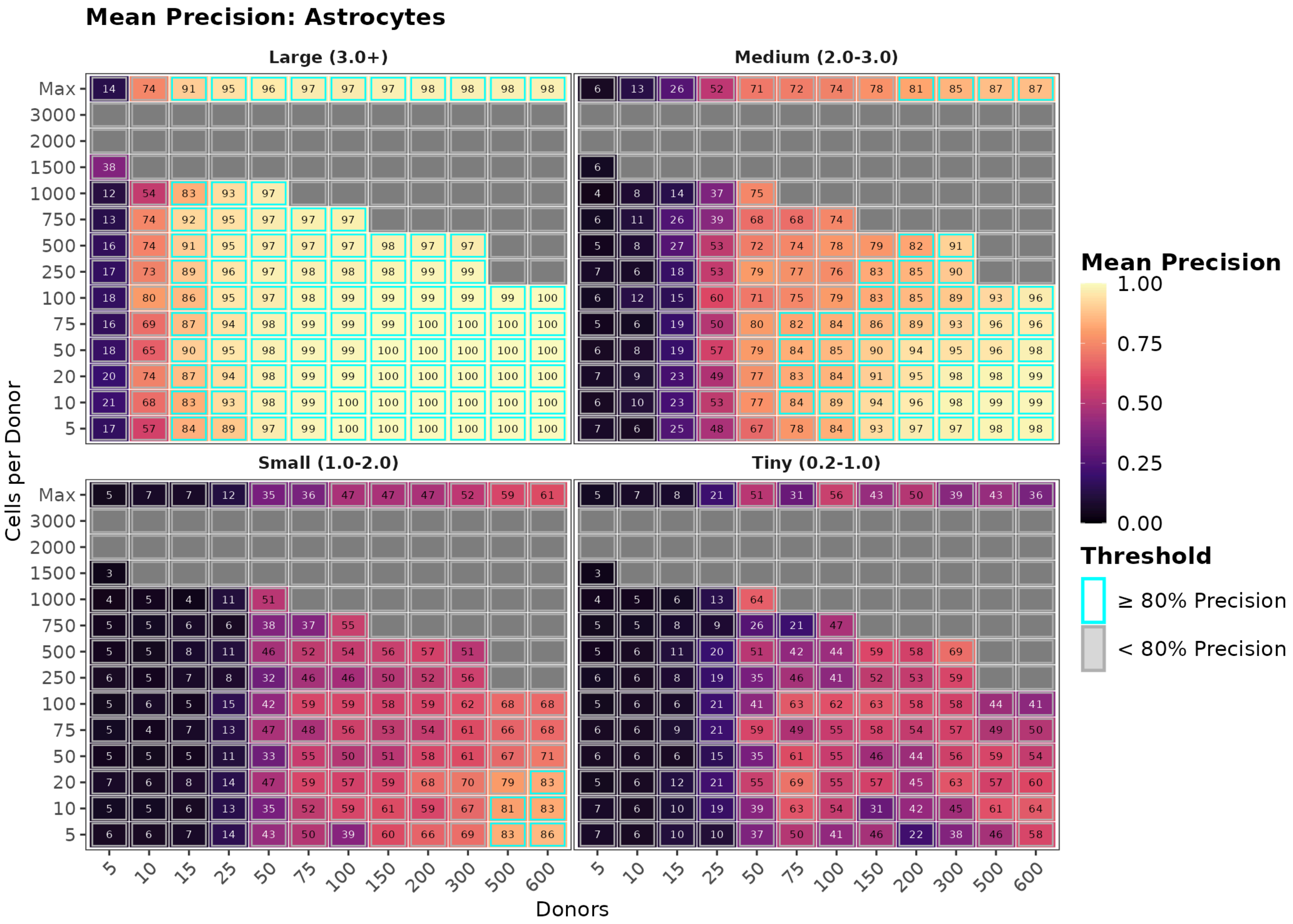
Mean Precision: Astrocytes
Large (3.0+)
Medium (2.0-3.0)
Small (1.0-2.0)
Tiny (0.2-1.0)
Cells per Donor
Max
3000
2000
1500
1000
750
500
250
100
75
50
20
10
5
Donors
5
10
15
25
50
75
100
150
200
300
500
600
Mean Precision
1.00
0.75
0.50
0.25
0.00
Threshold
≥ 80% Precision
< 80% Precision

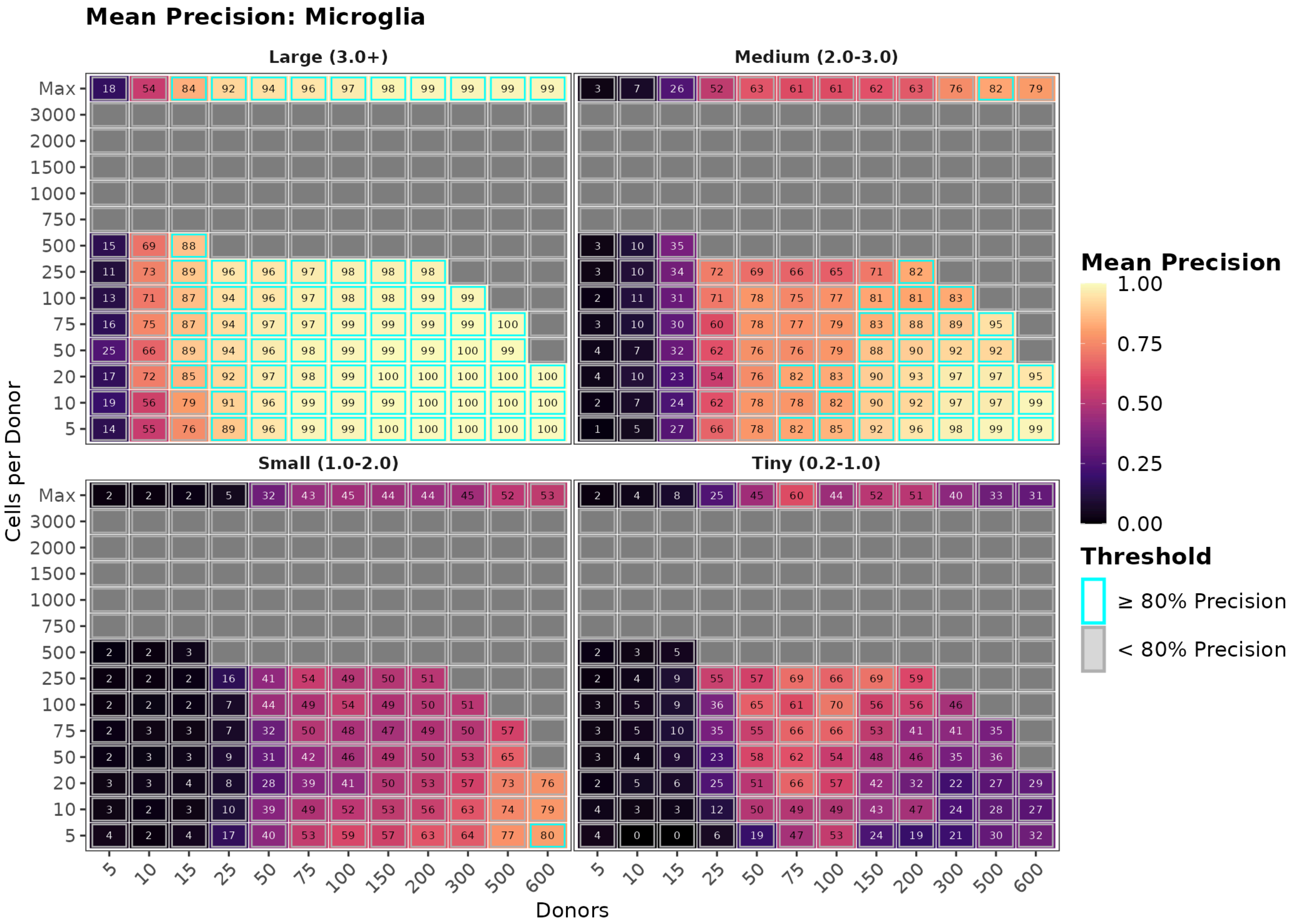
Mean Precision: Microglia
Large (3.0+)
Medium (2.0-3.0)
Small (1.0-2.0)
Tiny (0.2-1.0)
Cells per Donor
Max
3000
2000
1500
1000
750
500
250
100
75
50
20
10
5
Donors
5
10
15
25
50
75
100
150
200
300
500
600
Mean Precision
1.00
0.75
0.50
0.25
0.00
Threshold
≥ 80% Precision
< 80% Precision

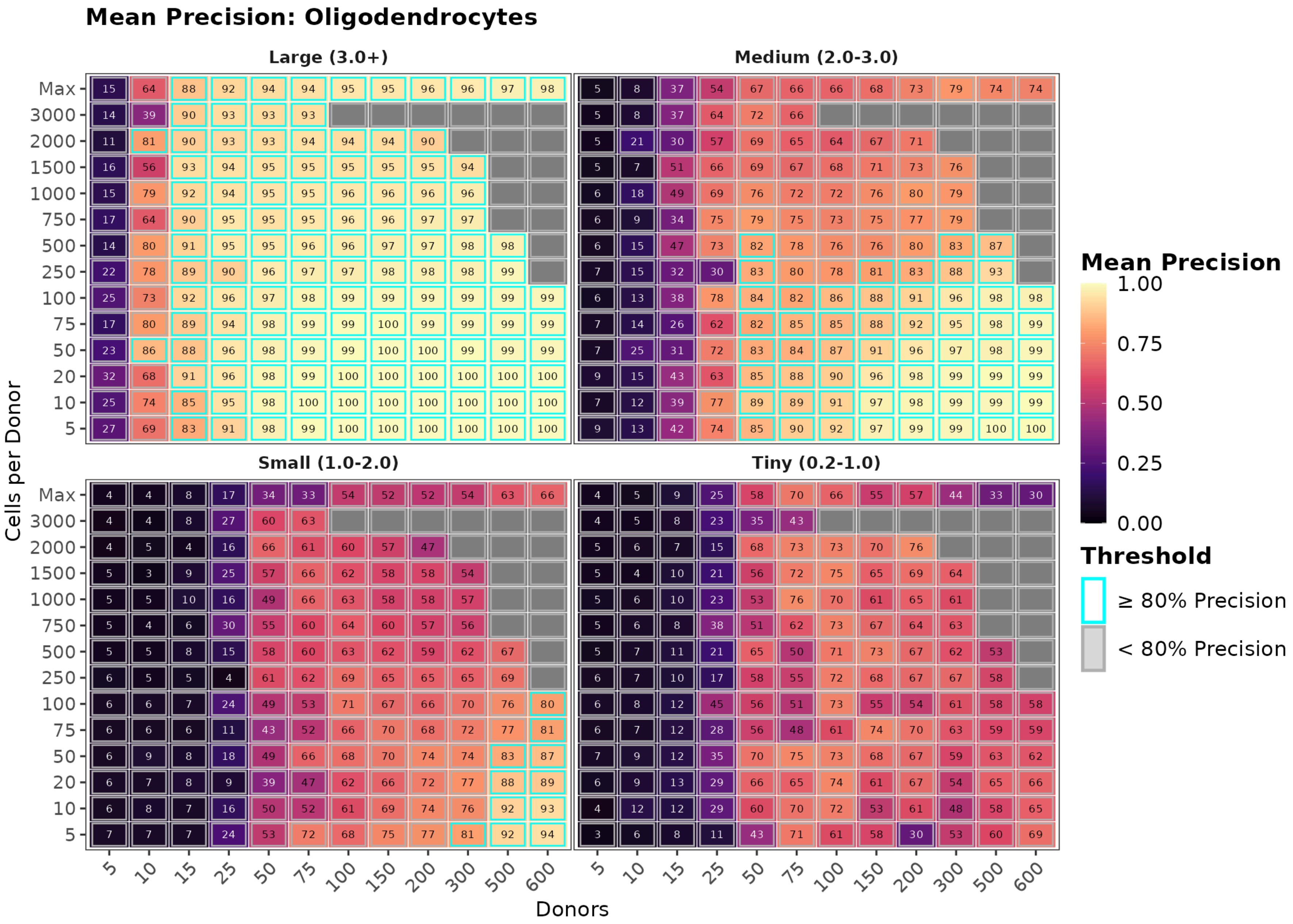


Figure 27: Precision heatmaps for cells-based downsampling, astrocytes (top), microglia (middle), and oligodendrocytes (bottom). Large-effect precision remains >0.90 across most configurations; small-effect precision degrades at higher cell counts, reflecting the precision-recall trade-off.

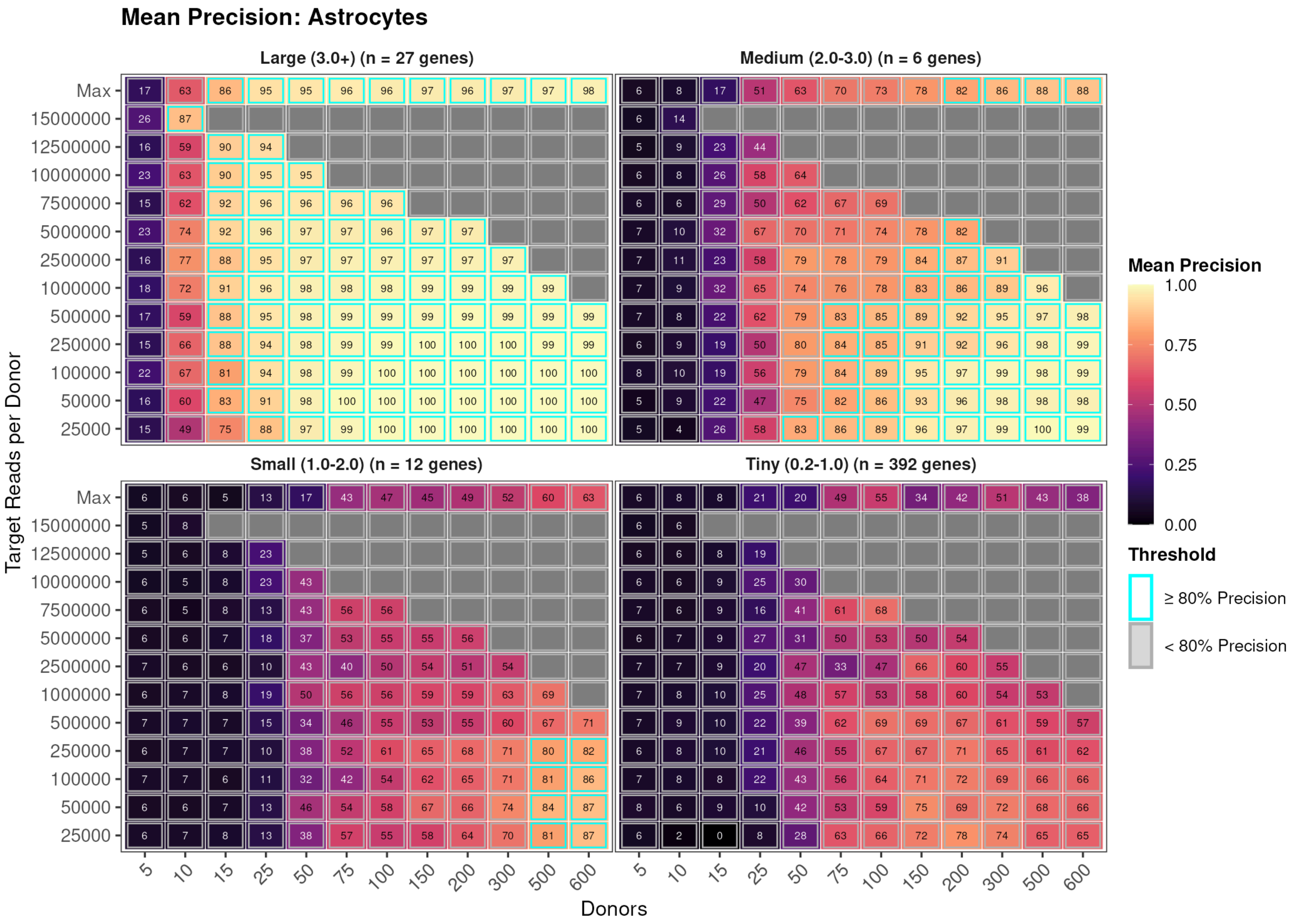
Mean Precision: Astrocytes
Large (3.0+) (n = 27 genes)
Medium (2.0-3.0) (n = 6 genes)
Small (1.0-2.0) (n = 12 genes)
Tiny (0.2-1.0) (n = 392 genes)
Target Reads per Donor
Max
15000000
12500000
10000000
7500000
5000000
2500000
1000000
500000
250000
100000
50000
25000
Donors
5
10
15
25
50
75
100
150
200
300
500
600
Mean Precision
1.00
0.75
0.50
0.25
0.00
Threshold
≥ 80% Precision
< 80% Precision

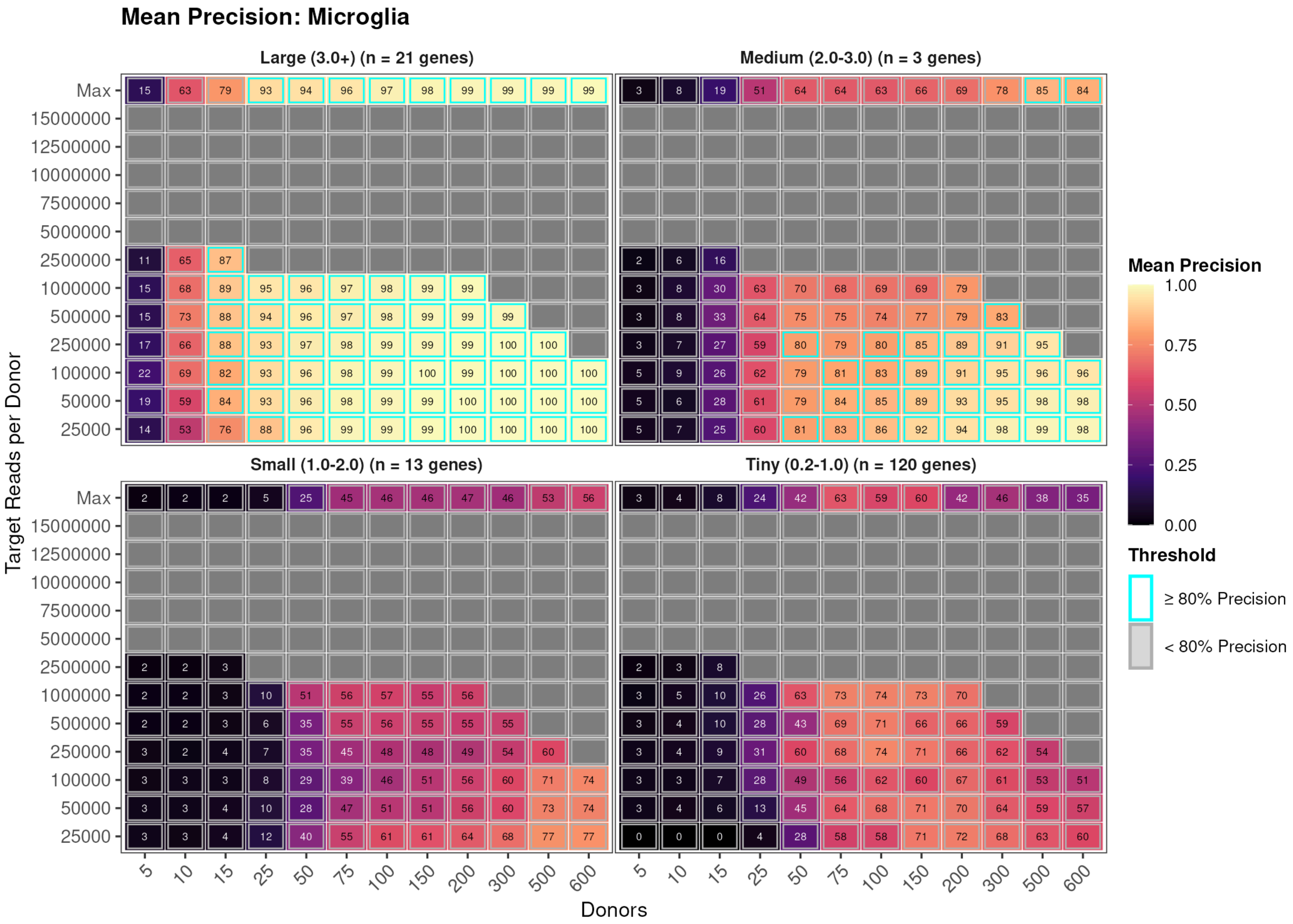
Mean Precision: Microglia
Large (3.0+) (n = 21 genes)
Medium (2.0-3.0) (n = 3 genes)
Small (1.0-2.0) (n = 13 genes)
Tiny (0.2-1.0) (n = 120 genes)
Target Reads per Donor
Max
15000000
12500000
10000000
7500000
5000000
2500000
1000000
500000
250000
100000
50000
25000
Donors
5
10
15
25
50
75
100
150
200
300
500
600
Mean Precision
1.00
0.75
0.50
0.25
0.00
Threshold
≥ 80% Precision
< 80% Precision

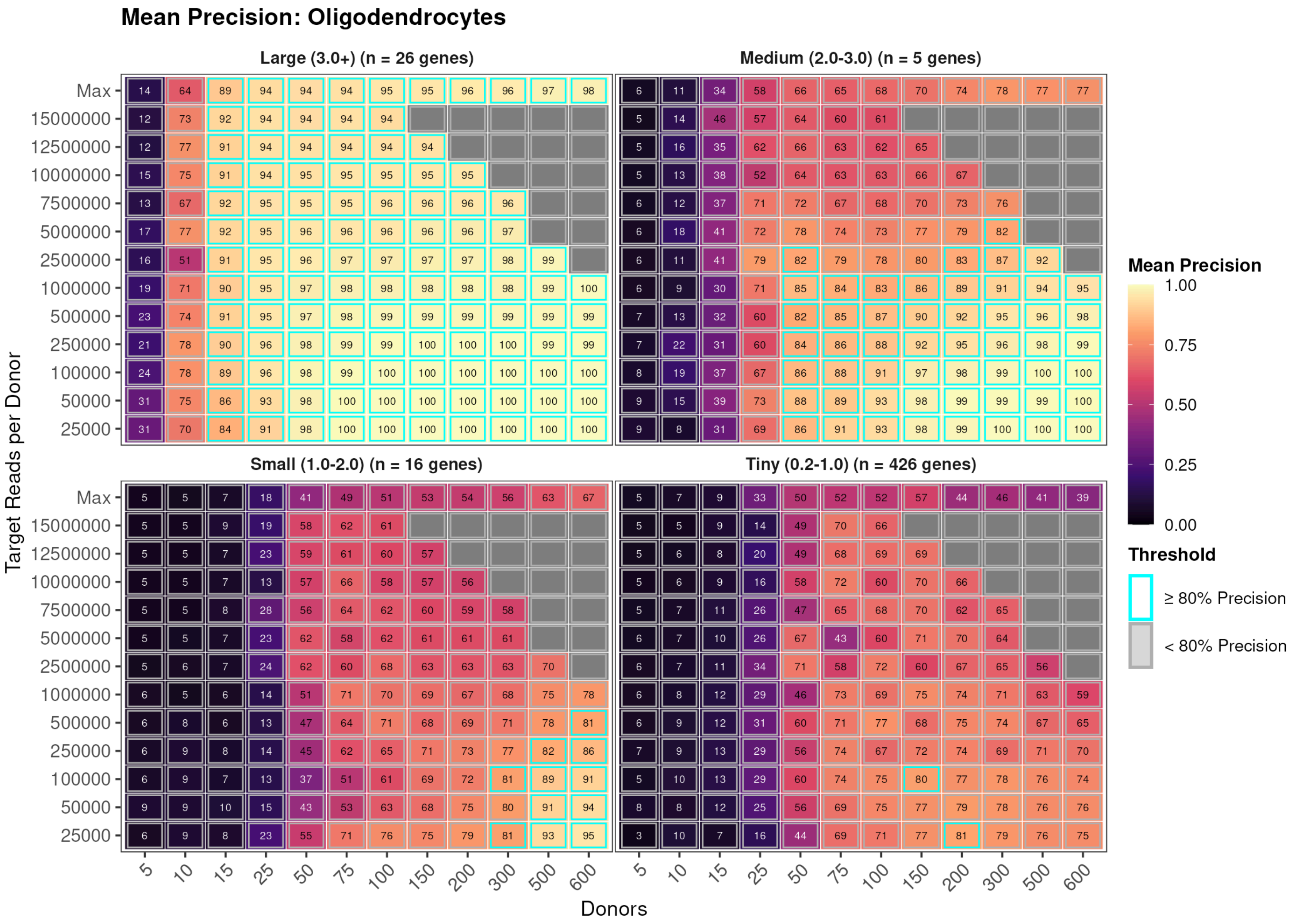


Figure 28: Precision heatmaps for counts-based downsampling, astrocytes (top), microglia (middle), and oligodendrocytes (bottom). Layout identical to cells-based mode, but columns represent counts per cell rather than cells per donor.

### Supplementary Fig. 26. Discrete recall heatmaps (cells-based and counts-based modes)

Discrete recall grids for astrocytes, microglia, and oligodendrocytes under both cells-based sampling, and counts-based sampling. The same qualitative patterns are observed across all three cell types: large-effect genes are readily detected at modest sample sizes, while small-effect genes remain below 80% recall across the entire experimental space. The apparent power ordering (microglia > astrocytes > oligodendrocytes) is an artefact of ground-truth censorship, as explained in the main text.

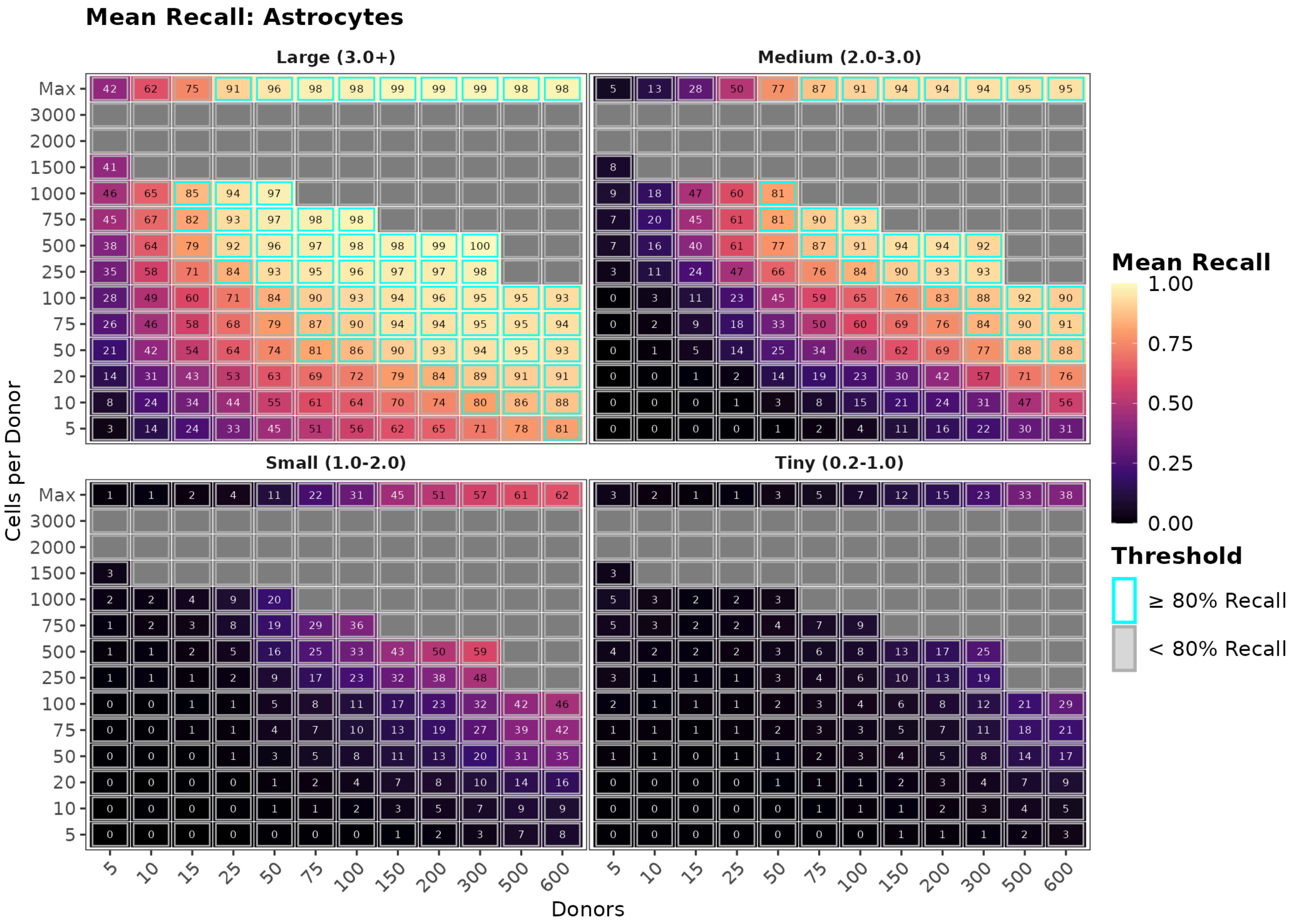
Mean Recall: Astrocytes
Large (3.0+)
Medium (2.0-3.0)
Small (1.0-2.0)
Tiny (0.2-1.0)
Cells per Donor
Max
3000
2000
1500
1000
750
500
250
100
75
50
20
10
5
Donors
5
10
15
25
50
75
100
150
200
300
500
600
Mean Recall
1.00
0.75
0.50
0.25
0.00
Threshold
≥ 80% Recall
< 80% Recall

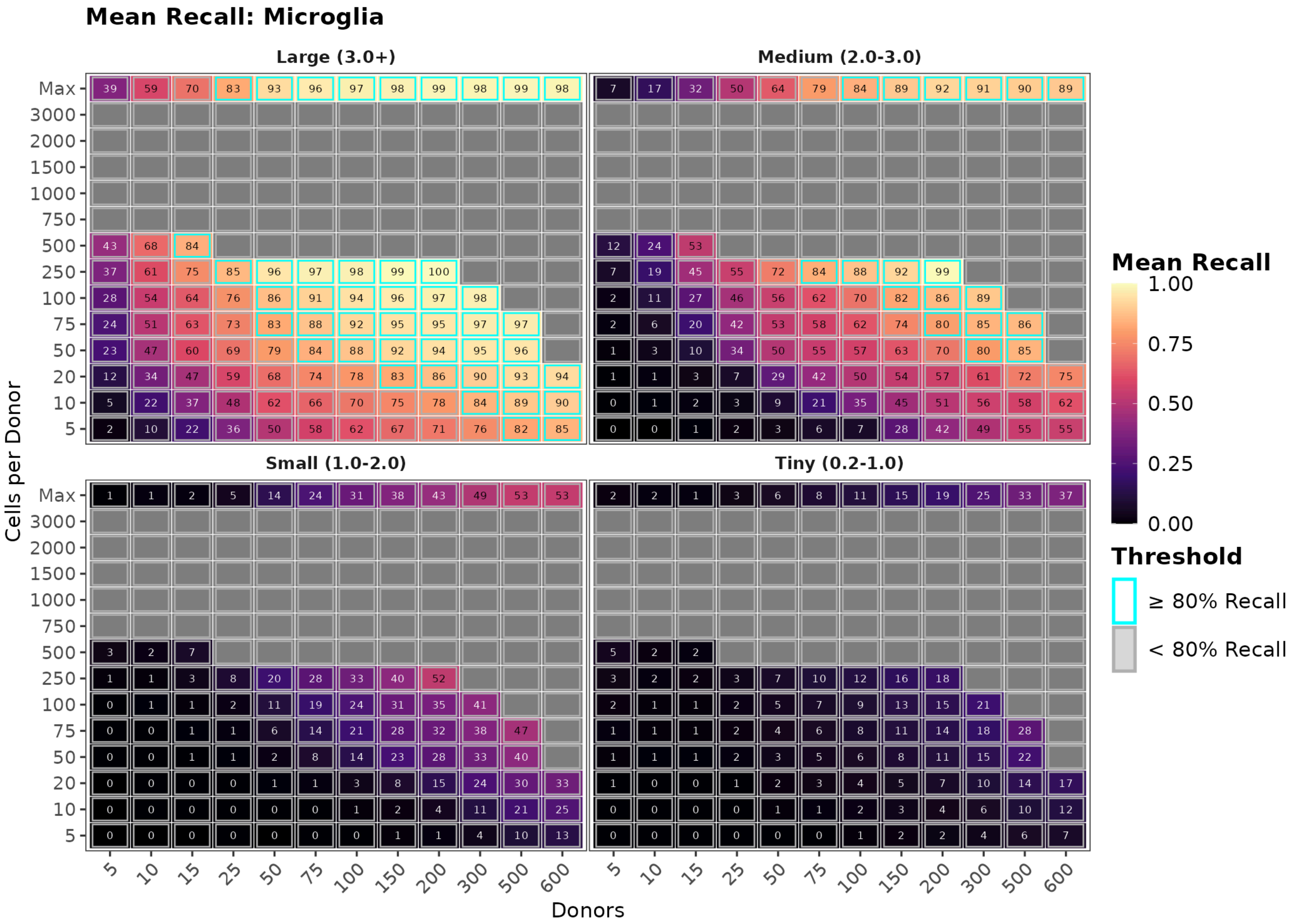
Mean Recall: Microglia
Large (3.0+)
Medium (2.0-3.0)
Small (1.0-2.0)
Tiny (0.2-1.0)
Cells per Donor
Max
3000
2000
1500
1000
750
500
250
100
75
50
20
10
5
Donors
5
10
15
25
50
75
100
150
200
300
500
600
Mean Recall
1.00
0.75
0.50
0.25
0.00
Threshold
≥ 80% Recall
< 80% Recall

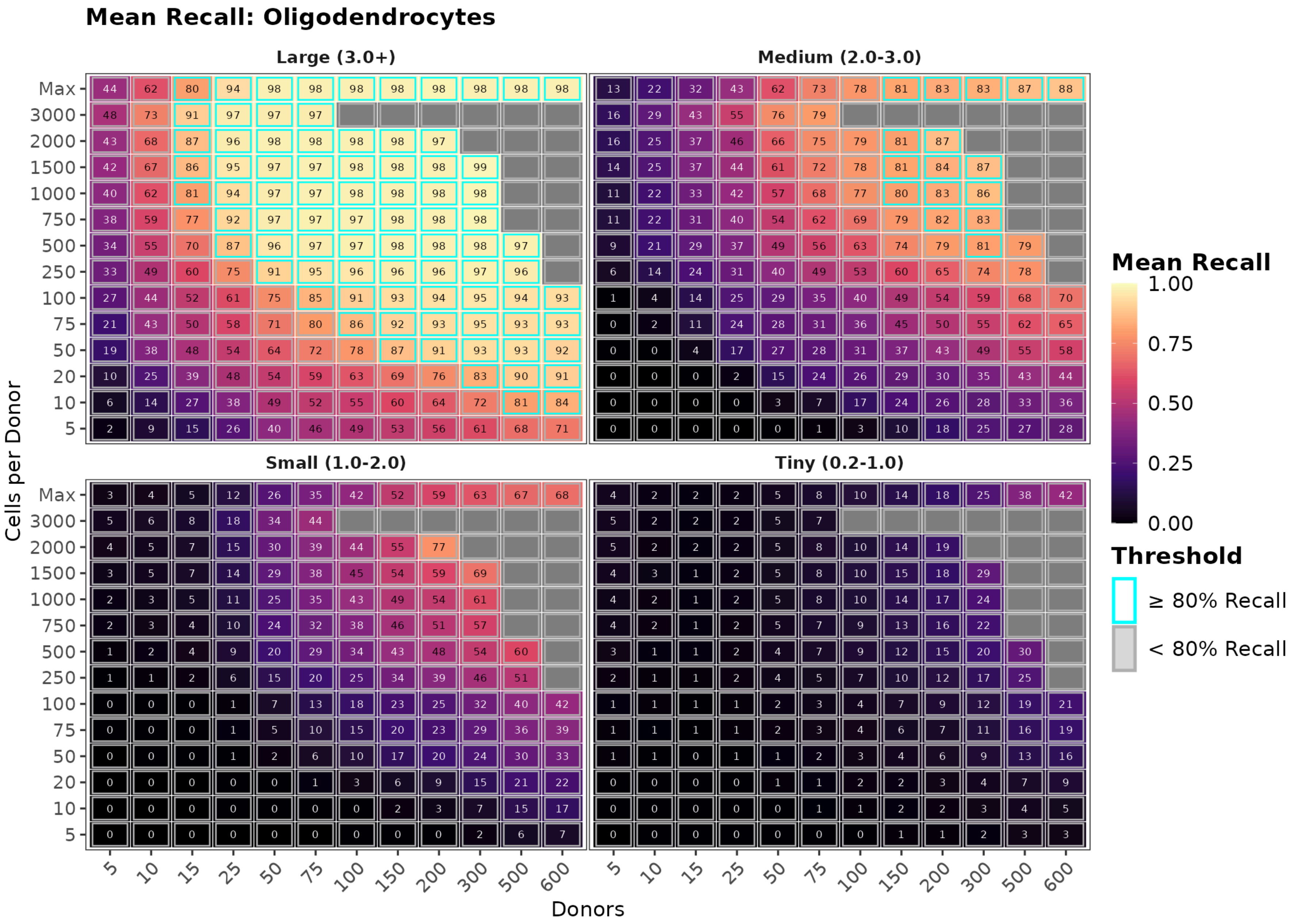


Figure 29: Discrete recall heatmaps from Cartesian downsampling (cells-based mode) for astrocytes (top), microglia (middle), and oligodendrocytes (bottom).

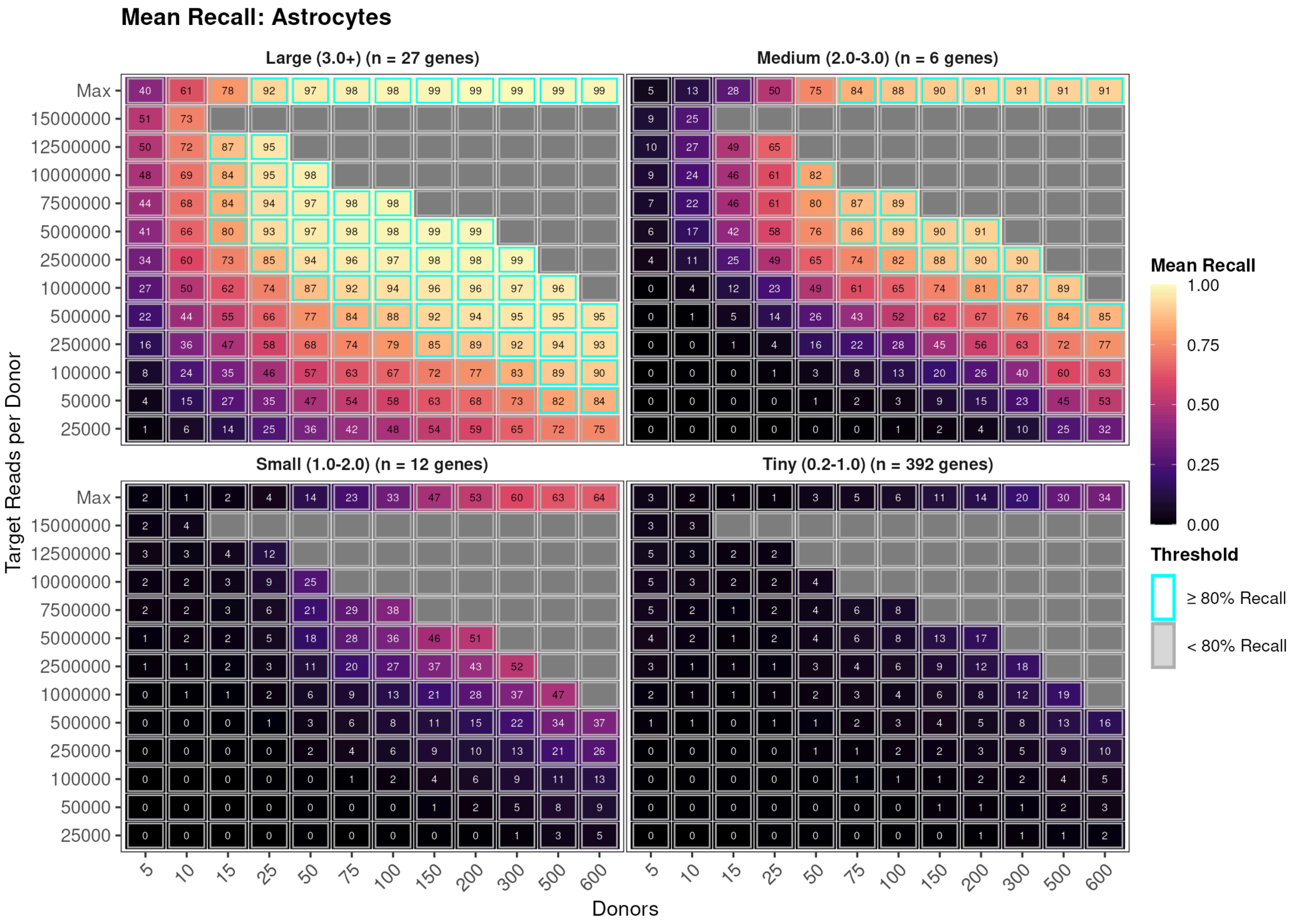
Mean Recall: Astrocytes
Large (3.0+) (n = 27 genes)
Medium (2.0-3.0) (n = 6 genes)
Small (1.0-2.0) (n = 12 genes)
Tiny (0.2-1.0) (n = 392 genes)
Target Reads per Donor
Max
15000000
12500000
10000000
7500000
5000000
2500000
1000000
500000
250000
100000
50000
25000
Donors
5
10
15
25
50
75
100
150
200
300
500
600
Mean Recall
1.00
0.75
0.50
0.25
0.00
Threshold
≥ 80% Recall
< 80% Recall

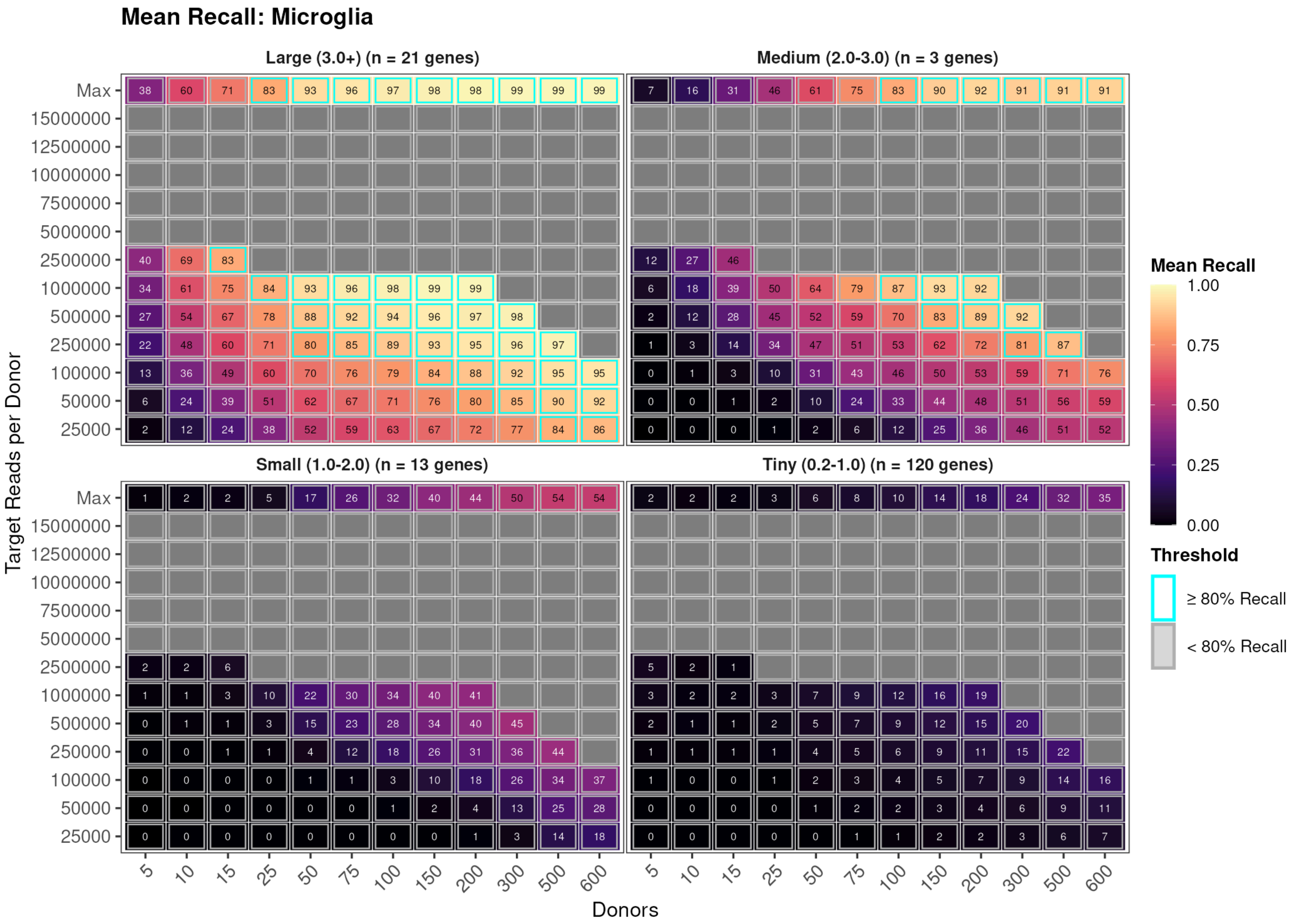
Mean Recall: Microglia
Large (3.0+) (n = 21 genes)
Medium (2.0-3.0) (n = 3 genes)
Small (1.0-2.0) (n = 13 genes)
Tiny (0.2-1.0) (n = 120 genes)
Target Reads per Donor
Max
15000000
12500000
10000000
7500000
5000000
2500000
1000000
500000
250000
100000
50000
25000
Donors
5
10
15
25
50
75
100
150
200
300
500
600
Mean Recall
1.00
0.75
0.50
0.25
0.00
Threshold
≥ 80% Recall
< 80% Recall

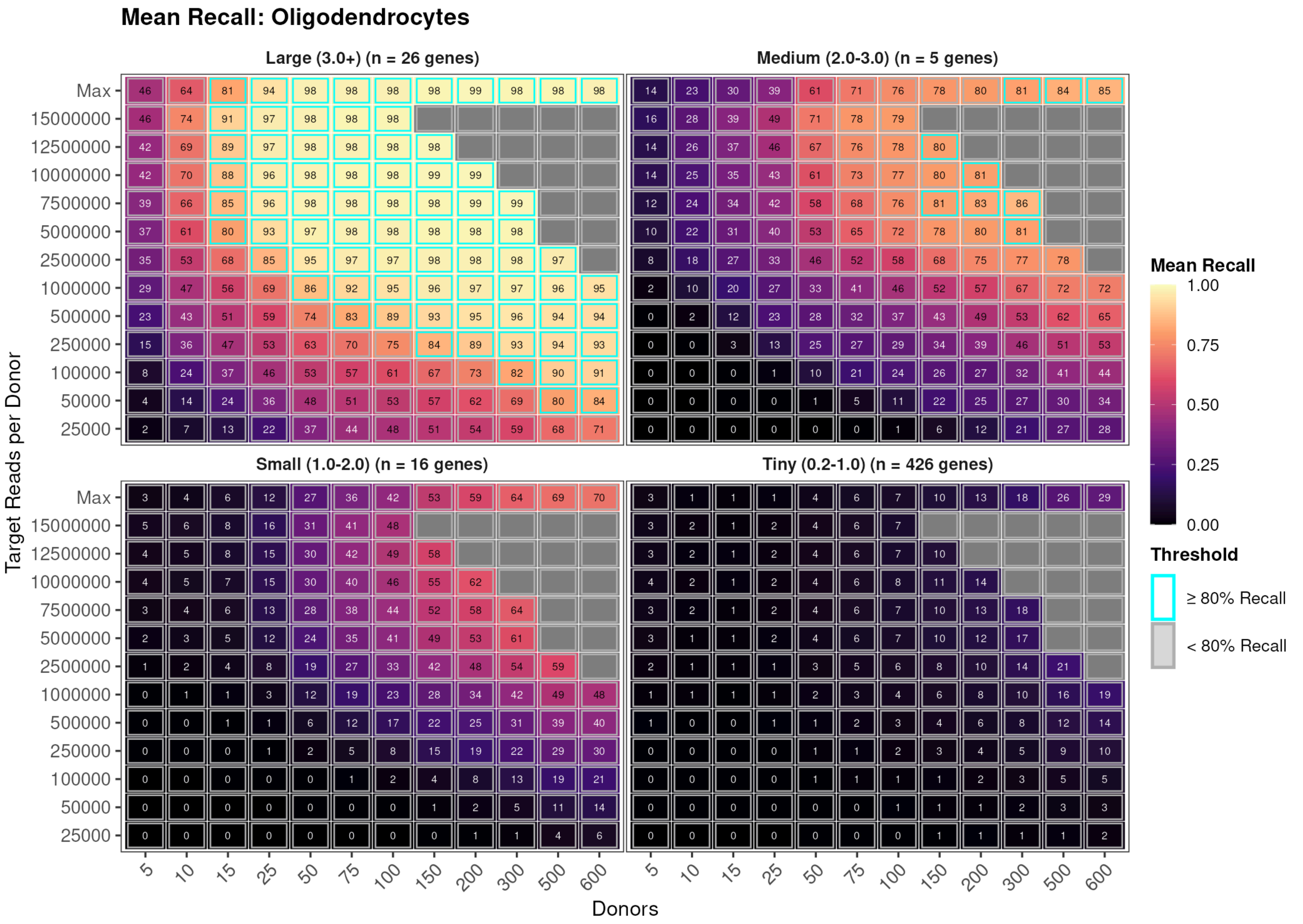


Figure 30: Discrete recall heatmaps from Cartesian downsampling (counts-based mode) for astrocytes (top), microglia (middle), and oligodendrocytes (bottom).

### Supplementary Figures: power-model diagnostics and additional analyses

#### Supplementary Fig. 27. SHAP dependence of detection on design covariates

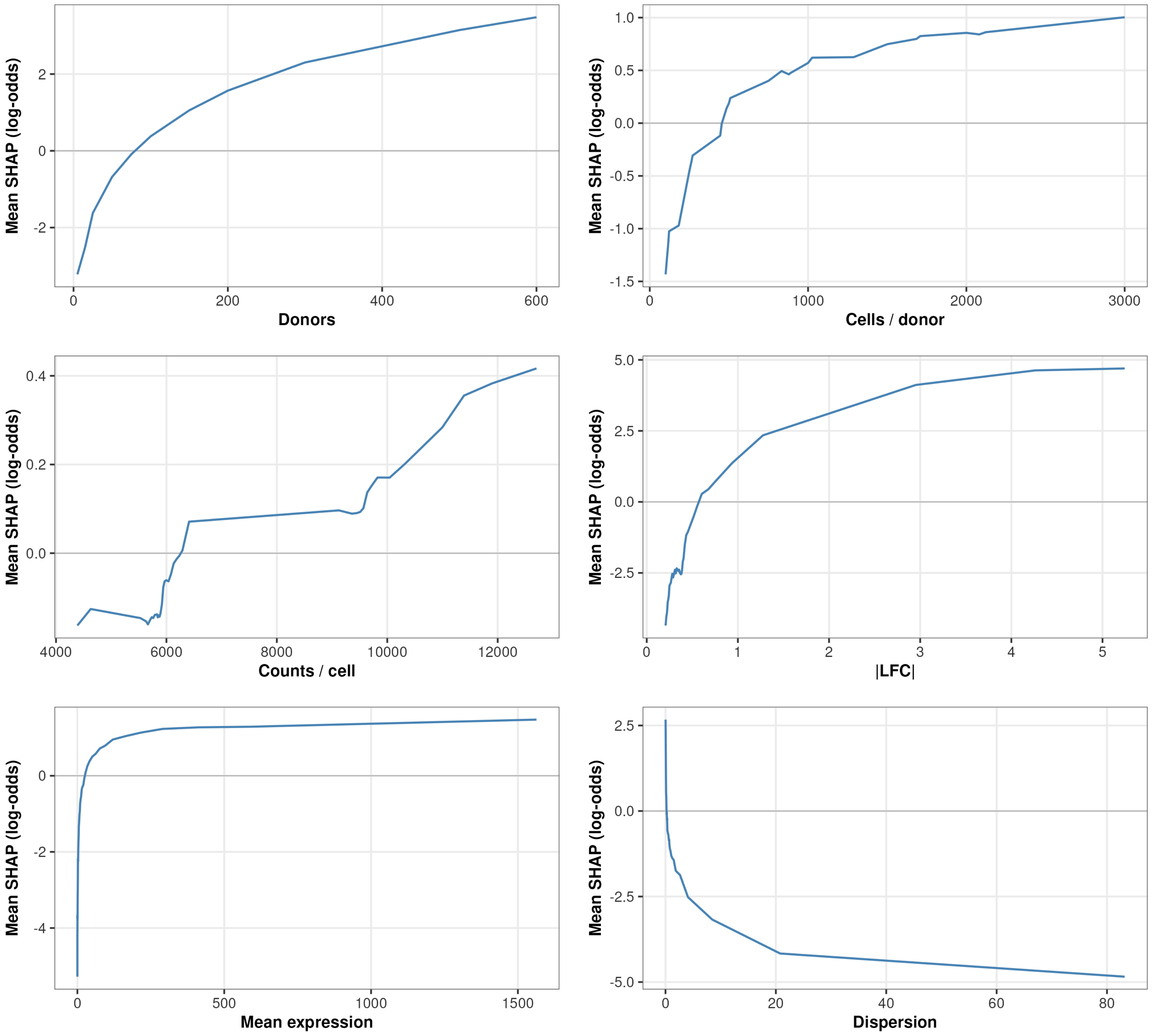


Figure 31: SHAP dependence plots for the production gradient-boosted-tree (XGBoost) power model (realistic $n_{cells} \geq 100$ sub-grid). Each panel plots a design covariate (cells per donor, counts per cell, donor count, effect size, mean expression, dispersion) against its SHAP contribution to predicted detection, with points coloured by covariate value. The steeper SHAP response to cells per donor than to counts per cell reproduces, in the boosted model, the diminishing returns of deeper sequencing relative to broader cellular sampling. Replaces the archived logistic depth-decomposition figure.

### Supplementary Fig. 28. Chromosome-stratified recall heatmaps

Recall heatmaps stratified by chromosome type and LFC magnitude bin. Rows are cell types (astrocytes, microglia, oligodendrocytes); columns are sex-chromosome genes (left) and autosomal genes (right). Autosomal genes show substantially lower recall than sex-chromosome genes at matched effect sizes, reflecting both their rarity and smaller effect sizes. See Supplementary Table 2 for the per-gene characteristics that underlie this difference.

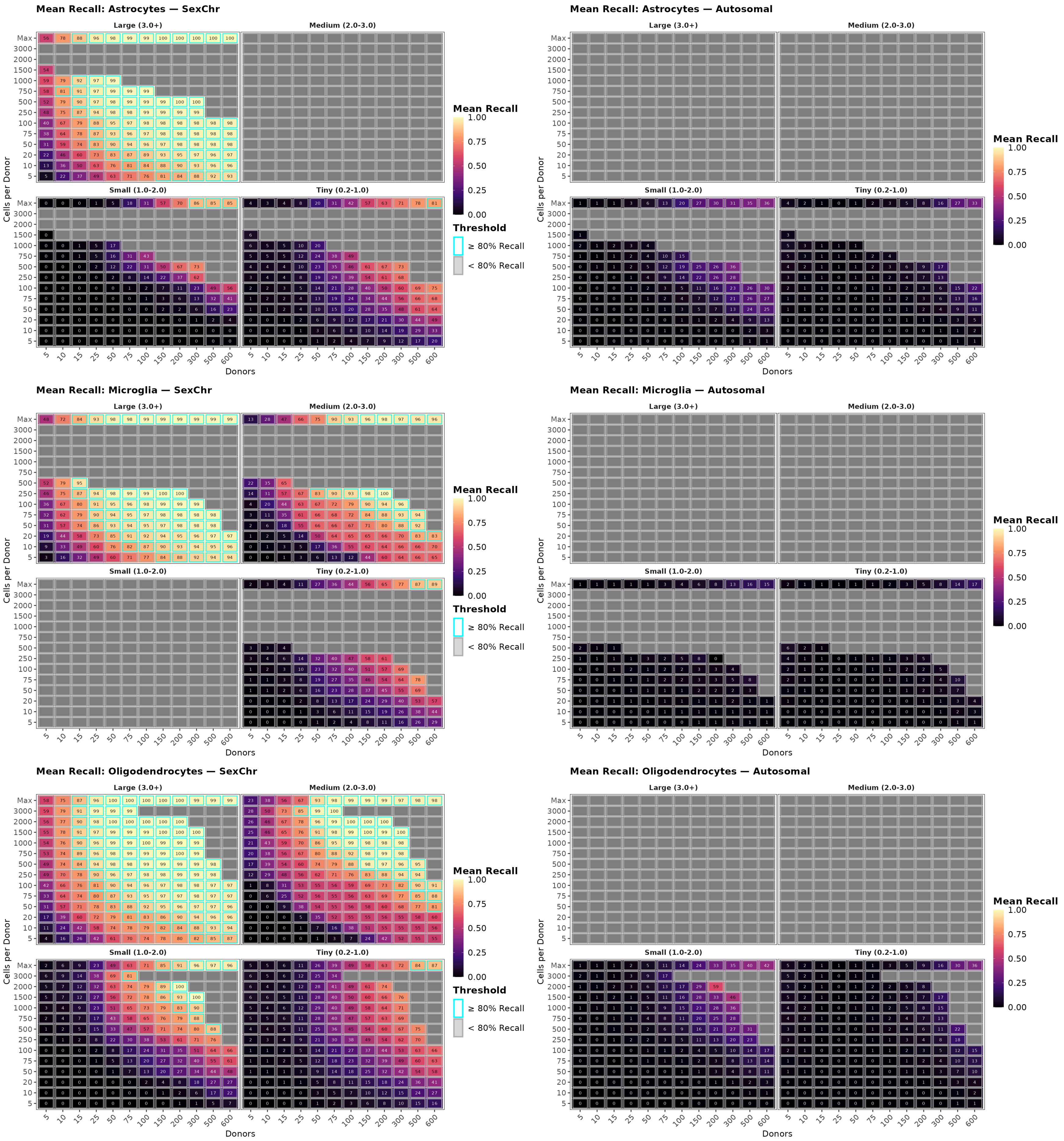


Figure 32: Chromosome-stratified recall heatmaps (cells-based mode). Rows are cell types (astrocytes, microglia, oligodendrocytes); columns are sex-chromosome genes (left) and autosomal genes (right). Sex-chromosome genes dominate the large-LFC bin and are detected with high recall at moderate sample sizes, driving the overall pattern in the total heatmaps. Autosomal sex-biased genes do not show large effect sizes but contribute the majority of tiny-effect genes, with substantially lower recall owing to larger per-gene dispersion and lower mean expression, as explained in the main text.

### Supplementary Fig. 29. DEG chromosome distribution

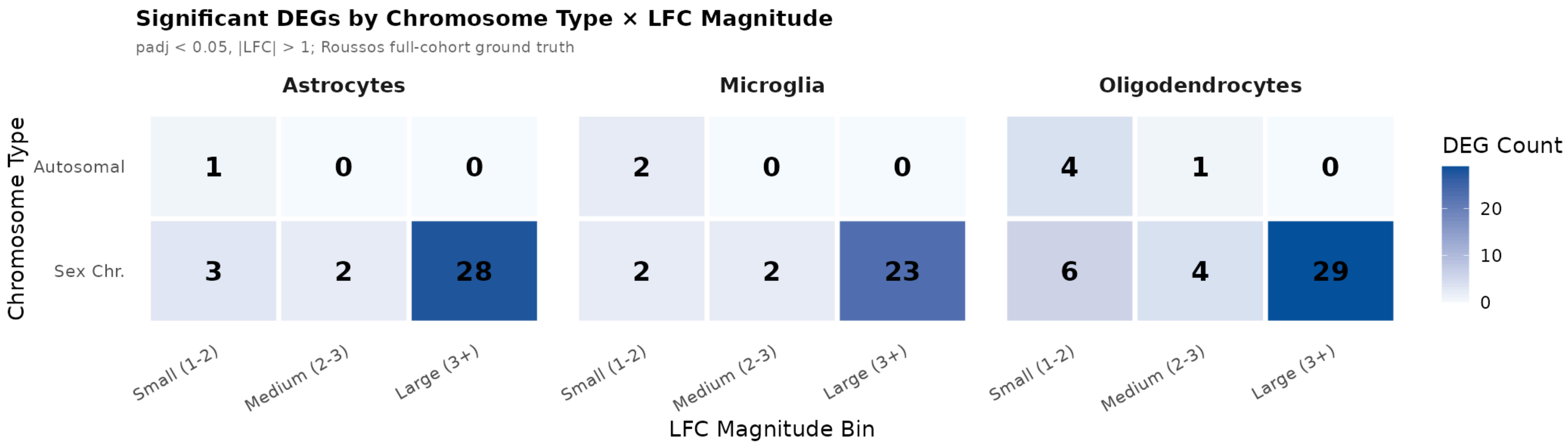


Figure 33: Chromosomal origin of sex-biased DEGs from the PsychAD full ground truth, stratified by |LFC| magnitude and chromosome type (autosomal, X, Y). Top row: counts by chromosome type and LFC bin (1.0-2.0, 2.0-3.0, >3.0) for astrocytes (left), microglia (centre), and oligodendrocytes (right). Bottom row: corresponding proportions. At |LFC| > 3.0, DEGs are predominantly Y-linked; at moderate LFC, autosomal contribution increases.

### Supplementary Fig. 30. Distribution of covariate effects across the grid (SHAP violin)

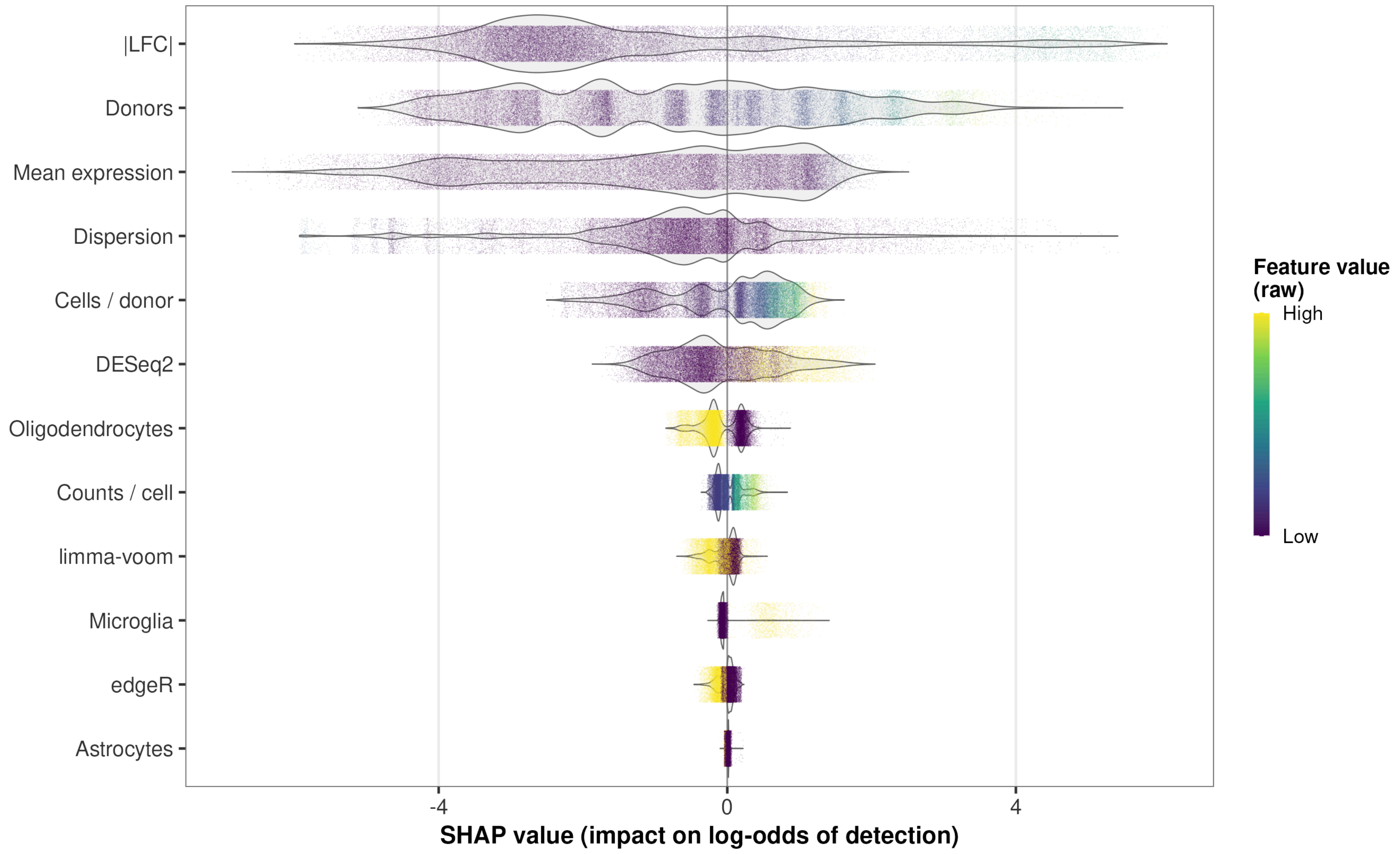


Figure 34: SHAP violin summarising the distribution of each covariate's contribution to predicted detection across the full realistic grid for the production XGBoost model. The spread of each violin is the sensitivity of predicted power to that covariate over its observed range: effect size (|LFC|), donor count, and mean expression carry the widest, most skewed distributions, whereas counts per cell, cell type, and DE method are tightly concentrated near zero. Replaces the archived logistic covariate sensitivity grid.

### Supplementary Fig. 31. SHAP dependence across loss-reweighting configurations

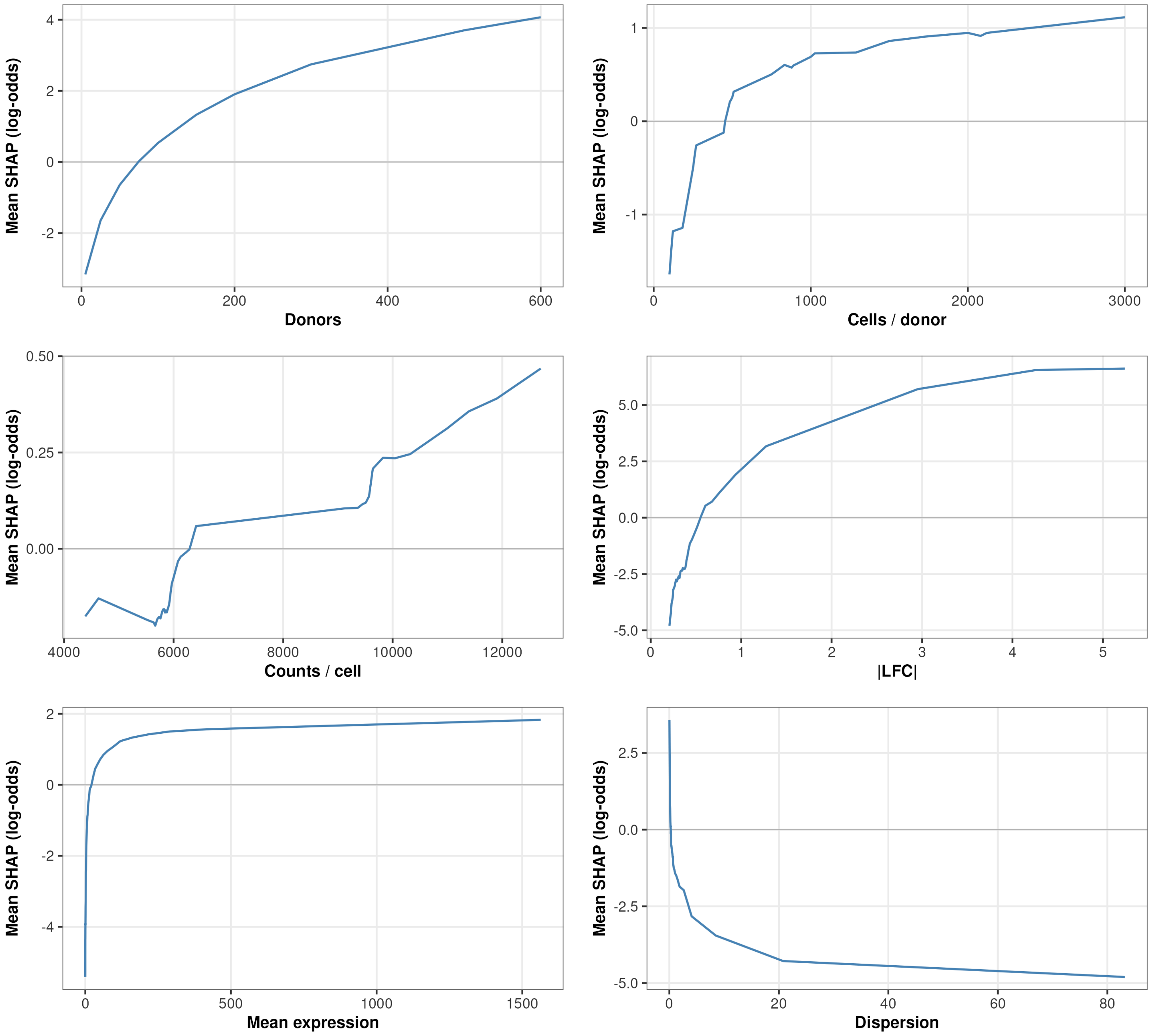
Mean SHAP (log-odds)
Donors
Cells / donor
Counts / cell
|LFC|
Mean expression
Dispersion

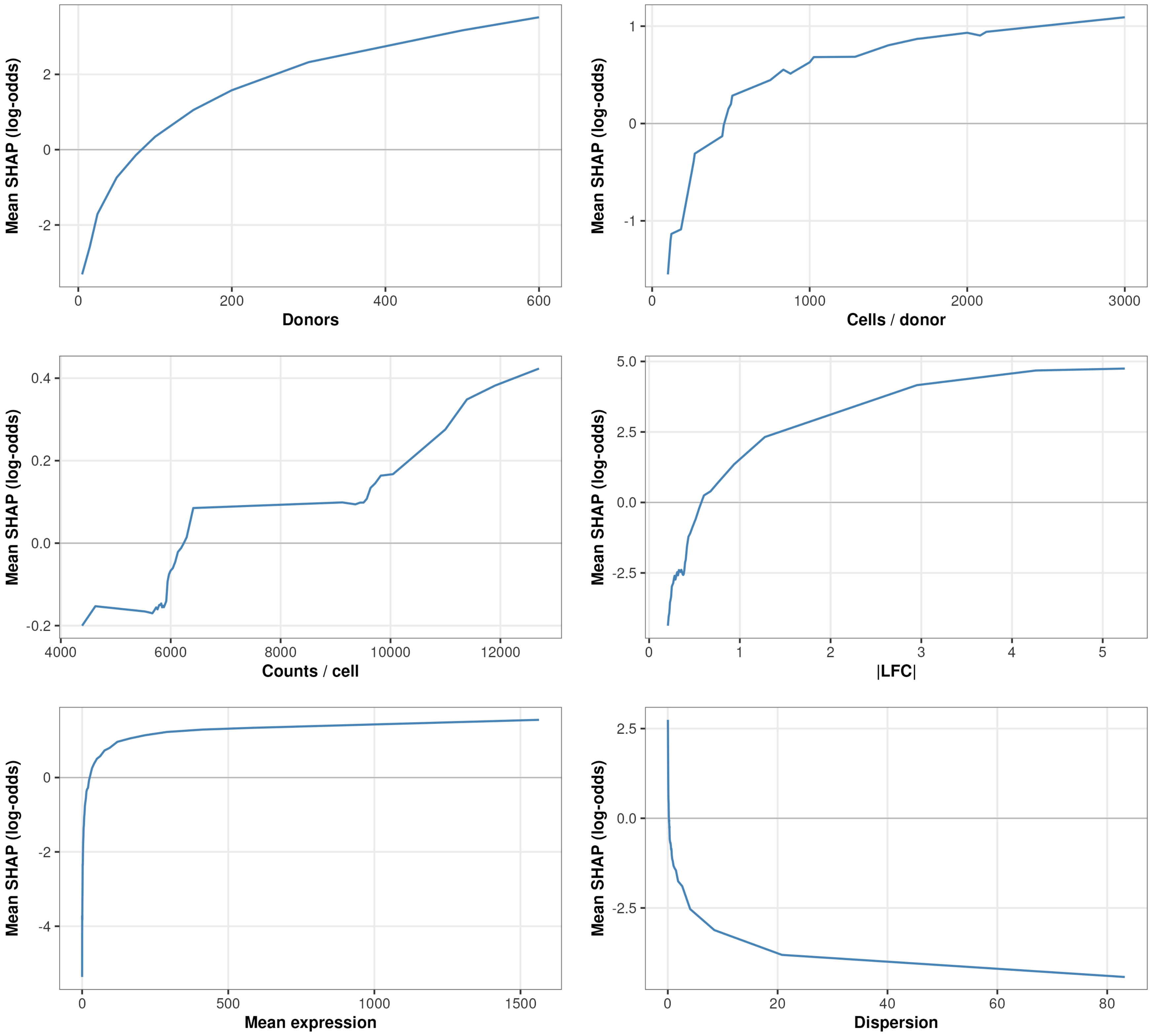

Mean SHAP (log-odds)
Donors
Cells / donor
Counts / cell
|LFC|
Mean expression
Dispersion

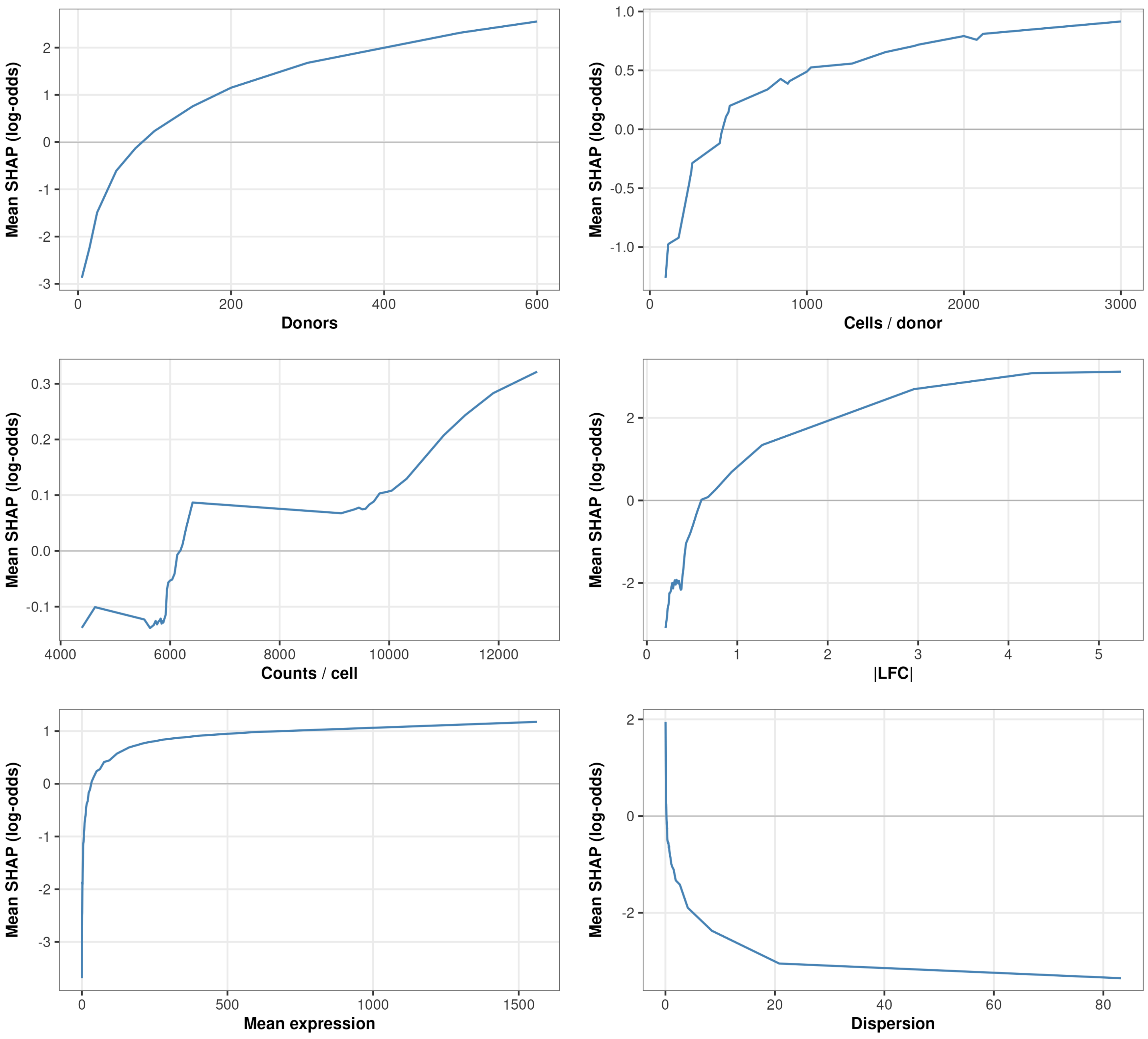


Figure 35: SHAP dependence plots for the three loss-reweighting configurations of the XGBoost power model: unweighted ($\alpha = 0$; top, the promoted production model), and reweighted ($\alpha = 0.5$, middle; $\alpha = 1$, bottom; per-row weight $w = 1/|B|^{\alpha}$ over observed-power decile bins $B$). Each row shows the same six design covariates against their SHAP contribution to predicted detection. The dependence shapes are stable across reweighting exponents, so the covariate-response relationships the model learns are not artefacts of the loss weighting; the unweighted configuration was promoted on out-of-fold error.

### Supplementary Tables

#### Supplementary Table 1. Human brain sc/snRNA-seq DE studies surveyed (2022-2026)

Table 1: Human brain sc/snRNA-seq differential expression studies surveyed (2019-2026). Ordered by donor count (ascending). LFC thresholds are $|\log_2 \mathrm{FC}|$ unless otherwise noted. 'FC' denotes untransformed fold change; 'ln' denotes natural-log fold change; 'coeff' denotes a MAST regression coefficient.

| # | Study | Year | Donors | LFC threshold |
|---|---|---|---|---|
| 1 | Soreq et al.[1] | 2023 | 4 | FC > 1.2 |
| 2 | Puvogel et al.[2] | 2024 | 8 | 0.1 / 0.25 |
| 3 | Bøstrand et al.[3] | 2024 | 12 | 0.25 / 0.8 |
| 4 | Belchikov et al.[4] | 2025 | 12 | 0.1 |
| 5 | Chia et al.[5] | 2025 | 20 | 1.0 / none |
| 6 | Christodoulou et al.[6] | 2023 | 23 | 1.0 |
| 7 | Gamache et al.[7] | 2023 | 24 | 0.2 |
| 8 | Almeida et al.[8] | 2024 | 27 | none / 0.25 |
| 9 | Martirosyan et al.[9] | 2024 | 29 | 0.3 |
| 10 | DeCasien et al.[decasien2026?] | 2026 | 30 | not stated |

| # | Study | Year | Donors | LFC threshold |
|---|---|---|---|---|
| 11 | Li et al.[10] | 2025 | 31 | 0.25 |
| 12 | Nishioka et al.[11] | 2026 | 41 | 0.14-0.26 |
| 13 | Sayed et al.[12] | 2021 | 46 | 0.5 |
| 14 | Murphy et al.[13] | 2023 | 48 | none |
| 15 | Mathys et al.[14] | 2019 | 48 | 0.5 |
| 16 | Yu et al.[15] | 2024 | 53 | 0.25 |
| 17 | Soelter et al.[16] | 2024 | 56 | 0.2 / 0.5 |
| 18 | Barba-Reyes et al.[17] | 2025 | 63 | 0.5 (ln) |
| 19 | Maitra et al.[18] | 2023 | 71 | 0.137 |
| 20 | Johansen et al.[19] | 2023 | ~75 | not found |
| 21 | Miyoshi et al.[20] | 2024 | ~75 | 0.25 |
| 22 | Arbabi et al.[21] | 2025 | 76 | not found |
| 23 | Clarence et al.[22] | 2025 | 97 | 0.2 |
| 24 | Ruzicka et al.[23] | 2024 | 140 | 0.1 |
| 25 | Hoffman et al.[24] | 2024 | 299 | not found |
| 26 | Green et al.[25] | 2024 | 424 | not found |
| 27 | Mathys et al.[26] | 2024 | 427 | not found |
| 28 | Sun et al.[27] | 2023 | 443 | 0.58 |
| 29 | Tang et al.[28] | 2025 | 748 | 0.2 |
| 30 | Lee et al.[29] | 2024 | 1,494 | not found |

### Supplementary Table 2. Autosomal versus sex-chromosome truth-gene characteristics

Table 2: Autosomal versus sex-chromosome gene properties in the PsychAD truth set, stratified by cell type and |LFC| bin. Within each bin, sex-chromosome genes show systematically higher baseMean and lower dispersion than autosomal genes, placing them in a favourable region of the power surface at equivalent fold change. Rows with n < 5 are suppressed.

| Cell Type | \|LFC\| Bin | Group | N | Median BaseMean | Median Dispersion | Median \|LFC\| |
|---|---|---|---|---|---|---|
| Astro | Large (3+) | Sex chr | 28 | 42.7 | 0.230 | 4.92 |
| Astro | Tiny (0.2-1) | Autosomal | 332 | 9.7 | 0.462 | 0.26 |
| Astro | Tiny (0.2-1) | Sex chr | 42 | 62.4 | 0.100 | 0.31 |
| Micro | Large (3+) | Sex chr | 23 | 18.0 | 0.238 | 4.57 |
| Micro | Tiny (0.2-1) | Autosomal | 179 | 5.1 | 0.591 | 0.29 |
| Micro | Tiny (0.2-1) | Sex chr | 28 | 27.9 | 0.073 | 0.40 |
| Oligo | Large (3+) | Sex chr | 29 | 34.4 | 0.302 | 4.43 |
| Oligo | Small (1-2) | Sex chr | 6 | 11.8 | 0.301 | 1.30 |
| Oligo | Tiny (0.2-1) | Autosomal | 321 | 8.1 | 0.515 | 0.27 |
| Oligo | Tiny (0.2-1) | Sex chr | 60 | 44.7 | 0.135 | 0.34 |

### Supplementary Table 3. DEG loss under depth and cell-count equalisation, across DE methods and |LFC| thresholds

Mean number of ground-truth sex-biased DEGs recovered in the PsychAD astrocyte truth set under a 2x2 grid of sequencing-depth and cell-count equalisation, computed independently for each of the three DE methods at two effect-size thresholds. Each cell reports the mean DEG count over 10 robustness runs (50% donor split) with, in parentheses, that count as a percentage of the native (full depth, full cells) count for the same method and threshold. "Depth-equalised" multinomially downsamples every astrocyte to microglia-equivalent counts per cell (factor 0.4679); "Cell-count-equalised" randomly subsamples each donor's astrocytes to microglia-equivalent cell counts at native depth; "Depth + cell equalised" applies both. At the strict |LFC| > 1 threshold all three methods behave consistently, losing DEGs monotonically as cells and depth are reduced. At the permissive |LFC| > 0.2 threshold DESeq2 and edgeR remain well behaved (roughly 55% of DEGs lost under full equalisation), but limma-voom becomes numerically unstable under cell downsampling, inflating its DEG count to several times the native value (327% and 525% of native). The contrast between the two thresholds is the reason limma-voom is excluded from the permissive-threshold analysis.

Table 3: Mean ground-truth DEG count (percentage of native count) under depth and cell-count equalisation, by DE method and |LFC| threshold. limma-voom is numerically unstable at |LFC| > 0.2 under cell downsampling.

| Configuration | DESeq2 (\|LFC\| > 0.2) | DESeq2 (\|LFC\| > 1) | edgeR (\|LFC\| > 0.2) | edgeR (\|LFC\| > 1) | limma (\|LFC\| > 0.2) | limma (\|LFC\| > 1) |
|---|---|---|---|---|---|---|
| Native (full depth & cells) | 444 (100.0%) | 39 (100.0%) | 574 (100.0%) | 57 (100.0%) | 291 (100.0%) | 35 (100.0%) |
| Depth-equalised | 315 (71.0%) | 34 (85.9%) | 409 (71.3%) | 52 (90.1%) | 214 (73.3%) | 31 (89.7%) |
| Cell-count-equalised | 308 (69.5%) | 35 (89.7%) | 393 (68.5%) | 49 (84.7%) | 952 (326.7%) | 32 (90.3%) |
| Depth + cell equalised | 197 (44.5%) | 32 (82.6%) | 258 (45.0%) | 44 (75.8%) | 1528 (524.5%) | 29 (82.9%) |

### Supplementary Table 4. Sex label validation by chromosomal marker expression

Sex labels for all donors were validated by comparing annotated sex against pseudobulk expression of *XIST* (X-inactivation transcript, expected in females) and two Y-chromosome markers (*DDX3Y*, *KDM5D*) using logistic regression on library-size-normalised expression (counts per million, CPM). Expression was aggregated to the donor level before modelling to avoid single-cell noise[30]. Four donors (0.3% of the cohort) showed strong discordance between their assigned sex and chromosomal marker expression; their statistics are shown below. All flagged donors were retained in the main analyses, as the ambiguity speaks to population-level variability, not a labeling error.

Table 4: Donors with discordant sex labels identified by pseudobulk chromosomal marker analysis. XIST CPM = library-size-normalised expression of XIST (ENSG00000229807); Y-gene CPM = combined normalised expression of DDX3Y (ENSG00000067048) and KDM5D (ENSG00000012817). P(female|XIST) and P(male|Y) are logistic regression probabilities fitted on all donors in each cell type. n_cells = number of nuclei for that donor in that cell type.

| Donor ID | Cell Type | Labelled Sex | Discordance Flag | XIST CPM | P(female\|XIST) | Y-gene CPM | P(male\|Y) | N Cells |
|---|---|---|---|---|---|---|---|---|
| Donor_1296 | Astro | Male | XIST in labelled male | 1071.2 | 0.999 | 71.1 | 0.999 | 1004 |
| Donor_1331 | Astro | Female | Y gene in labelled female | 450.1 | 0.920 | 62.0 | 0.996 | 159 |
| Donor_1296 | Micro | Male | XIST in labelled male | 1149.9 | 0.999 | 79.8 | 0.998 | 371 |
| Donor_484 | Micro | Female | Y gene in labelled female | 1515.4 | 1.000 | 44.7 | 0.874 | 92 |
| Donor_695 | Micro | Male | Y gene absent in labelled male | 124.8 | 0.001 | 7.8 | 0.000 | 138 |
| Donor_1296 | Oligo | Male | XIST in labelled male | 1475.3 | 0.999 | 96.3 | 0.997 | 3625 |
| Donor_1331 | Oligo | Female | Y gene in labelled female | 790.1 | 0.985 | 85.4 | 0.992 | 351 |
